# From single-sequence structure prediction to protein fitness landscape through a composable, epistasis-aware mutation atlas

**DOI:** 10.64898/2026.09.24.753701

**Authors:** Weizhe Wang, Zimu Yu, Endi Yang, Ziyu Shi, Shize Yu, Jian Hu, Yunxin Xu, Haipeng Gong

**Author notes:** These authors contributed equally (Weizhe Wang, Zimu Yu).

## Abstract

Mapping the multi-mutant fitness landscape is vital to protein engineering, but is challenging due to the vast combinatorial sequence space awaiting exploration. A central difficulty lies in the accurate and efficient modeling of non-additive epistatic effects among individual mutations, which partially arise from the physical inter-residue interactions prescribed by the protein structure. Existing fitness predictors usually perform well on single mutants but become less powerful for higher-order mutants, due to the lack of explicitly considering the relationship between sequence, structure and function of the target protein. Here, we present an end-to-end framework named Cerebra-Epistasis, which couples a single-sequence structure predictor that explicitly endows the structure awareness beyond the conventional sequence-fitness mapping with a downstream fitness prediction network that deliberately models the non-linear epistatic effects beyond the traditional additive terms. When evaluated across diverse assays, Cerebra-Epistasis outperforms the other state-of-the-art baselines in multi-mutant fitness prediction, with enhanced advantage over increasing mutation orders. Moreover, our special design on the epistasis modeling allows reliable extrapolation from low-order mutant data to unseen higher-order combinations, enabling one-shot inference of the overall mutation atlas from the starting sequence, a benefit that supposedly introduces three orders of magnitude acceleration in the landscape-scale prediction.

## Introduction

Engineering protein function by sequence variation has broad biotechnological impacts, and has been extensively applied in practical fields including enzyme engineering and therapeutic development. Experimental methods like deep mutational scanning (DMS) can only capture a tiny fraction of the full sequence combinations from a wild-type protein [1, 2]. This deficit leaves the large-scale unmeasured mutational spectrum to be explored and predicted by computational methods. To guide protein optimization, a computational model is supposed to accurately map the multi-mutant fitness landscape, particularly the epistatic effects between individual mutations, while supporting efficient inference at the landscape scale.

Early evolutionary models infer mutational effects from the multiple sequence alignment (MSA) using sitewise conservation or pairwise sequence covariation [3]. Subsequently developed protein language models (PLMs) enable zero-shot mutation scoring from representations learned on large-scale sequence data [4, 5]. In these methods, epistatic effects are either completely neglected by the additive processing of sitewise scores or oversimplified as sparse pairwise terms using graphical models. Sparse regression methods, such as LASSO, extend such pairwise decomposition by directly fitting selected low-order interaction terms to measured fitness landscapes [6], but remain powerless in inferring unseen mutants due to the limited model-specified terms. More recent supervised methods begin to learn the nonlinear projection from complete variant sequences to measured fitness. ProteinNPT combines protein representations with available property measurements via a non-parametric transformer, and thus supports prediction in label-scarce and multi-property settings [7]. By combining PLM embeddings with a regression model, EVOLVEpro iteratively updates the model using small batches of experimental measurements to guide protein optimization [8]. MULTI-evolve focuses more directly on higher-order mutant design, by training a nonlinear predictor on strategically measured single- and double-mutant variants and then applying it to higher-order combinations [9].

Despite versatile objectives, these methods directly predict the total fitness, and more importantly, require each candidate variant to be represented and evaluated separately, which leads to the scaling of storage, GPU memory and computation time with the candidate size during the construction of a large-scale fitness landscape. One possible solution is to construct separate reusable representations of single-mutation effects and epistatic effects, respectively, which not only allows instant derivation of the total fitness through direct assembly but also retains the flexibility of results across mutation orders. The epistatic representation should be able to reflect the change of pairwise effects upon the alteration of mutational background, so as to capture the prominent three-way and higher-order epistasis revealed in previous research [6, 10]. Thus, the remaining challenge is to combine accurate fitness prediction, explicit epistasis modeling across varying mutation orders, and efficient landscape-scale inference within a unified framework.

The 3D structure of a protein provides physical context for epistatic interactions: residues that are distant in sequence can be proximal in space, giving rise to coupled effects. PLMs and MSA-based methods capture evolutionary constraints but do not explicitly represent structural information such as local packing, long-range contacts, and/or side-chain geometry. Structure-based inverse-folding scores offer complementary information to sequence-based models, and multimodal approaches that integrate sequence, MSA and structure have been shown to improve mutation-effect prediction across diverse assay types [11, 12]. Building on these efforts, Kermut combines sequence and structural information through a composite Gaussian process kernel for supervised variant-effect prediction [13]. SPIRED-Fitness moves further by coupling a single-sequence structure predictor to a fitness model and training both components end to end [14]. Together, by demonstrating that structural context can inform mutation-effect prediction, these studies motivate the integration of structure awareness with explicit epistasis modeling. However, realizing such an integration scheme within an end-to-end fitness framework requires a fast, reliable structure predictor that supports end-to-end optimization at an acceptable computational cost.

MSA-based structure predictors such as AlphaFold 2 require extensive preprocessing and substantial computation [15]. Single-sequence models such as ESMFold and OmegaFold circumvent the MSA-search bottleneck to accelerate inference, but still require computation and memory intensive training [16, 17], which hinders end-to-end training with downstream fitness models. SPIRED-Fitness overcomes this problem by using a lightweight single-sequence predictor SPIRED, which represents structures mainly through C_*α*_ coordinates but has to rely on a separate pipeline for all-atom reconstruction [14]. An ideal structure-aware fitness framework therefore demands a structure predictor that combines high structural accuracy, single-sequence efficiency, side-chain-resolved representations, and sufficiently low computational cost for end-to-end training.

In this study, we present Cerebra-Epistasis, an end-to-end structure-aware framework for efficient construction of the protein fitness landscape. Its structure prediction module, Cerebra-Seq, is an accurate and efficient single-sequence model derived from the original MSA-based predictor Cerebra [18]. Cerebra-Seq produces side-chain-resolved structural representations and supports end-to-end training with the downstream fitness prediction network, a design that provides explicit 3D structure awareness for each target protein. The downstream fitness predictor decomposes multi-mutant fitness into additive single-mutation components and an independent epistatic contribution modeled across various mutation orders. By this means, Cerebra-Epistasis performs sequence and structure encoding only once to construct a reusable mutation-indexed atlas for a target protein. When evaluated across diverse multi-mutant benchmarks, Cerebra-Epistasis consistently achieves the leading performance in predicting both multi-mutant fitness and epistatic effects, with end-to-end training introducing further benefits to fitness prediction. The reusable mutation atlas also provides an interpretable decomposition of multi-mutant fitness and enables efficient evaluation of the large combinatorial fitness landscape.

## Results

### Overview of Cerebra-Epistasis

We propose Cerebra-Epistasis, a structure-aware framework that encodes the wild-type (WT) protein once and reuses the resulting representations to predict arbitrary-order mutant fitness (Fig 1A). The WT encoding is used to construct a mutation atlas covering all 20 amino-acid states at every sequence position. For a given multi-mutant, the corresponding entries are retrieved from the atlas and assembled to predict its fitness value.

**Figure 1.**
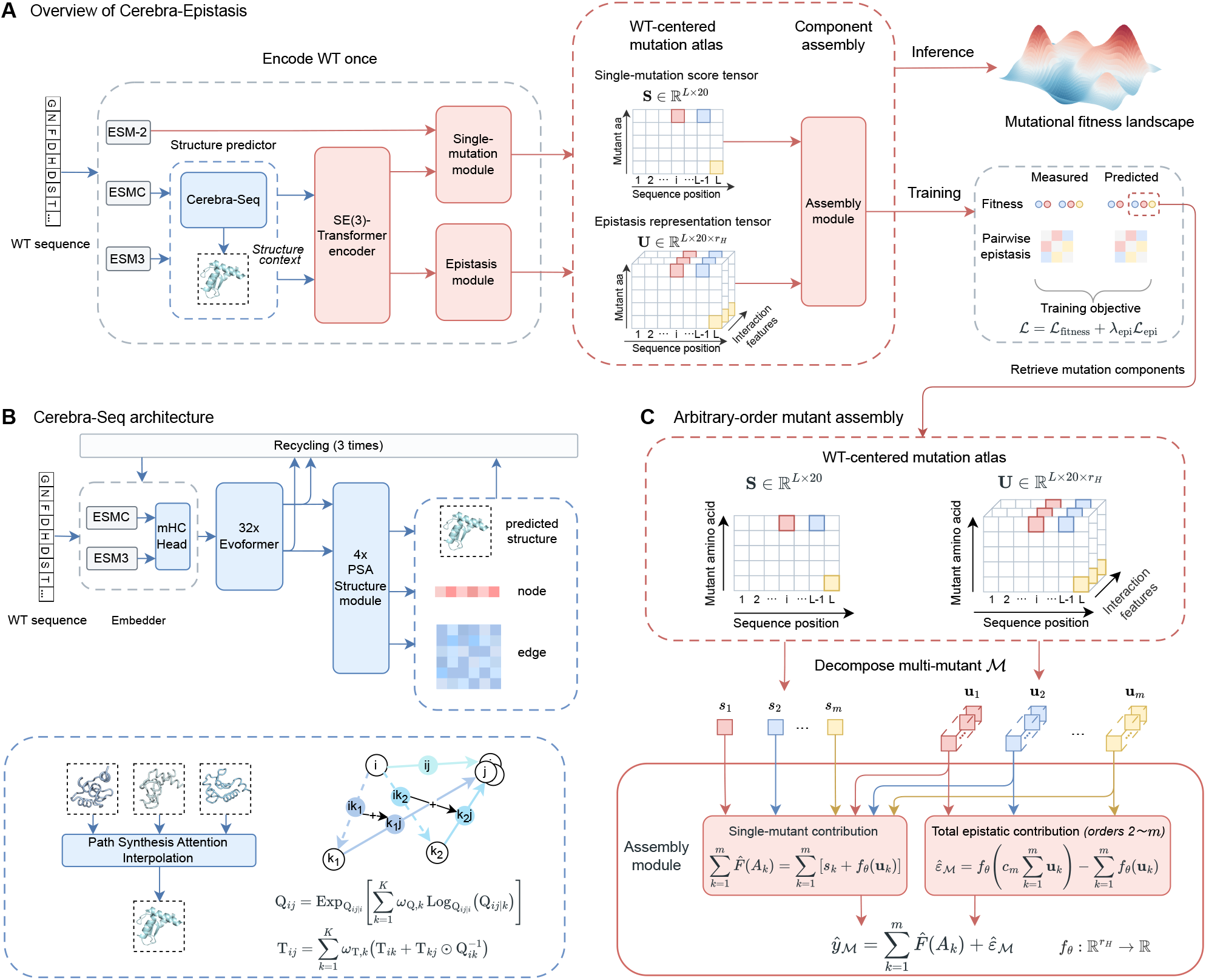
Overview of Cerebra-Epistasis. **A**, The WT protein is encoded once to construct a mutation-indexed atlas comprising the single-mutation score tensor **S** and epistasis representation tensor **U**. Mutation components are then assembled to predict fitness and epistasis across the mutant landscape. The framework supports downstream-only training with Cerebra-Seq frozen and end-to-end training. **B**, Architecture of Cerebra-Seq, which combines ESMC and ESM3 representations through an mHC-based mixing head, Evoformer blocks, and PSA structure modules to generate structural, node, and edge representations. **C**, Arbitrary-order mutant assembly through direct retrieval and permutation-invariant combination of the corresponding components from **S** and **U**, enabling lightweight fitness-landscape prediction without repeated sequence or structure encoding.

The structure awareness is accomplished through Cerebra-Seq, a single-sequence structure predictor that combines ESMC [19] and ESM3 [20] representations through an mHC-based mixing head [21], followed by Evoformer blocks and PSA [18] structure modules for information processing (Fig 1B). It predicts atomic-level protein structures and simultaneously generates residue-level node and residue-pair edge representations, providing structural context for downstream fitness prediction.

In the downstream fitness model, an SE(3)-equivariant network integrates the coordinates, node representations, and edge representations generated by Cerebra-Seq to capture the 3D geometry and residue-pair context, as well as ESM-2 [16] embedding to facilitate the residue-level representation. These features are transformed into a single-mutation score tensor **S** and an epistasis representation tensor **U** (Fig 1C), where **S** scores individual substitutions and **U** provides representations for modeling epistatic interactions. This design explicitly separates single-mutation contributions from the joint epistatic terms, effectively disentangling the components of multi-mutant fitness. The corresponding atlas entries can be assembled for arbitrary mutant combinations, enabling evaluation of the large combinatorial fitness landscape without repeated sequence or structure encoding.

### Cerebra-Seq achieves accurate single-sequence structure prediction

We evaluated Cerebra-Seq on two CAMEO [22] benchmarks to assess whether its single-sequence structure-prediction performance was reproducible across independent test periods. The cleaned CAMEO 2024 and CAMEO 2025 sets contain 642 and 573 protein targets released in the preceding two years, respectively. Cerebra-Seq was compared with ESMFold [16], OmegaFold [17], direct structure generation by ESM3 [20], and SPIRED [14]. Within each benchmark, all summary statistics and paired comparisons were calculated on the targets shared by all five methods. Prediction accuracy was evaluated using TM-score, which measures global fold similarity.

On CAMEO 2024, Cerebra-Seq achieved a mean TM-score of 0.83, second only to ESMFold at 0.86 (Fig 2A), presenting a mean paired advantage of 0.01, 0.04, and 0.05 over OmegaFold, ESM3, and SPIRED, respectively. In terms of the individual proteins, Cerebra-Seq outperformed ESMFold on 255 out of the 642 shared targets, which confirms its complementarity to this state-of-the-art baseline.

**Figure 2.**
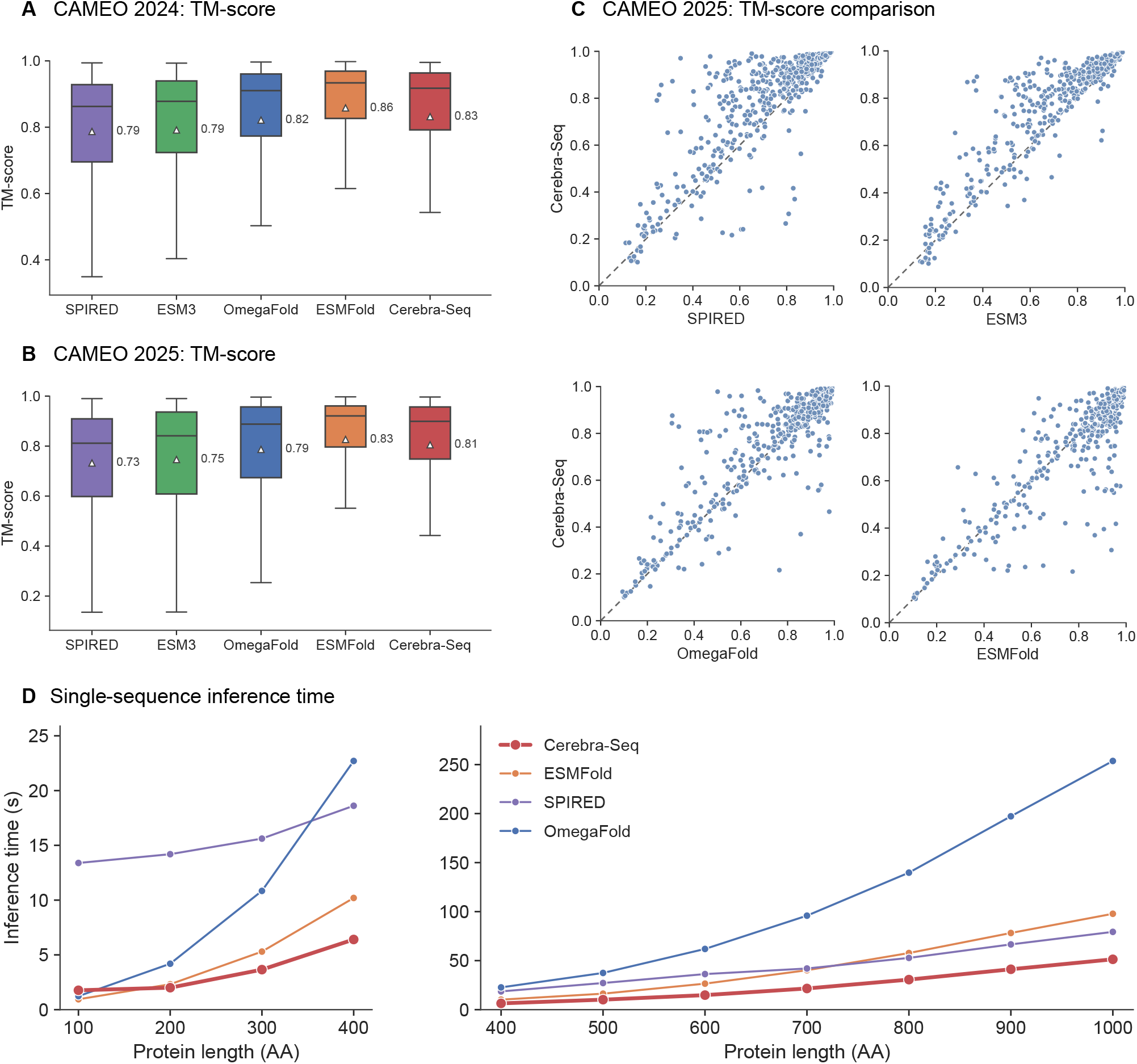
Accuracy and inference efficiency of Cerebra-Seq for single-sequence structure prediction. **A**,**B**, Per-target TM-score distributions for Cerebra-Seq and the baseline methods on the CAMEO 2024 (**A**) and CAMEO 2025 (**B**) benchmarks. White triangles and adjacent values indicate mean TM-scores; center lines denote medians, boxes represent the interquartile range, and whiskers extend to 1.5 times the interquartile range. **C**, Paired per-target comparison of Cerebra-Seq with SPIRED, direct ESM3 structure generation, OmegaFold, and ESMFold on CAMEO 2025. Each point denotes a target shared by all five methods. The diagonal line marks equal TM-scores; points above the line favor Cerebra-Seq. **D**, Single-sequence inference times of Cerebra-Seq, ESMFold, SPIRED, and OmegaFold for proteins ranging from 100 to 1,000 residues. The left and right panels show the 100–400- and 400–1,000-residue ranges, respectively.

On CAMEO 2025, Cerebra-Seq again ranked second by the mean TM-score, reaching 0.81 compared with 0.83 for ESMFold (Fig 2B). Its advantage over OmegaFold, ESM3, and SPIRED was 0.02, 0.06, and 0.07, respectively (Fig 2C). At the individual-protein level, Cerebra-Seq exceeded ESMFold on 225 out of the 573 shared targets.

Across both benchmarks, Cerebra-Seq consistently outperformed direct ESM3 structure generation. This improvement suggests that integrating ESM3 with ESMC and subsequent structural refinement provides additional information for structure prediction. Although ESMFold achieved the highest mean TM-score, Cerebra-Seq outperformed it on substantial subsets of targets. The two CAMEO benchmarks show that Cerebra-Seq can predict all-atom protein structures from single sequences with sufficient accuracy.

### Training and inference cost of Cerebra-Seq

The trainable structure-prediction component of Cerebra-Seq contains 77.0M parameters, and final-model training required 572 GPU-days. Using the same structure-component counting convention, the corresponding values reported in SPIRED-Fitness are 125M parameters and 85 GPU-days for SPIRED, 124M and 3,456 GPU-days for OmegaFold, and 690M and approximately 896 GPU-days for ESMFold [14]. Cerebra-Seq therefore has the smallest reported structure-predictor parameter count and a training cost below those of ESMFold and OmegaFold, although GPU-days are not directly comparable across different training protocols and hardware.

We benchmarked Cerebra-Seq against ESMFold, OmegaFold, and SPIRED on sequences of 100–1,000 residues for inference cost (Fig 2D). Albeit slower than ESMFold and OmegaFold at 100 residues (1.77 s vs. 0.95 and 1.23 s), Cerebra-Seq was the fastest predictor at every tested length from 200 residues onward. Across the 200–1,000-residue range, Cerebra-Seq took 20.20 s per target on average, corresponding to 1.84-, 4.53-, and 1.94-fold speedups over ESMFold, OmegaFold, and SPIRED, respectively. For sequences 1,000 residues long, the speedup reached 1.90-, 4.94-, and 1.55-fold, respectively.

These inference benchmarks further show that Cerebra-Seq attains structural accuracy and efficient single-sequence inference simultaneously, supporting its use as the structure-prediction module within Cerebra-Epistasis.

### Cerebra-Epistasis accurately predicts mutant fitness across diverse tasks

We evaluated Cerebra-Epistasis in two benchmark settings grounded in established protein-fitness evaluation protocols (Fig 3A,F). The 1M+ to 1M+ benchmark follows the random cross-validation setting commonly used in ProteinGym [5], with position balancing to ensure comparable mutation-site coverage across folds (Supplementary Section S2.3). The more protein-engineering-relevant Low-*N* 1M/2M to 3M+ benchmark follows the low-data higher-order extrapolation setting used in MULTI-evolve [9], by training on fewer than 300 single- and double-mutant measurements and testing on unseen higher-order combinations (Supplementary Section S2.4).

**Figure 3.**
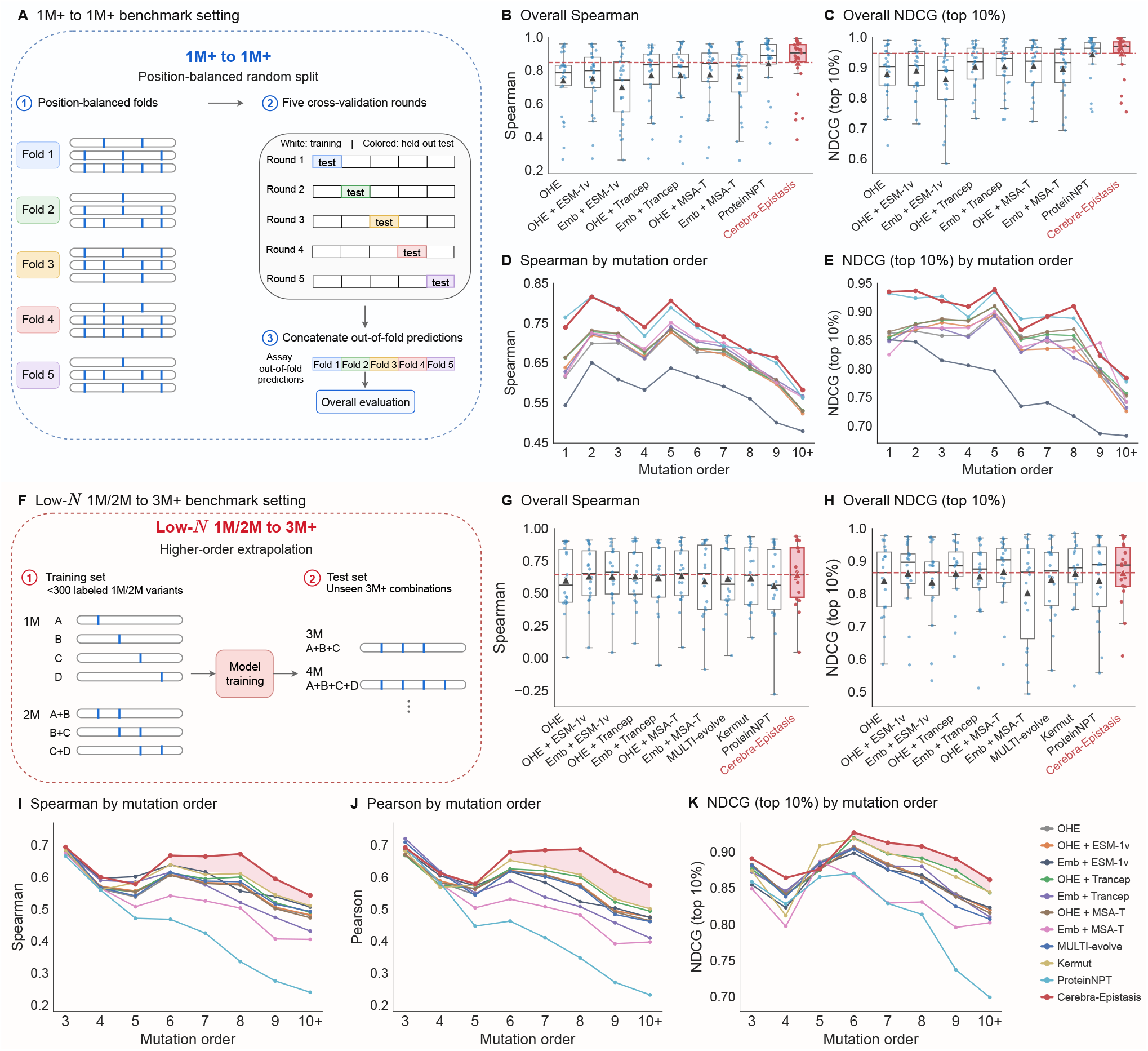
Cerebra-Epistasis accurately predicts mutant fitness across diverse tasks. **A**, Position-balanced 1M+ to 1M+ benchmark setting. **B**,**C**, Overall assay-level Spearman correlation and NDCG (top 10%), respectively; points denote assays, triangles denote assay means, and the red dashed line indicates the Cerebra-Epistasis mean. **D**,**E**, Assay-averaged Spearman correlation and NDCG (top 10%) at each mutation order, respectively. **F**, Low-*N* 1M/2M to 3M+ benchmark setting. **G**,**H**, Overall assay-level Spearman correlation and NDCG (top 10%) in the Low-*N* benchmark, respectively; points denote assays, triangles denote assay means, and the red dashed line indicates the Cerebra-Epistasis mean. **I–K**, Assay-averaged Spearman correlation, Pearson correlation, and NDCG (top 10%) at each mutation order in the Low-*N* benchmark, respectively.

On the 1M+ to 1M+ benchmark, Cerebra-Epistasis achieved the strongest overall assay-level performance (Supplementary Table S5), with statistics of assay-level results shown in Fig 3B,C. Specifically, it achieved a mean Spearman correlation of 0.85 and an NDCG (top 10%) of 0.95, compared with 0.84 and 0.94, respectively, for the second-best baseline ProteinNPT [7]. Notably, Cerebra-Epistasis slightly outperformed this leading supervised multi-mutant fitness predictor despite encoding only the wild-type protein rather than each mutant sequence, indicating that multi-mutant fitness can be effectively predicted within a shared wild-type context by learning how mutation-specific effects combine. This advantage was broadly maintained across mutation orders (Fig 3D,E).

Because practical protein engineering often relies on limited measurements of low-order mutants to identify promising higher-order variants, we next evaluated Cerebra-Epistasis in the Low-*N* 1M/2M to 3M+ benchmark, where it showed a clearer advantage over competing methods (Fig 3G,H; Supplementary Table S6). Specifically, it achieved an overall mean Spearman correlation of 0.65, compared with 0.62 for Kermut, 0.61 for MULTI-evolve, and 0.56 for ProteinNPT. When performance was averaged across mutation orders, Cerebra-Epistasis remained the strongest at 0.65, superior to 0.62 for Kermut, 0.63 for MULTI-evolve, and 0.57 for ProteinNPT. Whereas MULTI-evolve [9] learns epistatic patterns through a nonlinear fitness predictor trained on single- and double-mutant measurements, Cerebra-Epistasis uses a dedicated epistasis representation optimized with both fitness and epistasis supervision. As mutation order increased, all methods showed declining performance, but Cerebra-Epistasis declined more slowly, resulting in a progressively larger advantage at higher orders (Fig 3I-K). This trend supports the value of explicitly modeling epistasis for extrapolating from sparse low-order measurements to unseen higher-order combinations.

We also evaluated Cerebra-Epistasis in three additional benchmark settings. The 1M/2M to 3M+ benchmark further examined the model’s ability to extrapolate from low-order measurements to unseen higher-order variants in the presence of more available data (Supplementary Section S2.5; Supplementary Table S7), while the cDNA proteolysis multi-mutant benchmark evaluated performance across a large collection of independent multi-mutant assays (Supplementary Section S2.6; Supplementary Table S8). We further validated Cerebra-Epistasis on an independent avGFP wet-lab–measured test set (Supplementary Section S2.7; Supplementary Table S9). Across all three additional benchmarks, Cerebra-Epistasis achieved the strongest overall performance among the tested methods.

### Cerebra-Epistasis predicts unseen epistatic interactions and reveals biologically meaningful patterns

Epistasis—the non-additive interactions between mutations—is a major source of complexity in the protein fitness landscape. Unlike many variant-level fitness predictors, in which epistasis remains implicit and must be derived post hoc from predicted mutant fitness values, Cerebra-Epistasis explicitly separates epistatic effects from single-mutation contributions. To assess the contribution of explicit epistasis modeling, we compared Cerebra-Epistasis with two additive formulations, Ridge-additive and Cerebra-Epistasis-additive. Ridge-additive uses *L*_2_-regularized linear regression to model multi-mutant fitness as the sum of single-mutation effects, whereas Cerebra-Epistasis-additive disables the epistasis branch of Cerebra-Epistasis and retains only its single-mutation contributions. On the Low-*N* 1M/2M-to-3M+ benchmark, Cerebra-Epistasis consistently outperformed both additive formulations across mutation orders (Fig 4A).

**Figure 4.**
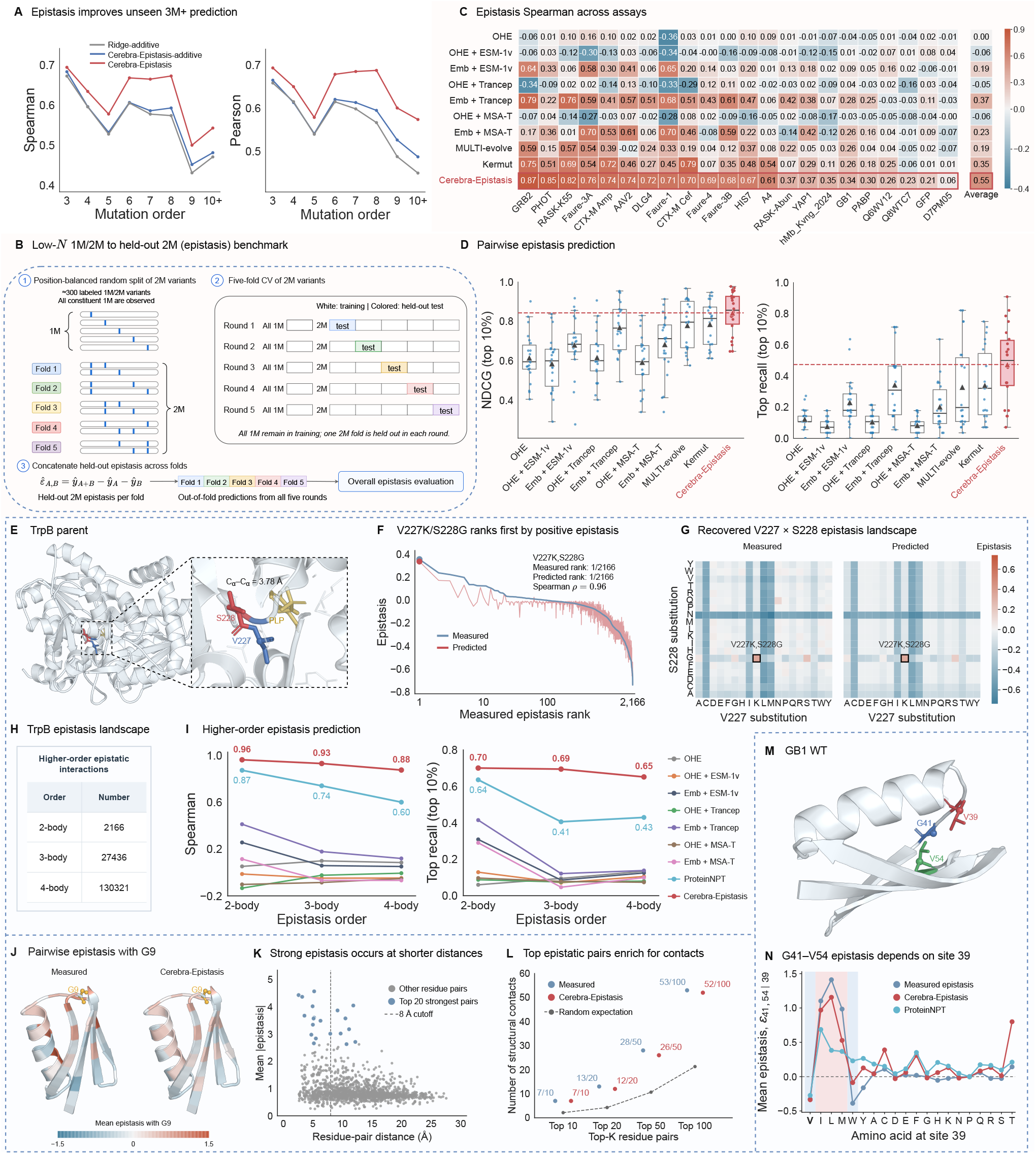
Cerebra-Epistasis predicts unseen epistasis and recovers biologically relevant interaction patterns. **A**, Cerebra-Epistasis, Cerebra-Epistasis-additive, and Ridge-additive on the Low-*N* 1M/2M-to-3M+ benchmark across mutation orders. **B**, Low-*N* 1M/2M-to-held-out-2M epistasis benchmark; all constituent single mutants remain in training while double-mutant combinations are held out in five position-balanced folds. **C**, Assay-level Spearman correlations for held-out pairwise epistasis prediction. **D**, Assay-level NDCG@10% and top recall@10%. Points denote assays, triangles denote assay means, and red dashed lines indicate the Cerebra-Epistasis means. **E**, Structural context of V227 and S228 in TrpB near the PLP-containing active site. **F**, Measured and predicted epistasis of 2,166 TrpB double mutants ordered by measured epistasis; V227K/S228G ranks first in both. **G**, Measured and predicted V227×S228 epistasis across amino-acid substitutions. **H**, Numbers of measured TrpB variants used to derive two-, three-, and four-body epistasis. **I**, Prediction of two-, three-, and four-body epistasis on the TrpB landscape. **J**, G9-centered epistasis mapped onto the GB1 structure. For each partner residue, epistasis was averaged across all available substitution pairs with G9 and centered across partner residues before structural mapping. **K**, Mean absolute epistasis for GB1 residue pairs versus spatial distance; blue points denote the 20 strongest measured pairs and the dashed line marks the 8-Å contact cutoff. **L**, Structural-contact enrichment among residue pairs ranked by mean absolute epistasis. The dashed line denotes the random expectation. **M**, GB1 wild-type structure highlighting sites V39, G41, and V54. **N**, Background-dependent G41–V54 epistasis. For each amino acid at site 39, conditional epistasis was calculated for all available G41/V54 substitution pairs and averaged. Shaded bands indicate the standard error of the mean (SEM).

To directly assess whether Cerebra-Epistasis could predict epistatic interactions in unseen mutation combinations, we designed a Low-*N* 1M/2M to held-out 2M (epistasis) benchmark. The constituent single mutations of each test double mutant were observed during training, while their specific combination was held out, thereby isolating prediction of unseen pairwise epistasis (Fig 4B; Supplementary Section S2.8). Cerebra-Epistasis substantially outperformed all applicable baselines across correlation- and ranking-based metrics (Fig 4C,D; Supplementary Table S10). It achieved a mean Spearman correlation of 0.55, compared with 0.35 for Kermut and 0.19 for MULTI-evolve, and a top-10% recall of 0.47, compared with 0.34 for Kermut and 0.33 for MULTI-evolve. Kermut [13] remained relatively competitive among the baselines, suggesting that 3D structural context is informative for modeling mutational coupling. Cerebra-Epistasis achieved further gains by explicitly modeling epistatic effects.

We next examined whether Cerebra-Epistasis could recover experimentally observed and biologically meaningful epistatic patterns across diverse protein systems. TrpB, the *β*-subunit of tryptophan synthase, catalyzes the final step of tryptophan biosynthesis. Johnston et al. [23] mapped a complete combinatorial fitness landscape across four active-site residues, including all double-, triple-, and quadruple mutants, enabling pairwise and higher-order epistasis to be quantified (Fig 4H). Across this landscape, Cerebra-Epistasis recovered epistatic effects from two- to four-body interactions and achieved the strongest performance across epistasis orders (Fig 4I). At the pairwise level, our reanalysis of the experimental landscape identified V227K/S228G as the strongest positive epistatic interaction among 2,166 double mutants (Fig 4F). Cerebra-Epistasis likewise ranked this pair first and reproduced the V227×S228 epistatic landscape across substitutions (Fig 4F,G). Structural mapping further showed that V227 and S228 are spatially adjacent near the PLP-bound active site (Fig 4E).

Beyond individual epistatic interactions, we asked whether Cerebra-Epistasis could recover broader structural patterns of epistasis using the GB1 landscape. GB1 is a small domain of streptococcal protein G that binds the Fc region of IgG. Olson et al. [24] comprehensively mapped pairwise epistasis across the GB1 domain and analyzed its relationship with protein structure. Following Olson et al.’s G9-centered structural analysis, we applied the same analysis to the measured and predicted epistatic interactions and observed similar spatial patterns (Fig 4J). More broadly, residue pairs with stronger measured epistasis tended to occur at shorter spatial distances, with 7 of the 10 strongest pairs forming structural contacts. The predicted interactions showed similar distance dependence and contact enrichment (Fig 4K,L).

Finally, we asked whether Cerebra-Epistasis could recover an experimentally established pattern of background-dependent epistasis in GB1. Wu et al. [6] found that higher-order interactions were enriched among sites 39, 41, and 54, whose side chains are spatially coupled in the GB1 core (Fig 4M), and further showed that the interaction between G41 and V54 depends on the amino acid at site 39. Our reanalysis showed that the G41–V54 interaction exhibited positive epistasis in the V39I, V39L, and V39M backgrounds, but negative epistasis in the V39V and V39W backgrounds. Cerebra-Epistasis closely reproduced this background-dependent pattern across site-39 substitutions, whereas ProteinNPT captured it less consistently (Fig 4N).

### Cerebra-Epistasis enables one-shot fitness landscape prediction

Protein engineering often involves screening a large number of variants. The number of possible variants increases exponentially with mutational depth. For a protein of only 100 residues, the sequence space already contains approximately 1.86 × 10^14^ possible five-mutation variants and 1.06 × 10^26^ possible ten-mutation variants (Fig 5B). Such a combinatorial growth trend highlights the importance of efficiency for computational baselines. Conventional supervised predictors require candidate-dependent feature construction and model evaluation for each variant. In contrast, Cerebra-Epistasis encodes the WT protein only once to generate the single-mutation score tensor *S* and epistasis representation tensor *U*. The entire fitness landscape can then be scored through lightweight indexing and assembly of the corresponding mutation components, without repeated sequence or structure encoding (Fig 5A).

**Figure 5.**
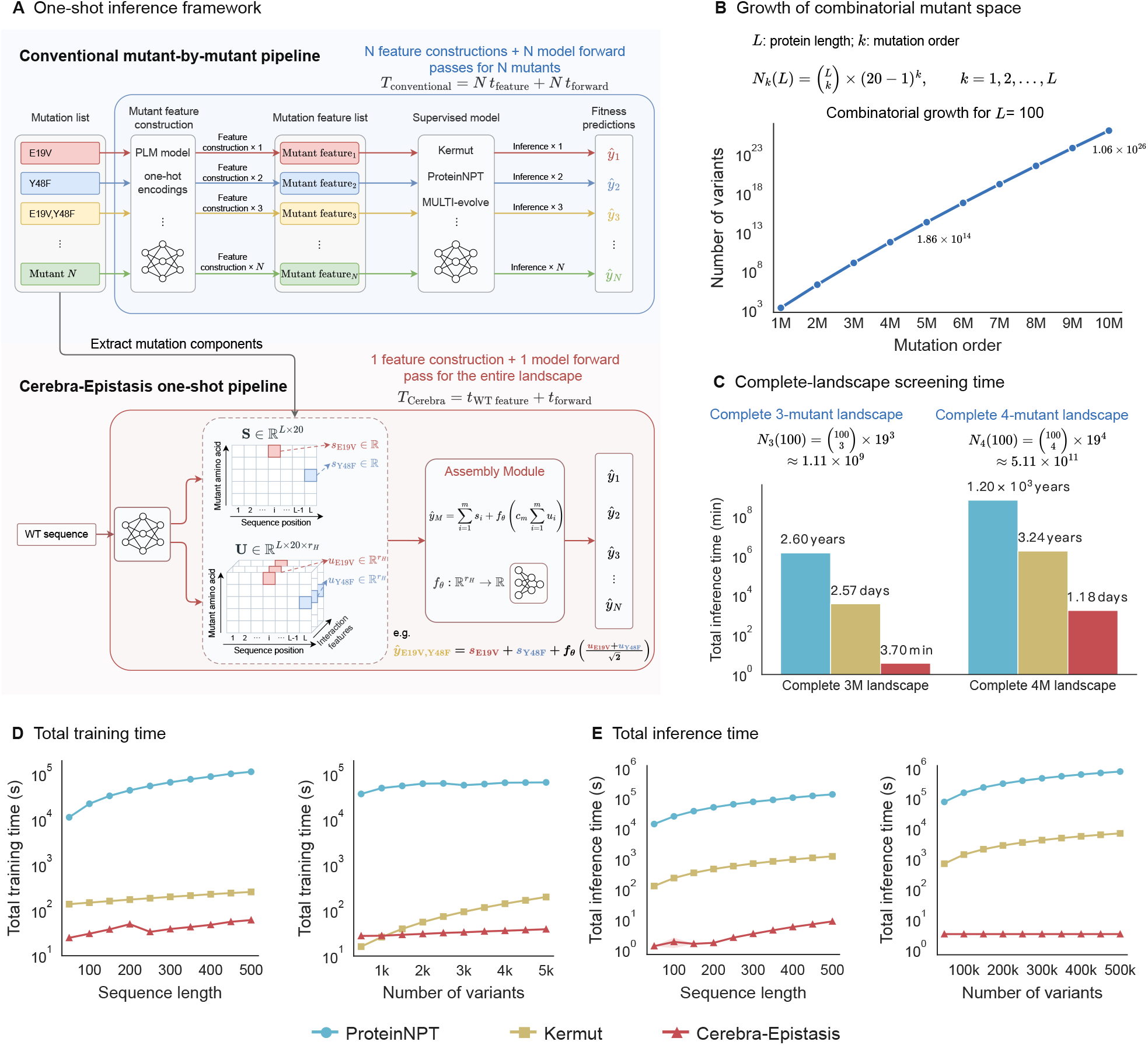
One-shot inference for fitness landscape prediction. **A**, Comparison of conventional mutant-by-mutant prediction with the one-shot inference paradigm of Cerebra-Epistasis. **B**, Combinatorial growth of mutant space for a protein of length *L* = 100. **C**, Estimated inference time for screening complete 3M and 4M landscapes at *L* = 100, extrapolated from measured inference throughput. **D**, Overall training time, including feature generation and model training, with increasing sequence length at 5,000 labeled variants (left) and increasing training-set size at *L* = 300 (right). **E**, Overall inference time, including feature generation and prediction, with increasing sequence length for 50,000 candidates (left) and increasing candidate-library size at *L* = 300 (right).

To quantify this computational advantage, we compared Cerebra-Epistasis with two of the strongest supervised fitness predictors, ProteinNPT and Kermut. For training, we adopted two modes: i) varying the protein length while sampling a fixed number of 5,000 labeled variants; and ii) varying the training-set size at a fixed protein length of 300 residues. During inference, we relaxed the number of candidate variants to 50,000 in mode (i), while varying the candidate-library size at a fixed length of 300 residues in mode (ii) (Fig 5D,E). Clearly, Cerebra-Epistasis required substantially less training time as protein length and training-set size increased, while its inference time remained low across sequence lengths and nearly constant as the candidate library grew. The computational gap widened further when screening complete higher-order landscapes. Based on the measured inference throughput, we estimated by extrapolation that screening the complete 3-mutant landscape of a length-100 protein would require only 3.70 min with Cerebra-Epistasis, compared with 2.57 days for Kermut and 2.60 years for ProteinNPT; for the complete 4-mutant landscape, the corresponding time was 1.18 days, 3.24 years, and 1.20 × 10^3^ years, respectively. These estimates correspond to an approximately 99.9% reduction in inference time relative to Kermut and a greater than 99.999% reduction relative to ProteinNPT at both mutation orders (Fig 5C).

### End-to-end training improves fitness prediction while preserving structural accuracy

Backpropagation through protein structure predictors incurs substantial computational and memory costs. Consequently, structure predictors and downstream fitness models are typically optimized separately. The lightweight Cerebra-Seq architecture makes end-to-end training with Cerebra-Epistasis computationally tractable. We therefore asked whether task-specific adaptation could improve fitness prediction without degrading the structural representations acquired during pretraining.

Reference proteins from the cDNA proteolysis multi-mutant dataset were clustered by sequence similarity and assigned, together with all associated variants, to five folds. In each round, end-to-end-trained and frozen-Cerebra-Seq models were trained on four folds and evaluated on the held-out fold using identical partitions. Further details of the protein-level data split and training protocol are provided in Supplementary Section S3.5.

The two settings exhibited distinct training trajectories (Fig 6A). End-to-end training maintained higher five-fold mean Spearman correlations after early crossings of the training trajectories. Under the reported 100-epoch schedule, the end-to-end-trained model achieved a five-fold mean Spearman correlation of 0.72 ± 0.02, compared with 0.68 ± 0.02 for frozen Cerebra-Seq. Frozen Cerebra-Seq peaked at 0.71 at epoch 36, whereas the end-to-end-trained model reached its maximum of 0.73 at epoch 54. These trajectories support a sustained benefit of end-to-end training for fitness prediction on held-out cDNA proteins during later training, and indicate alleviation of late-training overfitting.

**Figure 6.**
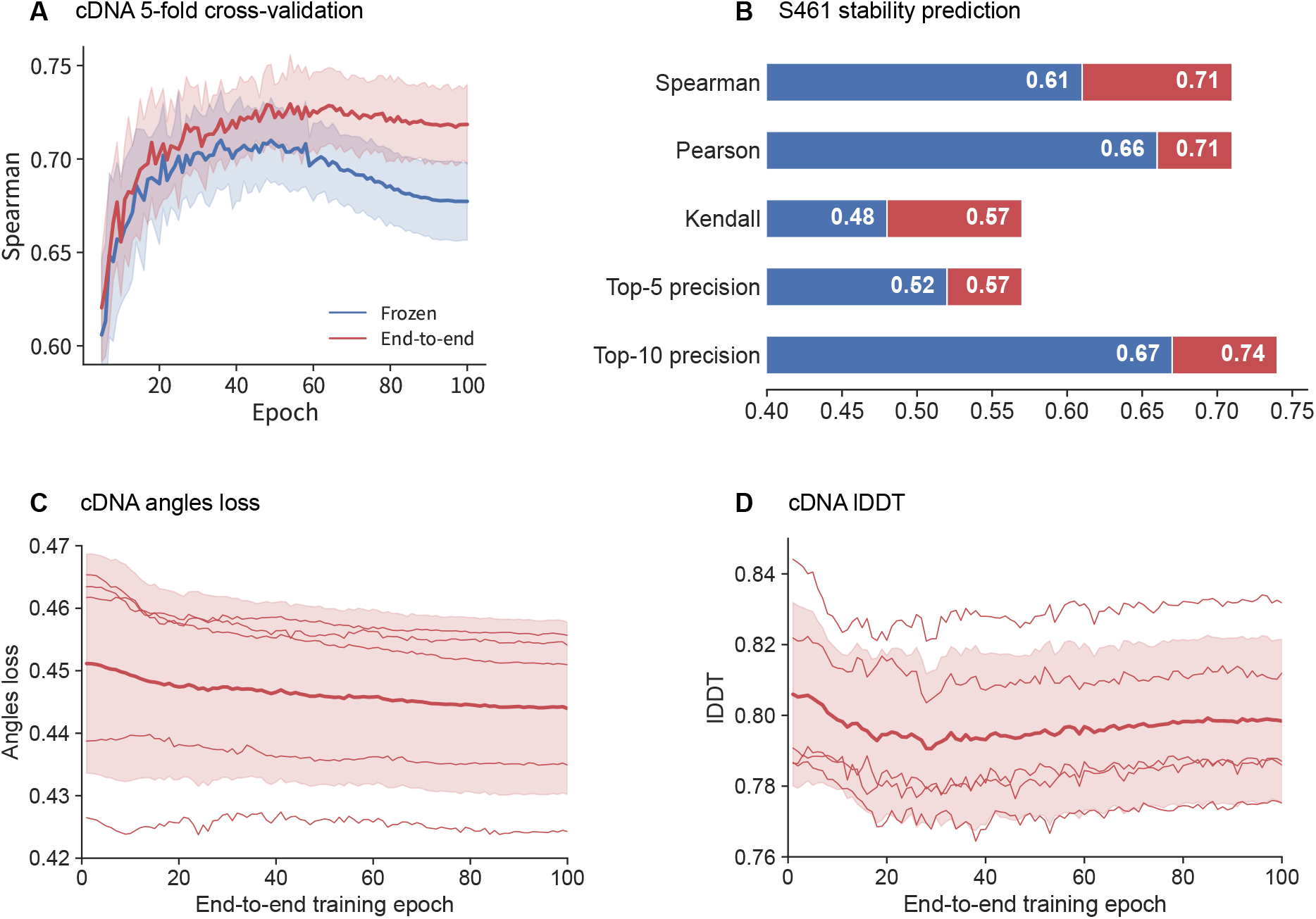
Effect of end-to-end training on downstream fitness prediction and structure prediction. **A**, Spearman correlations on held-out cDNA proteins in five-fold cross-validation, with splits defined by sequence-similarity clusters. Blue curves show the frozen-Cerebra-Seq baseline, in which only the downstream model was trained; red curves show end-to-end training of both components. Both settings used identical partitions. Curves show the five-fold mean at shared evaluation epochs, and shading indicates one ± standard deviation. **B**, Prediction performance on the independent S461 stability benchmark after training on the complete cDNA proteolysis dataset. Blue bars show the frozen-Cerebra-Seq baseline, and red extensions indicate the improvement from end-to-end training. Numeric labels report the mean scores across proteins for both settings. **C**, Side-chain *χ* torsion-angle loss during end-to-end training; lower values indicate better predictions. **D**, lDDT of the predicted structures during end-to-end training. In **C** and **D**, metrics were computed on proteins with experimental structures in each held-out fold. Epoch 0 denotes the pretrained Cerebra-Seq baseline prior to end-to-end training. Thin red curves represent individual folds, the thick red curve shows the five-fold mean, and red shading indicates ± one standard deviation.

To assess whether this pattern extended beyond the cDNA proteins, we trained both settings on the complete dataset and evaluated them on the independent S461 ΔΔ*G* benchmark (Fig 6B; Supplementary Table S11). End-to-end training increased mean Spearman, Kendall and Pearson correlations from 0.61, 0.48 and 0.66 to 0.71, 0.57 and 0.71, respectively. Similarly, the Top-5 and Top-10 precision rose from 0.52 and 0.67 to 0.57 and 0.74, respectively. The end-to-end-trained model matched the highest mean Spearman and Kendall correlations and achieved the highest mean Pearson correlation and Top-10 precision, although its mean Top-5 precision remained below the benchmark-leading value of 0.63. These results extend the cross-validation observation by showing improved performance after full-dataset training, when evaluated on an external protein-stability benchmark.

We next asked whether these fitness gains came at the cost of the pretrained structural representation. Structural performance was evaluated on held-out proteins with experimentally determined structures, with epoch 0 corresponding to the pretrained baseline before end-to-end training (Fig 6C,D). Mean side-chain *χ* torsion-angle loss decreased from 0.45 ± 0.02 to 0.44 ± 0.01 by epoch 100, corresponding to a reduction of 2.60 %. Mean lDDT remained between 0.79 and 0.81 and ended at 0.80 ± 0.02, comparable to the baseline range of 0.80 ± 0.03. Ablations showed that fitness gradients improved prediction beyond structural supervision alone, while additional structural supervision modestly improved fitness prediction and helped preserve structural accuracy (Supplementary Fig S5). Taken together, under the reported schedules, end-to-end training yielded higher final-epoch cDNA and S461 performance while modestly improving the side-chain torsion sampling and largely preserving the local structural quality.

## Discussion

Accurate construction of the multi-mutant fitness landscape requires epistasis to be modeled across varying mutation orders without increasing landscape-scale inference cost. Cerebra-Epistasis addresses these linked requirements through a reusable mutation-indexed atlas. The assembly module retrieves the relevant components to predict each mutant’s fitness and its epistatic contributions. Across the evaluated benchmark settings, Cerebra-Epistasis improves the overall multi-mutant fitness prediction over leading baselines. Its growing advantage at increasing mutation orders further supports the importance of explicit epistasis modeling. When the same mutation-specific representations are assembled for different mutation sets, the predicted epistasis is supposed to reflect the influence of mutational background. This design bridges low-order interaction expansions and nonlinear variant-level predictors while avoiding repeated sequence and structure encoding for every candidate [6, 7, 9].

Earlier sequence–structure models show that structural context can improve variant-effect prediction [11, 12, 14]. Our structural-feature ablations extend this evidence to higher-order multi-mutant fitness extrapolation. Adding Cerebra-Seq coordinates, residue-level node representations and residue-pair edge representations to sequence features generally benefits prediction. End-to-end training of Cerebra-Seq and the fitness model further improves fitness prediction while largely preserving local structural accuracy, as measured by lDDT. These findings support end-to-end training of the structure predictor and fitness model rather than treating predicted structure solely as fixed preprocessing. However, Cerebra-Seq still requires intensive pretraining to maintain its competitive predictive power. Improving structural accuracy without sacrificing end-to-end trainability therefore remains an important development goal.

The held-out interaction benchmark and analyses of TrpB and GB1 suggest that the explicit epistasis output captures patterns beyond aggregate fitness. These examples include local structural coupling, enrichment of spatial contacts, and changes in interaction sign across sequence backgrounds. However, the WT-centered formulation places an important boundary on the mechanistic interpretation of these patterns. The mutation-indexed atlas is constructed from the WT sequence and its predicted structure. It therefore does not explicitly represent mutation-induced rearrangements, conformational ensembles, or changes in oligomeric state, ligands or binding partners. This limitation may matter most when mutations alter structural states or context-dependent molecular interactions. Future models could incorporate conformational ensembles, molecular context, or selective mutant-specific structural refinement, while retaining most of the computational benefit of atlas reuse.

One-time WT encoding shifts the expensive sequence and structure computation from large candidate libraries to a single reference protein. The assembly module then retrieves and combines the relevant atlas components for each candidate. This inference paradigm underlies the projected speed advantage over models that encode and evaluate each mutant separately. However, efficient assembly does not eliminate the combinatorial growth of the candidate space. Likewise, the dependence on double-mutant coverage shows that explicit epistasis modeling cannot replace informative experimental measurements. Uncertainty-calibrated ranking and active selection could focus computation and experiments on the most informative regions of the landscape. Prospective design cycles will be needed to determine whether fitness and epistasis predictions improve protein optimization beyond retrospective benchmarks. Within these limits, Cerebra-Epistasis provides a unified framework for structure-aware multi-mutant fitness prediction, explicit epistasis modeling, and efficient landscape-scale inference.

## Methods

Cerebra-Epistasis links single-sequence structure prediction with protein fitness landscape modeling through a composable, epistasis-aware mutation atlas (Fig 1). For a given target protein, Cerebra-Seq first predicts residue-level and pairwise representations together with side-chain-resolved coordinates. These outputs are encoded once by an SE(3)-Transformer and converted into two mutation-indexed tensors covering all 20 amino-acid states at every sequence position. An assembly module then retrieves the entries corresponding to a specified mutation set and predicts both its fitness and its collective epistatic contribution. This design provides explicit 3D context while avoiding repeated sequence and structure encoding for individual mutants.

### Cerebra-Seq architecture

#### Single-sequence feature construction

Cerebra-Seq was developed from our previously reported MSA-based Cerebra architecture [18]. The original model integrates evolutionary information from an MSA and reconciles parallel coordinate hypotheses through path synthesis attention (PSA). Cerebra-Seq retains the Evoformer and PSA-based structure-generation principles but removes the requirement for an MSA. Instead, frozen ESMC and ESM3 models provide complementary per-residue embeddings directly from the target sequence. Their concatenated 2,688-dimensional representation is projected to 2,560 channels and reshaped into ten streams of width 256.

To enable information exchange among these streams while preserving stable signal propagation, we introduced a stream-mixing head inspired by manifold-constrained hyper-connections (mHC) [21]. A learnable 10 × 10 matrix is projected by Sinkhorn normalization onto the space of doubly stochastic matrices and used to mix the streams:

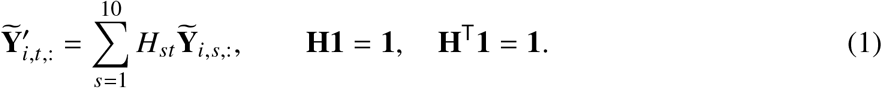

This module adapts the residual-stream mixing principle of mHC to protein language-model feature fusion rather than implementing the complete mHC architecture. Learnable amino-acid-type embeddings are added to the mixed streams to form the initial single representation. In parallel, relative sequence positions and the amino-acid identities of each residue pair initialize the pair representation. The single and pair representations are then jointly refined by the Evoformer trunk with recycling.

#### Structure generation and downstream representations

The refined representations are supplied to a PSA-based structure module derived from Cerebra. For multiple approximately uniformly distributed anchor residues, the module predicts residue translations and rotations in the corresponding local coordinate systems. PSA combines direct coordinate predictions with transformations propagated through intermediate anchors, integrating multiple internally consistent structural hypotheses [18]. Predicted residue frames and torsion angles are subsequently converted into side-chain-resolved atom14 coordinates.

Cerebra-Seq outputs a residue representation **N** ∈ ℝ^*L*×256^, a pair representation **E** ∈ ℝ^*L*×*L*×128^, and atom14 coordinates **X** ∈ ℝ^*L*×14×3^ for the downstream fitness model. Unless otherwise specified, Cerebra-Seq is frozen during assay-specific fitness training. In end-to-end training experiments, fitness gradients are additionally propagated through Cerebra-Seq, allowing its structure-aware representations to adapt to mutation-effect prediction. Detailed architectural and implementation settings are provided in Supplementary Algorithm S2.

### Downstream fitness model architecture

For a fitness landscape centered at the wild type, consider a mutant 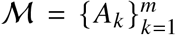 containing *m* substitutions, where each substitution *A*_*k*_ = (*i*_*k*_, *a*_*k*_) is specified by its sequence position *i*_*k*_ and mutant amino-acid state *a*_*k*_. Its fitness can be decomposed into two components: the summed contributions of its constituent single mutations and the combined contributions of epistatic interactions of different orders:

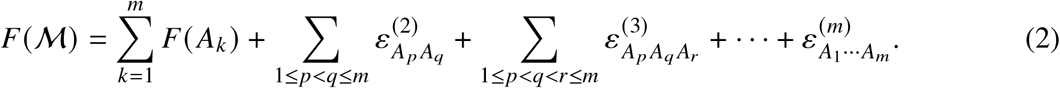

Here, *F* (*A*_*k*_) denotes the fitness contribution of single mutation *A*_*k*_ relative to the wild type, whereas *ε*^(*n*)^ denotes an *n*-body epistatic contribution. The first term therefore represents the total contribution of the constituent single mutations, while the remaining terms collectively represent the total epistatic contribution from pairwise to *m*-body interactions.

Motivated by this decomposition, Cerebra-Epistasis comprises two complementary branches: a single-mutation score branch that provides mutation-specific scalar scores and an epistasis representation branch that constructs mutation-specific representations for modeling collective epistatic contributions. Fig 1A,C and Supplementary Algorithm S1 summarize the overall workflow, which consists of three stages. First, Cerebra-Seq-derived structure-aware features of the wild-type protein are processed once by the SE(3)-Transformer encoder. Second, the resulting structure-aware representations, together with the residue-level ESM-2 embedding, are transformed into a single-mutation score tensor **S** ∈ ℝ^*L*×20^ and an epistasis representation tensor 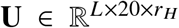. Third, for an arbitrary-order mutant, the corresponding entries are indexed from **S** and **U** and assembled to predict its fitness and total epistatic contribution.

#### SE(3)-Transformer encoding of the wild-type protein

Cerebra-Epistasis is designed as a structure-aware fitness prediction model. To effectively integrate 3D structural information while preserving the geometric transformation properties of the input, we employ an SE(3)-equivariant neural network to encode the Cerebra-Seq-derived structure-aware features of the wild-type protein. For a protein of length *L*, these features comprise Cerebra-Seq-derived residue representations **N** ∈ ℝ^*L*×256^, pairwise representations **E** ∈ ℝ^*L*×*L*×128^, and predicted atom14 coordinates **X** ∈ ℝ^*L*×14×3^. The encoder follows the general formulation of the SE(3)-Transformer [25].

For each central residue, the neighborhood includes spatially proximal residues together with residues exhibiting strong Cerebra pairwise representations. We introduce two task-specific adaptations to better integrate these features within the SE(3)-equivariant encoder. First, the input equivariant convolution constructs degree-1 geometric features from the atom14 coordinates of each neighboring residue expressed relative to the C_*α*_ coordinate of the central residue. This center-referenced representation preserves local atom-level geometry while removing dependence on global translation. Second, the projected Cerebra pairwise representations are combined with Fourier-encoded inter-residue distances to parameterize the radial components of the equivariant kernels, allowing SE(3)-equivariant message passing to incorporate both explicit geometry and learned residue-pair context.

The encoder outputs a degree-0 invariant representation **Z** ∈ ℝ^*L*×128^, a degree-1 equivariant representation **V** ∈ ℝ^*L*×32×3^, and local distance statistics **D**^stat^ ∈ ℝ^*L*×2^, comprising the mean and standard deviation of the C_*α*_ distances within the selected neighborhood of each residue. The wild-type protein is encoded only once, and the resulting representations are shared by the downstream mutation-specific branches. Detailed SE(3)-Transformer operations and implementation of the two adaptations are provided in Supplementary Algorithm S5, Supplementary Algorithm S14, and Supplementary Algorithm S13.

#### Construction of the single-mutation score and epistasis representation tensors

The encoded wild-type representations are subsequently transformed into two mutation-specific tensors covering all 20 amino-acid states at every sequence position: a single-mutation score tensor **S** ∈ ℝ^*L*×20^ and an epistasis representation tensor 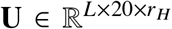. The former provides a scalar score for each position–amino-acid combination, whereas the latter provides a mutation-specific representation that can be assembled with other substitutions to model epistatic effects.

Specifically, the single-mutation score branch maps the invariant representation **Z** ∈ ℝ^*L*×128^ to sequence–structure-informed substitution scores and combines them with a stochastically gated correction derived from the residue-level ESM-2 embedding. In parallel, the epistasis representation branch converts the equivariant representation **V** ∈ ℝ^*L*×32×3^ into channel-wise magnitudes ∥**V**∥ ∈ ℝ^*L*×32^ and concatenates them with **Z** and the local distance statistics **D**^stat^ ∈ ℝ^*L*×2^. The resulting residue-level features are transformed by a multilayer perceptron (MLP) into a position-specific context tensor 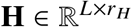. A learnable amino-acid embedding dictionary 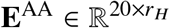 assigns an *r*_*H*_-dimensional representation to each amino-acid state. The position-specific context and amino-acid embeddings are combined through broadcasted element-wise multiplication to produce an intermediate mutation-specific tensor 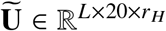, followed by a learnable linear mixing layer to obtain 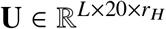.

Finally, both tensors are centered relative to the wild-type amino acid at each position by subtracting the corresponding wild-type score or representation from all amino-acid states, such that 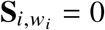 and 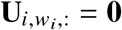. The resulting tensors therefore encode mutation-induced changes relative to the wild-type state and can be directly indexed for arbitrary-order mutant assembly. Details of the two tensor constructions are provided in Supplementary Algorithm S6 through Supplementary Algorithm S8.

#### Assembly and prediction of arbitrary-order mutants

For a mutant 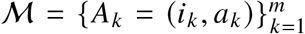, its mutation-specific score and epistasis representation are retrieved directly from the precomputed tensors:

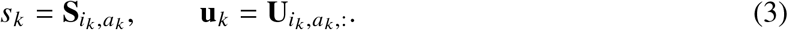

The selected epistasis representations are assembled through a permutation-invariant summation, optionally scaled according to mutation order, and the mutant fitness is predicted as

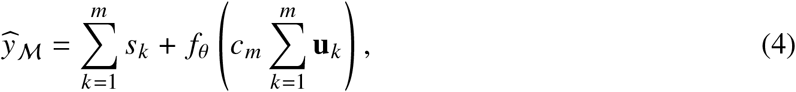

where *c*_*m*_ denotes the mutation-order scaling factor, with *c*_1_ = 1, and *f*_*θ*_ is a shared MLP that maps an epistasis representation to a scalar response.

To distinguish the contributions of individual substitutions from those arising through their joint assembly, the prediction can be decomposed into predicted single-mutation contributions and a total epistatic contribution:

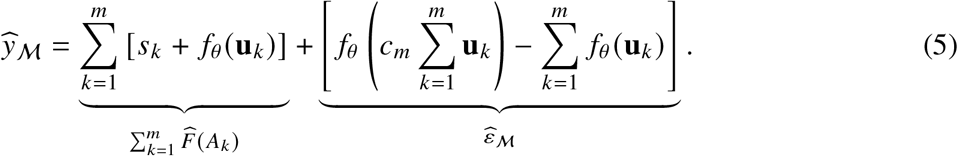

Here, 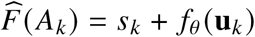 is the predicted single-mutation contribution of substitution *A*_*k*_. Although **U** is learned to support interaction-aware assembly, *f*_*θ*_ (**u**_*k*_) also provides a mutation-specific unary contribution. The predicted total epistatic contribution 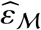 is defined after subtracting these constituent unary responses from their joint response. Consequently, 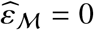 for a single mutant by construction. For a double mutant, 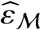 corresponds to predicted pairwise epistasis, whereas for higher-order mutants it represents the aggregate epistatic contribution beyond the sum of the constituent single-mutation contributions, consistent with the *n*-body fitness decomposition in Equation (2).

The permutation-invariant assembly and shared MLP enable Cerebra-Epistasis to model collective interactions among an arbitrary number of substitutions, independently of mutation order and without explicitly parameterizing individual pairwise or higher-order interaction terms. Once **S** and **U** have been constructed from a single encoding of the wild-type protein, large mutant libraries can be evaluated through tensor indexing and lightweight assembly without repeated sequence or structure encoding. Detailed assembly operations and the architecture of *f*_*θ*_ are provided in Supplementary Algorithm S9 and Supplementary Algorithm S10.

### Cerebra-Epistasis training strategy

Cerebra-Seq was first pretrained independently for protein structure prediction using structural supervision. Cerebra-Epistasis was subsequently trained using experimentally measured protein fitness data. Unless otherwise specified, Cerebra-Seq was kept frozen during downstream training; end-to-end training updated both Cerebra-Seq and the downstream fitness model. Full training objectives and implementation details are provided in the Supplementary Section S3, and benchmark-specific datasets and data-splitting protocols are described in Supplementary Section S2.

### Evaluation metrics

#### Metrics for structure prediction

We used TM-score [26] and lDDT [27] to evaluate the quality of predicted structures.

##### TM-score

TM-scores were calculated with the TM-score program [26]. TM-score measures the global topological similarity between a predicted structure and its reference after optimal alignment. The score is normalized by protein length and ranges from 0 to 1; values above 0.5 are generally considered to indicate the same overall fold.

##### Local distance difference test

The local distance difference test (lDDT) measures local structural accuracy without global superposition [27]. lDDT was computed over all C*α* atom pairs separated by less than 15 Å in the reference structure. For each pair, the absolute difference between the predicted and reference distances was compared against thresholds of 0.5, 1, 2 and 4 Å. The lDDT score is the mean, over the four thresholds, of the fraction of satisfied pairs.

#### Metrics for fitness prediction

For an assay containing *n* variants, let *y*_*i*_ and *ŷ*_*i*_ denote the measured and predicted fitness values of variant *i*, respectively. Fitness and epistasis prediction performance was evaluated using Pearson correlation, Spearman correlation, Kendall’s *τ*, NDCG (top 10%), and Top recall (top 10%). For the S461 protein stability benchmark, Top-5 and Top-10 precision were additionally used following the evaluation protocol of SPIRED-Stab [14]. For the avGFP experimental benchmark, NDCG (top 5) and Top recall (top 5) were used instead.

##### Pearson correlation

The Pearson correlation coefficient measures the linear relationship between measured and predicted values:

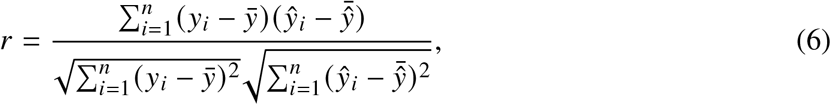

where 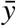 and 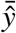 denote their respective means.

##### Spearman correlation

Spearman correlation measures the monotonic relationship between measured and predicted values and is calculated as the Pearson correlation between their ranks:

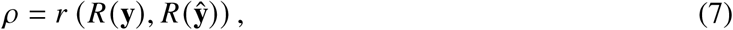

where *R*(·) denotes rank transformation.

##### Kendall’s rank correlation

Kendall’s *τ* measures ranking consistency based on concordant and discordant pairs. Kendall’s *τ*_*b*_, which accounts for ties, was used in all calculations.

##### NDCG

Following the ranking-based evaluation used for protein fitness prediction, measured fitness values were first converted to non-negative gains by min–max normalization:

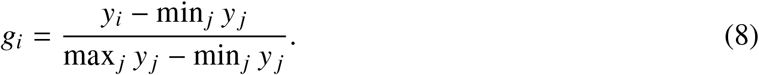

Let *K* denote the number of top-ranked variants considered, and let *π* _*j*_ denote the index of the variant at predicted rank *j*. The discounted cumulative gain over the top *K* predicted variants and its ideal value are defined as

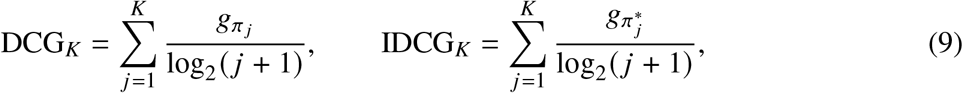

where 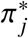 denotes the ranking obtained by sorting variants according to measured fitness. NDCG is then defined as

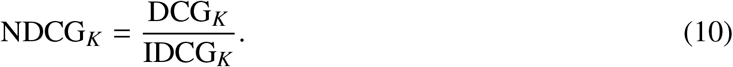

For the main fitness and epistasis benchmarks, *K* = ⌊0.1*n*⌋, corresponding to NDCG (top 10%). For the avGFP experimental benchmark, *K* = 5, corresponding to NDCG (top 5).

##### Top recall

Top recall measures the overlap between the top *K* variants according to experimental measurements and the top *K* variants according to model predictions. Let T_*K*_ and P_*K*_ denote these two sets, respectively. The metric is defined as

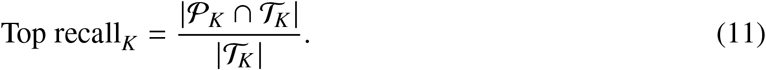

For the main fitness and epistasis benchmarks, *K* = ⌊0.1*n*⌋, corresponding to Top recall (top 10%). For the avGFP experimental benchmark, *K* = 5, corresponding to Top recall (top 5).

##### Top-*K* precision

For the S461 benchmark, Top-*K* precision measures the fraction of the *K* highest-ranked predicted mutations that are also among the *K* mutations with the highest experimentally measured values. Let *P*_*K*_ and *T*_*K*_ denote the predicted and experimentally determined top-*K* sets, respectively. The metric is defined as

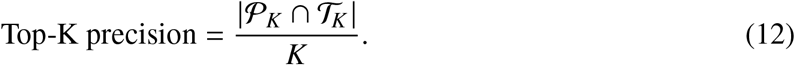

We used *K* = 5 and *K* = 10 for S461.

## Data availability

The benchmark datasets, precomputed input features, Cerebra-Epistasis training code, and prediction outputs generated by Cerebra-Epistasis and the baseline methods are publicly available on Zenodo at https://doi.org/10.5281/zenodo.22899137.

## Code availability

Cerebra-Seq and Cerebra-Epistasis were implemented in PyTorch. The source code and model weights for Cerebra-Seq are publicly available on Hugging Face at https://huggingface.co/GongLab-THU/Cerebra-Seq. The source code for Cerebra-Epistasis is publicly available on GitHub at https://github.com/xulab-research/Cerebra-Epistasis. A web server for online Cerebra-Epistasis prediction is available at https://structpred.life.tsinghua.edu.cn/server_cerebra_epistasis.html.

## Acknowledgements

This work was supported by the National Natural Science Foundation of China grant 32671644 (H.G.) and 32500561 (Y.X.), the Ministry of Science and Technology of China grant 2023YFF1204400 (H.G.), the Beijing Frontier Research Center for Biological Structure, the Postdoctoral Fellowship Program of CPSF under Grant Number GZC20251844 (Y.X.), the China Postdoctoral Science Foundation under Grant Number 2026T190759 (Y.X.) and 2026M793139 (Y.X.) and the General Program of Hubei Provincial Natural Science Foundation of China grant 2026AFB647 (Y.X.).

## Author contributions

H.G. and Y.X. conceived the study, proposed the methodology and developed the overall research framework. For Cerebra-Seq, H.G., W.W. and J.H. designed the initial framework, while W.W. implemented the model, performed model training and testing, and developed the web server. For Cerebra-Epistasis, Y.X. and Z.Y. designed the initial framework, while Z.Y. implemented the model and performed model training and testing. E.Y., Z.S. and S.Y. curated and preprocessed the datasets and conducted the benchmark evaluations. H.G., Y.X., W.W., Z.Y. and Z.S. analyzed the results and wrote the manuscript. All authors reviewed and approved the final manuscript.

## S1 Datasets of Cerebra-Epistasis

### S1.1 Datasets for Cerebra-Seq model training and evaluation

#### Training datasets

We trained Cerebra-Seq using experimental structures from the OpenFold PDB training set and a separately curated AlphaFold Protein Structure Database (AFDB) distillation set. The original OpenFold PDB set contained ~130,000 protein monomers. After excluding samples shorter than 50 residues, ~116,000 PDB samples remained for training.

The AFDB distillation set was constructed from AlphaFold-predicted structures for final training [1]. We selected approximately 2.30 million candidate proteins from the published non-singleton structural cluster representatives obtained by Foldseek clustering of AFDB50 [2]. For each candidate, we downloaded the AlphaFold model_v4 structure in PDB format. By using DSSP secondary-structure assignments [3], we trimmed contiguous coil regions from both termini. We calculated the mean pLDDT and radius of gyration (*R*_*g*_) over the retained region. We retained only structures with a mean pLDDT of at least 80, an effective trimmed length greater than 30 residues, and a full sequence length below 800 residues. We also required the trimmed structures to satisfy the length-dependent compactness criterion

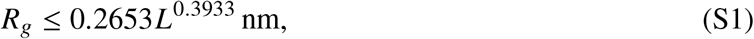

where *L* denotes the effective length of the trimmed region in residues. This length-dependent empirical upper bound was derived from statistics of monomeric PDB structures. These filters yielded 583,100 protein structures for the AFDB distillation set, which was incorporated during the final fine-tuning stage.

#### Test datasets

Since the OpenFold dataset contains PDB structures released by December 2021, we used the annual CAMEO datasets from the two most recent complete calendar years as the primary test sets to evaluate our model: CAMEO 2024 (2024.01–2025.01) and CAMEO 2025 (2025.01–2026.01). After excluding targets shorter than 50 amino acids, we retained 642 and 573 protein targets, respectively, for evaluating the structure-prediction accuracy of Cerebra-Seq and the baseline models.

### S1.2 Fitness and epistasis datasets for Cerebra-Epistasis training and evaluation

#### cDNA proteolysis dataset

Tsuboyama et al. developed a high-throughput cDNA display proteolysis assay to measure the thermodynamic folding stability of protein variants [4]. In this assay, proteins covalently linked to their corresponding cDNA were subjected to proteolysis, and the abundance of intact proteins was quantified by deep sequencing to infer folding stability. The resulting dataset contains large-scale stability measurements for single amino-acid variants and selected multiple mutants across hundreds of natural and designed protein domains.

For the **cDNA proteolysis dataset**, the `mut_type` field was used as the mutation annotation and the `ddG_ML` field as the mutation-effect label. Records for which `ddG_ML` could not be converted to a numeric value were removed. Records containing synonymous substitutions, insertions, or deletions were also excluded. Label values were then averaged across records with identical mutation annotations for the same protein, and records annotated as `wt` were subsequently excluded.

#### ArchStabMS1E10 Epistasis dataset

Faure et al. constructed large-scale combinatorial mutagenesis libraries to investigate the effects of higher-order amino-acid substitutions on protein stability and function [5]. Variant phenotypes were quantified using protein-fragment complementation assays, including AbundancePCA to measure protein abundance and, for selected libraries, BindingPCA to measure ligand binding, enabling systematic characterization of mutational effects and epistatic interactions across high-dimensional protein sequence spaces.

For the **ArchStabMS1E10 Epistasis dataset**, we retained libraries `1_lenient`, `2`, `3`, and `4`, and excluded library `1`, which contained fewer records and was a subset of `1_lenient`. Within each library, mutation annotations were inferred by comparing the mutant sequences with the corresponding wild-type sequence.

#### Antitoxin ParD3 Epistasis dataset

Lite et al. constructed a combinatorially complete library covering all 20^3^ = 8,000 amino-acid combinations at three key interaction positions in the ParD3 antitoxin and quantified the toxin-neutralization capacity of the variants using cellular growth competition and deep sequencing [6]. The processed variant-effect data used in this study were obtained from the subsequently released CoVES dataset [7].

For the **Antitoxin ParD3 Epistasis dataset**, the corresponding wild-type sequence was obtained from the original publication and added to the dataset. Records containing nonsense mutations were excluded, and label values were averaged across records with identical mutation annotations. Relative mutation effects were calculated by subtracting the wild-type label from each mutant label. The mutation annotations were then applied to the wild-type sequence to generate the corresponding mutant sequences.

#### TrpB Epistasis dataset

Johnston et al. constructed a combinatorially complete fitness landscape comprising all 20^4^ = 160,000 amino-acid combinations at four residues near the active site of a thermostable tryptophan synthase *β*-subunit (TrpB) [8]. Variant fitness was quantified using a pooled-culture enrichment assay in tryptophan-auxotrophic *Escherichia coli*, in which TrpB catalytic activity supports cellular growth, and changes in variant frequencies were measured by deep sequencing.

For the **TrpB Epistasis dataset**, the corresponding wild-type sequence was obtained from the original publication and added to the dataset. Records containing nonsense mutations were excluded. The processed mutation annotations were subsequently applied to the wild-type sequence to generate the corresponding mutant sequences.

#### Human Myoglobin Epistasis dataset

Küng et al. constructed a deep mutational scanning dataset for human myoglobin (hMb) using yeast surface display, fluorescence-activated cell sorting (FACS), and next-generation sequencing [9]. The variant library contained thousands of single and double mutants, and variant expression levels were quantified as expression fitness scores based on their distributions across fluorescence-sorting bins. These expression fitness scores were further shown to correlate with the thermostability of soluble hMb variants, providing a large-scale dataset for characterizing mutation effects and epistatic interactions.

For the **Human Myoglobin Epistasis dataset**, the wild-type amino-acid sequence was added to the dataset. Codon-level mutation annotations were converted to amino-acid-level mutation annotations, while records in which a mutant codon encoded a stop codon were excluded. Label values were averaged across records with identical mutation annotations, and records containing synonymous substitutions were further excluded.

#### CTXM Epistasis dataset

Judge et al. performed large-scale pairwise deep mutational scanning across 17 active-site residues of the CTX-M-14 *β*-lactamase to characterize epistatic interactions between amino acid substitutions [10]. Variant fitness was quantified by functional selection of *Escherichia coli* expressing the mutant enzymes in the presence of cefotaxime or ampicillin, followed by next-generation sequencing to measure changes in variant frequencies. The resulting fitness measurements quantify the relative antibiotic-resistance function of single and double mutants and enable systematic analysis of pairwise epistasis within the enzyme active site.

For the **CTXM Epistasis dataset**, low-confidence mutation records were excluded. Each original record was expanded into its two constituent single substitutions and the corresponding double substitution. The corresponding wild-type sequence was obtained from the original publication and added to the dataset. Mutation positions were then converted from Ambler numbering to the corresponding positions in the protein sequence. Within each antibiotic-selection condition, label values were averaged across records with identical mutation annotations.

#### ProteinGym DMS Substitutions dataset

Notin et al. developed ProteinGym, a large-scale benchmark that integrates deep mutational scanning datasets from diverse proteins and experimental systems for evaluating protein variant-effect prediction and protein design methods [11]. ProteinGym standardizes mutation annotations and experimental measurements across assays, with the direction of the `DMS_score` normalized such that higher values consistently indicate better variant fitness or performance in the measured phenotype, while the specific biological interpretation of the score remains assay dependent.

For the **ProteinGym DMS Substitutions dataset**, mutation annotations were validated and standardized. Within each assay, the wild-type sequence was inferred from the mutation annotations and corresponding mutant sequences, and the consistency of the inferred wild-type sequences was verified to ensure compatibility between the mutation annotations and mutant sequences.

For all datasets, the processed data were organized into separate directories for individual reference entries. Each directory contained three files: `data.csv`, `wt.fasta`, and `metadata.json`. The `data.csv` file contained three fields: `mutation_name`, representing the standardized mutation annotation; `mutated_sequence`, representing the corresponding mutant sequence; and `label`, representing the mutation-effect measurement. All mutation positions were standardized to 0-based indexing. The `wt.fasta` file contained the corresponding wild-type sequence, whereas the `metadata.json` file contained summary statistics for the corresponding mutation dataset. Detailed implementations of these preprocessing procedures are available in MutCleaner [12].

## S2 Cerebra-Epistasis benchmark settings

We evaluated the structural accuracy and computational efficiency of Cerebra-Seq, the single-sequence structure predictor within Cerebra-Epistasis. We assessed the complete framework through benchmarks for fitness and epistasis prediction.

For fitness and epistasis evaluation, we used three sources of mutational fitness data containing multi-mutant measurements. The cDNA proteolysis dataset contributed 154 assays. After removing assays overlapping with the cDNA proteolysis dataset, ProteinGym contained 19 assays. We also curated an Epistasis dataset from published studies and associated data repositories; after removing overlaps with ProteinGym, this collection contained 10 assays. All available assays were used in the benchmarks and were assigned to individual evaluation settings according to their mutation-order composition and the requirements of each benchmark. Details of dataset collection, curation, and preprocessing are provided in Supplementary Section S1.2. Here, 1M and 2M denote single- and double-mutant variants, respectively; 3M+ denotes variants containing three or more substitutions, and 1M+ denotes variants containing one or more substitutions. For order-averaged evaluation, only assay–order groups containing at least 15 test variants were included; metrics were then averaged equally across eligible mutation orders within each assay and subsequently across assays. The experimental settings for all benchmarks are summarized in Supplementary Table S1.

### S2.1 Structure prediction benchmark

The structure prediction benchmark was conducted on the CAMEO protein test sets described in Supplementary Section S1.1. Cerebra-Seq, ESMFold, OmegaFold, and SPIRED were run for four forward cycles, including the initial pass. For Cerebra-Seq, five checkpoints selected based on the training curves were used for prediction. Baseline checkpoints and candidate-generation settings, including the ESM3 sampling procedure, are described in Supplementary Section S4.1.

For each method and target, five candidate structures were generated and evaluated against the corresponding experimental reference structure. For ESMFold and OmegaFold, candidates from the two checkpoints were pooled within each method to form a single set of five structures. The highest TM-score among the five candidates was used as the target-level score for all methods (best-of-5 evaluation).

### S2.2 Structure inference time benchmark

Inference time was measured on a single NVIDIA A100 80GB GPU. Cerebra-Seq, ESMFold, OmegaFold, and SPIRED were each run for four forward cycles, including the initial pass. ESM3 was excluded from the inference-time benchmark because its iterative structure-generation protocol differed from the four-cycle prediction protocol used for the other models. Timing was collected after model loading and warm-up, and each measurement covered the complete inference workflow from sequence input to all-atom structure output. Measurement was repeated three times for each sequence length, and the mean inference time was reported.

For ESMFold and OmegaFold, chunking and subbatching were disabled, respectively, to exclude the associated computational overhead from the measurements. For SPIRED, the timed workflow included all-atom reconstruction by GDFold2 [13] with its default parameters.

### S2.3 1M+ to 1M+ fitness prediction benchmark

The 1M+ to 1M+ benchmark evaluates fitness prediction for single- and multi-mutant variants within the same assay. The benchmark comprised 29 assays, including 19 from ProteinGym and 10 from the Epistasis dataset, each containing both single- and multi-mutant variants. All available variants from each assay were included. The cDNA proteolysis assays were evaluated separately in a benchmark described below.

For each assay, variants were divided into five folds using a position-balanced random splitting strategy. Compared with a fully random split, this procedure approximately balanced the occurrence of each mutation position across folds.

In each cross-validation round, four folds were used for model training and the remaining fold was held out for testing. The procedure was repeated five times so that each variant was predicted exactly once in the held-out test fold. Predictions from the five test folds were concatenated to obtain out-of-fold predictions for the entire assay. Performance was evaluated both across all variants and separately by mutation order. All baseline methods were evaluated using the same five-fold partitions.

### S2.4 Low-*N* 1M/2M to 3M+ fitness prediction benchmark

Following the Low-*N* 1M/2M to 3M+ benchmarking strategy used in MULTI-evolve [14], we randomly sampled single mutants and their combinatorial variants from each assay. The benchmark comprised 18 assays, including 10 from ProteinGym and 8 from the Epistasis dataset. Only assays containing 1M, 2M, and 3M+ variants were included.

For each assay, the sampled 1M and 2M variants formed a training set containing fewer than 300 labeled variants, while the corresponding 3M+ combinatorial variants were reserved for testing. The 3M+ test variants represented unseen higher-order combinations composed exclusively of single mutations present in the training set. All baseline methods used the same training and test variants.

### S2.5 1M/2M to 3M+ fitness prediction benchmark

Consistent with the higher-order extrapolation setting used in MULTI-evolve [14], the full 1M/2M to 3M+ benchmark followed the same assay selection as the Low-*N* 1M/2M to 3M+ benchmark above, but without Low-*N* sampling. For each assay, all available single mutants were retained, multi-mutant variants were restricted to combinations of these single mutations, and all retained 1M and 2M variants were used for training while all retained 3M+ variants were reserved for testing. All baseline methods used the same training and test variants.

### S2.6 cDNA proteolysis multi-mutant fitness prediction benchmark

Among the 398 cDNA proteolysis assays, 154 containing multi-mutant variants were included in this benchmark. These assays were evaluated separately from the 1M+ to 1M+ benchmark because they share a common protein-stability-related label definition across proteins, providing a more homogeneous setting for cross-protein analysis.

For each assay, variants were divided into five folds using the same position-balanced random splitting strategy as in the 1M+ to 1M+ benchmark. In each cross-validation round, Cerebra-Epistasis was trained as a single shared model across all assays and evaluated on the corresponding held-out fold of each assay, whereas baseline methods were trained separately for each assay using the same five-fold splits. Details of the joint training procedure are provided in Supplementary Section S3.5.

Predictions from the five held-out test folds were concatenated for each assay before calculating assay-level performance. Performance was evaluated both across all variants and separately by mutation order.

### S2.7 avGFP experimental fitness prediction benchmark

The avGFP assay provided by ProteinGym [11] was used for model training, and 39 experimentally measured avGFP variants reported by Mi et al. [15] were used as an independent test set.

For Cerebra-Epistasis, 10% of the ProteinGym data were randomly held out as a validation set. A small number of candidate hyperparameter configurations were empirically evaluated, and the configuration with the highest validation Spearman correlation was selected. The model was then retrained on the complete ProteinGym avGFP dataset using the selected configuration and evaluated on the 39 experimental variants. No test-set measurements were used for hyperparameter selection.

### S2.8 Low-*N* 1M/2M to held-out 2M (epistasis) prediction benchmark

To evaluate pairwise epistasis prediction for unseen mutation combinations under limited training data, we constructed a Low-*N* 1M/2M to held-out 2M (epistasis) benchmark comprising 22 assays, including 15 from ProteinGym and 7 from the Epistasis dataset. For each assay, approximately 300–400 labeled 1M and 2M variants were sampled, with all retained double mutants composed exclusively of single mutations represented in the corresponding 1M set.

The 2M variants were divided into five folds using a position-balanced random splitting strategy, while all sampled 1M variants were retained in the training set of every cross-validation round. In each round, all sampled 1M variants together with four folds of 2M variants were used for training, and the remaining 2M fold was held out for testing. Thus, the constituent single mutations of each test double mutant were observed during training, while the specific pairwise combination was not.

For each held-out double mutant, predicted pairwise epistasis was calculated as the deviation of its predicted fitness from the additive expectation of its constituent single mutations, with experimental epistasis calculated analogously from the measured fitness values. This procedure was repeated for each of the five 2M folds so that every retained double mutant was predicted exactly once as a held-out test sample. The resulting out-of-fold epistasis predictions were concatenated for each assay before calculating assay-level performance. All baseline methods were evaluated using the same five-fold partitions.

### S2.9 S461 protein stability prediction benchmark

The S461 benchmark [16] is a manually corrected subset of the S669 benchmark [17] and contains 461 single-point mutations with experimentally measured protein stability changes (ΔΔ*G*). Only proteins containing at least 10 experimentally measured mutations were retained for evaluation.

Performance was evaluated separately for each retained protein and subsequently summarized across proteins.

**Table S1.** Overview of the Cerebra-Epistasis benchmark settings used for model evaluation.

| Benchmark | Dataset | Training data | Test data |
| --- | --- | --- | --- |
| 1M+ to 1M+ fitness | ProteinGym + Epistasis (29) | Four folds of 1M+ | Held-out fold of 1M+ |
| Low-*N* 1M/2M to 3M+ fitness | ProteinGym + Epistasis (18) | Sampled 1M + 2M (< 300) | 3M+ |
| 1M/2M to 3M+ fitness | ProteinGym + Epistasis (18) | All 1M + 2M | 3M+ |
| cDNA proteolysis multi-mutant fitness | cDNA proteolysis (154) | Four folds of 1M+ | Held-out fold of 1M+ |
| Low-*N* 1M/2M to held-out 2M (epistasis) | ProteinGym + Epistasis (22) | 1M + four folds of 2M | Held-out fold of 2M |
| S461 protein stability | S461 (12) | cDNA proteolysis | S461 |
| avGFP experimental fitness | avGFP | ProteinGym avGFP | 39 experimental variants |

## S3 Cerebra-Epistasis training and implementation details

### S3.1 Cerebra-Seq pretraining

The pretraining protocol for Cerebra-Seq was adapted from Cerebra [18]. We defined two PDB subsets: a 10K dataset containing 10,000 randomly selected monomers with sequence lengths between 100 and 400 residues and resolution better than 3 Å, and a 3 Å dataset containing all eligible PDB samples with resolution better than 3 Å. The full PDB training set comprised approximately 116,000 samples after length filtering. In the fourth stage, the model was fine-tuned on a mixture of the OpenFold PDB training set and the AFDB distillation set at a ratio of 1:2. The pLDDT prediction loss was applied only to PDB samples with experimental resolution better than 3 Å.

The training strategies are described in detail in Supplementary Table S2. The training objectives were adapted from Cerebra [18] with two modifications. First, we removed the masked-MSA prediction loss, as Cerebra-Seq uses single-sequence representations rather than MSA inputs. Second, for relative-rotation supervision, we replaced the component-wise *L*_1 error between the predicted and reference unit quaternions with the sign-invariant discrepancy.

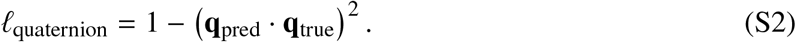

The inner product measures their angular agreement, and squaring it accounts for the equivalence of **q** and −**q** as representations of the same rotation. The resulting errors were aggregated as in the original Cerebra loss.

**Table S2.** Training strategy.

| Stage | Datasets | Loss function | Crop size | Side-chain prediction |
| --- | --- | --- | --- | --- |
| 1 | 10K PDB subset | $\mathcal{L}_{\text{Cerebra}}^{\text{init}}$ | 256 | × |
| 2 | 3 Å PDB subset | $\mathcal{L}_{\text{Cerebra}}$ | 256 | × |
| 3 | Full PDB training set | $\mathcal{L}_{\text{Cerebra}}^{\text{fine-tuning}}$ | 384 | ✓ |
| 4 | PDB + AFDB distillation data | $\mathcal{L}_{\text{Cerebra}}^{\text{fine-tuning}}$ | 384 | ✓ |

### S3.2 Downstream fitness model and training setup

Unless otherwise specified, the pretrained Cerebra-Seq module was kept frozen and only the downstream prediction network was optimized. The SE(3)-Transformer encoder consisted of an input ConvSE3 layer, followed by one SE(3)-Transformer block comprising an equivariant multi-head attention layer and a feed-forward layer, and an output ConvSE3 layer. The attention layer used four heads with a per-head dimension of 64. The hidden fibers contained 320 degree-0 channels and 32 degree-1 channels, whereas the output fibers contained 128 degree-0 channels and 32 degree-1 channels. The interaction function *f*_*θ*_ was implemented as an MLP with the layer dimension *r*_*H*_ → 256 → 128 → 1 and GELU activation after the first two linear layers, where *r*_*H*_ denotes the interaction rank.

Models were optimized using Adam without weight decay. Experimental fitness labels were standardized separately for each assay using the mean and standard deviation computed from the corresponding training partition. Models were trained to predict standardized fitness values, and predictions were transformed back to the original label scale using the corresponding training-set statistics before evaluation. The S461 protein stability benchmark was an exception, for which no label normalization was applied. Gradients were clipped to a maximum global norm of 2.0.

### S3.3 Fitness and epistasis training objectives

Training was based on a primary fitness prediction objective and, where applicable, an auxiliary pairwise epistasis objective. For a training set containing *N* variants, the fitness prediction loss was defined as

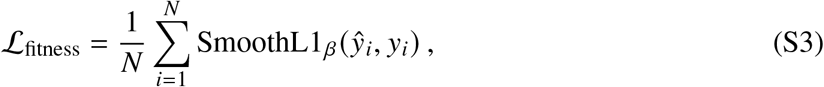

where *ŷ*_*i*_ and *y*_*i*_ denote the predicted and experimental fitness values on the training scale, respectively, and *β* = 1.0. For benchmarks using label normalization, the loss was evaluated in the standardized fitness space.

When both constituent single mutants of a training double mutant were available in the same training partition, an auxiliary pairwise epistasis target was constructed. Let 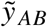 denote the standardized fitness of a double mutant and 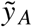 and 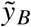 those of its constituent single mutants. The auxiliary pairwise epistasis target was defined as

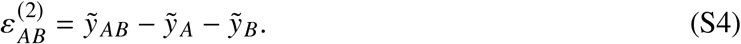

Consistent with the fitness assembly function, the corresponding model-predicted pairwise epistasis was defined as

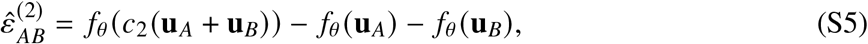

where 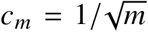 when mutation-order scaling was enabled and *c*_*m*_ = 1 otherwise. Only double mutants for which both constituent single-mutant measurements were present in the corresponding training partition were included in the auxiliary objective. The epistasis loss was then defined as

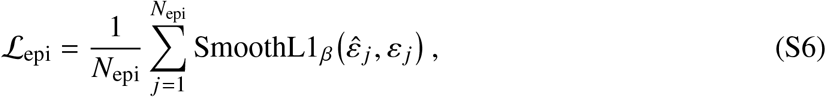

where *N*_epi_ is the number of eligible double mutants in the training partition. The overall training objective was

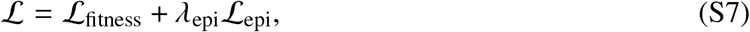

where *λ*_epi_ controls the contribution of explicit pairwise epistasis supervision. When *λ*_epi_ = 0, only the fitness prediction objective was optimized. Benchmark-specific values of *λ*_epi_ and the mutation-order scaling setting are summarized in Supplementary Table S3.

### S3.4 Benchmark-specific downstream training settings

Most training settings were shared across benchmarks. A small number of settings, including the learning rate, interaction rank *r*_*H*_, pairwise epistasis supervision, and mutation-order scaling, were selected separately for each benchmark. When hyperparameter selection or early stopping was required, a validation subset was held out from the training data for this purpose. The selected settings were then fixed and applied uniformly to all assays and data partitions within the same benchmark. Test data were never used for hyperparameter selection or early stopping. The final settings are summarized in Supplementary Table S3.

### S3.5 Specialized training strategies

#### Joint training across cDNA proteolysis assays

For the cDNA proteolysis multi-mutant benchmark, Cerebra-Epistasis was trained jointly across assays using a single shared model. During training, each assay contributed its own training variants and assay-specific loss to update the shared model parameters. Label normalization was performed separately for each assay using the mean and standard deviation computed from its corresponding training partition. The corresponding training configuration is provided in Supplementary Table S3.

**Table S3.** Benchmark-specific training settings.

| Benchmark | Epochs | Learning rate | $r_H$ | $c_m$ | $\lambda_{\text{epi}}$ |
| --- | --- | --- | --- | --- | --- |
| 1M+ to 1M+ | Early stopping | $5 \times 10^{-4}$ | 320 | 1 | 0 |
| cDNA proteolysis | 150 | $5 \times 10^{-4}$ | 320 | 1 | 0 |
| Low-$N$ 1M/2M to held-out 2M (epistasis) | 150 | $5 \times 10^{-4}$ | 320 | 1 | 2 |
| Low-$N$ 1M/2M to 3M+ | 150 | $1 \times 10^{-4}$ | 64 | $\frac{1}{\sqrt{m}}$ | 2 |
| 1M/2M to 3M+ | 150 | $1 \times 10^{-4}$ | 64 | $\frac{1}{\sqrt{m}}$ | 2 |

#### End-to-end training

End-to-end training was first evaluated on the 154 cDNA proteolysis assays containing multi-mutant measurements. This protein-level split differed from the within-assay, position-balanced split described in Supplementary Section S2.6. Reference protein sequences were grouped into five clusters using NW-align-based sequence similarities, without a predefined sequence-identity threshold. Each cluster defined one cross-validation fold, with all variants of each assay kept together.

In each cross-validation round, an independent model was trained on four folds and evaluated on the remaining fold. Cerebra-Seq was initialized from its pretrained weights and then trained end to end with the downstream fitness-prediction components on the training assays. In the frozen-Cerebra-Seq baseline, Cerebra-Seq was initialized from the same pretrained weights but kept fixed, so that only the downstream components were optimized. Both settings used identical partitions. Models were trained independently across the five rounds.

The downstream components were trained with a learning rate of 5 × 10^−4^. The pretrained Cerebra-Seq weights were fine-tuned at 5 × 10^−6^. Experimental PDB structures were available for 139 of the 154 reference proteins. Structural supervision was applied to training proteins with experimental structures, using the end-to-end training objective

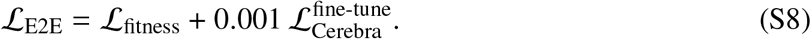

Here, *L*_fitness_ is the fitness prediction loss defined above. The structural term 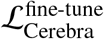 is the fine-tuning objective specified in Supplementary Table S2, incorporating the loss modifications described above. For the remaining 15 proteins without experimental structures, only the fitness prediction loss was applied. Structural metrics were evaluated on proteins with experimental structures in the held-out fold of each round.

To characterize training–test sequence similarity, we computed each held-out protein’s maximum sequence identity to any protein in the four training folds. Fold-specific nearest-neighbour identity statistics are provided in Supplementary Table S4. The maximum identity remained below 50% in every fold, with an overall maximum of 43.60%.

For the final S461 evaluation, we identified the best-performing epoch in each cDNA cross-validation round. The final training duration was determined from the arithmetic mean of these five epoch numbers. Models were then trained on all cDNA assays for the resulting number of epochs. They were evaluated on the independent S461 stability benchmark using the protocol described in Supplementary Section S2.9.

**Table S4.** Sequence similarity and structure-label availability across the cluster-based cDNA cross-validation folds. Nearest-neighbour (NN) identity is each held-out reference protein’s maximum sequence identity to any reference protein in the four training folds. The ≥ 40% column counts held-out assays meeting this identity threshold. Structure-label counts indicate reference proteins with experimental PDB structures used for structural supervision in the training folds or structural evaluation in the held-out fold.

| Test fold | Assays | Mean NN<br>identity (%) | Maximum NN<br>identity (%) | NN identity<br>$\geq 40\%$ (*n*) | Training<br>structure labels<br>(*n*) | Test<br>structure labels<br>(*n*) |
| --- | --- | --- | --- | --- | --- | --- |
| **Fold 0** | 44 | 33.18 | 43.60 | 5 | 104 | 35 |
| **Fold 1** | 19 | 33.73 | 43.60 | 3 | 122 | 17 |
| **Fold 2** | 33 | 33.20 | 38.90 | 0 | 108 | 31 |
| **Fold 3** | 29 | 34.79 | 42.90 | 2 | 110 | 29 |
| **Fold 4** | 29 | 34.17 | 41.90 | 3 | 112 | 27 |

## S4 Baseline methods and comparison settings

### S4.1 Structure prediction baselines

For structure prediction, Cerebra-Seq was compared with ESMFold [19], OmegaFold [20], ESM3 [21], and SPIRED [22]. ESMFold predictions used the officially released `esmfold_v0` and `esmfold_v1` checkpoints. OmegaFold predictions used the officially released `release1.pt` and `release2.pt` checkpoints. For ESM3, each candidate structure was generated in an independent run with 20 iterative sampling steps. Unless otherwise specified, all baseline models were run using their default settings.

### S4.2 Fitness and epistasis prediction baselines

For fitness and epistasis prediction, we included supervised baselines following the ProteinGym benchmark [11]. The one-hot encoding (OHE) baseline used one-hot-encoded mutant sequences as input to ridge regression. The OHE-based baselines additionally incorporated zero-shot fitness scores from ESM-1v [23], Tranception [24], or MSA Transformer [25]. The embedding-based baselines instead used mean-pooled sequence embeddings from each of these pretrained models, augmented with zero-shot fitness scores from the corresponding model, as inputs to ridge regression. We additionally compared Cerebra-Epistasis with MULTI-evolve [14], Kermut [26], and ProteinNPT [27]. All supervised baseline predictors were trained on the same training partitions as Cerebra-Epistasis and evaluated on the corresponding test partitions within each benchmark. The ProteinGym-derived baselines followed their published implementations [11, 27], while MULTI-evolve and Kermut followed their respective original implementations.

### S4.3 Fitness and epistasis prediction baselines

For fitness and epistasis prediction, we included supervised baselines following the ProteinGym benchmark [11]. The one-hot encoding (OHE) baseline used one-hot-encoded mutant sequences with a regularized linear prediction head. The augmented OHE baselines additionally used zero-shot fitness scores from ESM-1v [23], Tranception [24], or MSA Transformer [25] as prediction covariates. The embedding-based baselines processed residue-level embeddings from each of these pretrained models with a one-dimensional convolution followed by mean pooling, and used the corresponding zero-shot fitness score as an additional prediction covariate. We also compared Cerebra-Epistasis with MULTI-evolve [14], Kermut [26], and ProteinNPT [27].

All supervised baselines were trained on the same training partitions as Cerebra-Epistasis and evaluated on the corresponding test partitions within each benchmark. The ProteinGym-derived baselines used the publicly released ProteinNPT implementation [11, 27], while MULTI-evolve and Kermut followed their respective original implementations.

### S4.4 ΔΔ*G* prediction baselines

For protein stability prediction on the S461 benchmark, Cerebra-Epistasis was compared with established ΔΔ*G* prediction methods, including DDMut [28], ProteinEBM [29], PROSTATA [30],

RaSP [31], SPIRED-Fitness [22], GeoDDG-3D and GeoDDG-Seq [32], Mutate Everything (ESM-2) [33], ThermoMPNN [34], Pythia [35], SPURS [36], and SPIRED-Stab [22]. All baseline predictions were generated using the publicly available pretrained model weights and the corresponding original inference procedures. Performance was evaluated on the S461 benchmark following the protocol described in Supplementary Section S2.9.

## S5 Supplementary Results

### S5.1 Supplementary Results for 1M+ to 1M+

Results for the 1M+ to 1M+ benchmark described in Supplementary Section S2.3 are summarized in Supplementary Table S5. Cerebra-Epistasis achieved the strongest overall performance, ranking first on most evaluated correlation- and ranking-based metrics.

**Table S5.** Performance on the 1M+ to 1M+ benchmark under overall and order-averaged assay-level evaluation. Values are reported as mean ± SD across assays.

| Model | Spearman $\rho$ | Pearson $r$ | Kendall $\tau$ | NDCG<br>(top 10%) | Top recall<br>(top 10%) |
| --- | --- | --- | --- | --- | --- |
| **Overall assay-level evaluation** |  |  |  |  |  |
| OHE | 0.74 ± 0.18 | 0.73 ± 0.17 | 0.56 ± 0.17 | 0.88 ± 0.09 | 0.50 ± 0.17 |
| OHE + ESM-1v | 0.75 ± 0.18 | 0.75 ± 0.16 | 0.58 ± 0.16 | 0.89 ± 0.08 | 0.51 ± 0.18 |
| Embeddings + ESM-1v | 0.70 ± 0.20 | 0.72 ± 0.18 | 0.53 ± 0.19 | 0.86 ± 0.11 | 0.48 ± 0.20 |
| OHE + Tranception | 0.77 ± 0.18 | 0.76 ± 0.16 | 0.60 ± 0.17 | 0.90 ± 0.07 | 0.54 ± 0.17 |
| Embeddings + Tranception | 0.77 ± 0.19 | 0.80 ± 0.14 | 0.60 ± 0.18 | 0.90 ± 0.08 | 0.55 ± 0.18 |
| OHE + MSA Transformer | 0.78 ± 0.19 | 0.77 ± 0.16 | 0.60 ± 0.17 | 0.91 ± 0.07 | 0.55 ± 0.18 |
| Embeddings + MSA Transformer | 0.76 ± 0.18 | 0.79 ± 0.16 | 0.59 ± 0.18 | 0.90 ± 0.08 | 0.54 ± 0.18 |
| ProteinNPT | 0.84 ± 0.17 | 0.91 ± 0.12 | 0.69 ± 0.18 | 0.94 ± 0.07 | 0.67 ± 0.20 |
| Cerebra-Epistasis | **0.85 ± 0.17** | **0.91 ± 0.12** | **0.70 ± 0.17** | **0.95 ± 0.06** | **0.68 ± 0.19** |
| **Order-averaged assay-level evaluation** |  |  |  |  |  |
| OHE | 0.68 ± 0.17 | 0.68 ± 0.16 | 0.52 ± 0.15 | 0.86 ± 0.10 | 0.47 ± 0.14 |
| OHE + ESM-1v | 0.69 ± 0.16 | 0.69 ± 0.15 | 0.52 ± 0.14 | 0.86 ± 0.09 | 0.45 ± 0.14 |
| Embeddings + ESM-1v | 0.63 ± 0.21 | 0.65 ± 0.20 | 0.47 ± 0.18 | 0.81 ± 0.16 | 0.41 ± 0.16 |
| OHE + Tranception | 0.70 ± 0.16 | 0.70 ± 0.15 | 0.54 ± 0.14 | 0.87 ± 0.09 | 0.48 ± 0.14 |
| Embeddings + Tranception | 0.70 ± 0.17 | 0.73 ± 0.16 | 0.54 ± 0.16 | 0.86 ± 0.11 | 0.47 ± 0.14 |
| OHE + MSA Transformer | 0.71 ± 0.17 | 0.71 ± 0.16 | 0.55 ± 0.15 | 0.87 ± 0.09 | 0.49 ± 0.14 |
| Embeddings + MSA Transformer | 0.69 ± 0.17 | 0.71 ± 0.16 | 0.53 ± 0.15 | 0.85 ± 0.11 | 0.45 ± 0.15 |
| ProteinNPT | 0.78 ± 0.19 | **0.85 ± 0.16** | **0.63 ± 0.18** | 0.91 ± 0.09 | 0.60 ± 0.19 |
| Cerebra-Epistasis | **0.78 ± 0.17** | 0.85 ± 0.15 | 0.63 ± 0.17 | **0.92 ± 0.09** | **0.62 ± 0.18** |

### S5.2 Supplementary Results for Low-*N* 1M/2M to 3M+

Results for the Low-*N* 1M/2M to 3M+ benchmark described in Supplementary Section S2.4 are summarized in Supplementary Table S6. Cerebra-Epistasis achieved the strongest overall performance and retained its advantage under order-averaged evaluation.

**Table S6.**
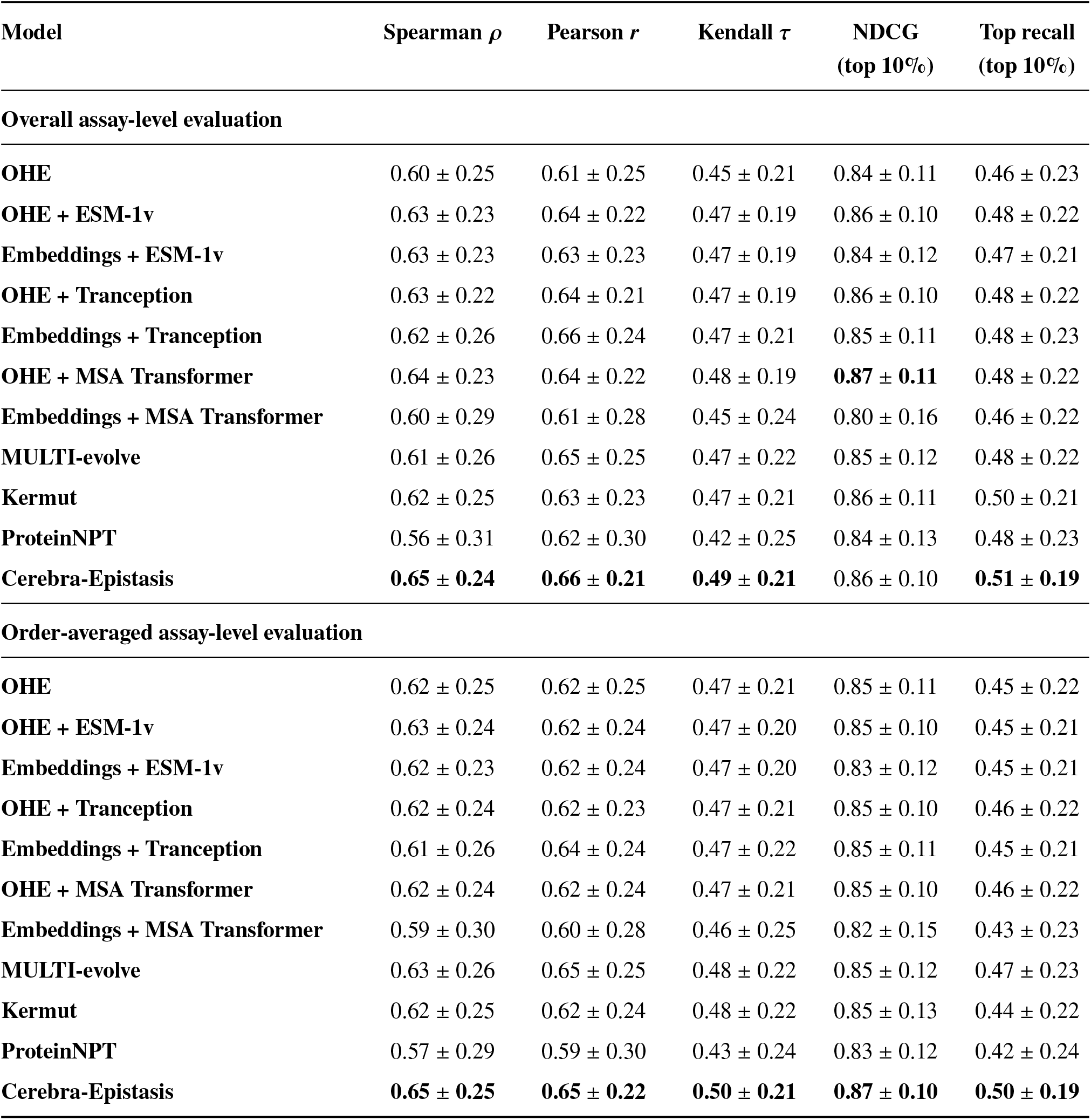
Performance on the Low-*N* 1M/2M to 3M+ benchmark under overall and order-averaged assay-level evaluation. Values are reported as mean ± SD across assays.

### S5.3 Supplementary Results for 1M/2M to 3M+

Following the 1M/2M to 3M+ benchmark setting described in Supplementary Section S2.5, we further evaluated higher-order extrapolation across the full collection of eligible assays.

Overall and order-averaged assay-level performance is summarized in Supplementary Table S7, where Cerebra-Epistasis achieved the best performance on most aggregate metrics. Performance at individual mutation orders is shown in Supplementary Fig S1. Although the performance of all methods generally decreased with increasing mutation order, Cerebra-Epistasis maintained the strongest overall performance across mutation orders.

**Figure S1.**
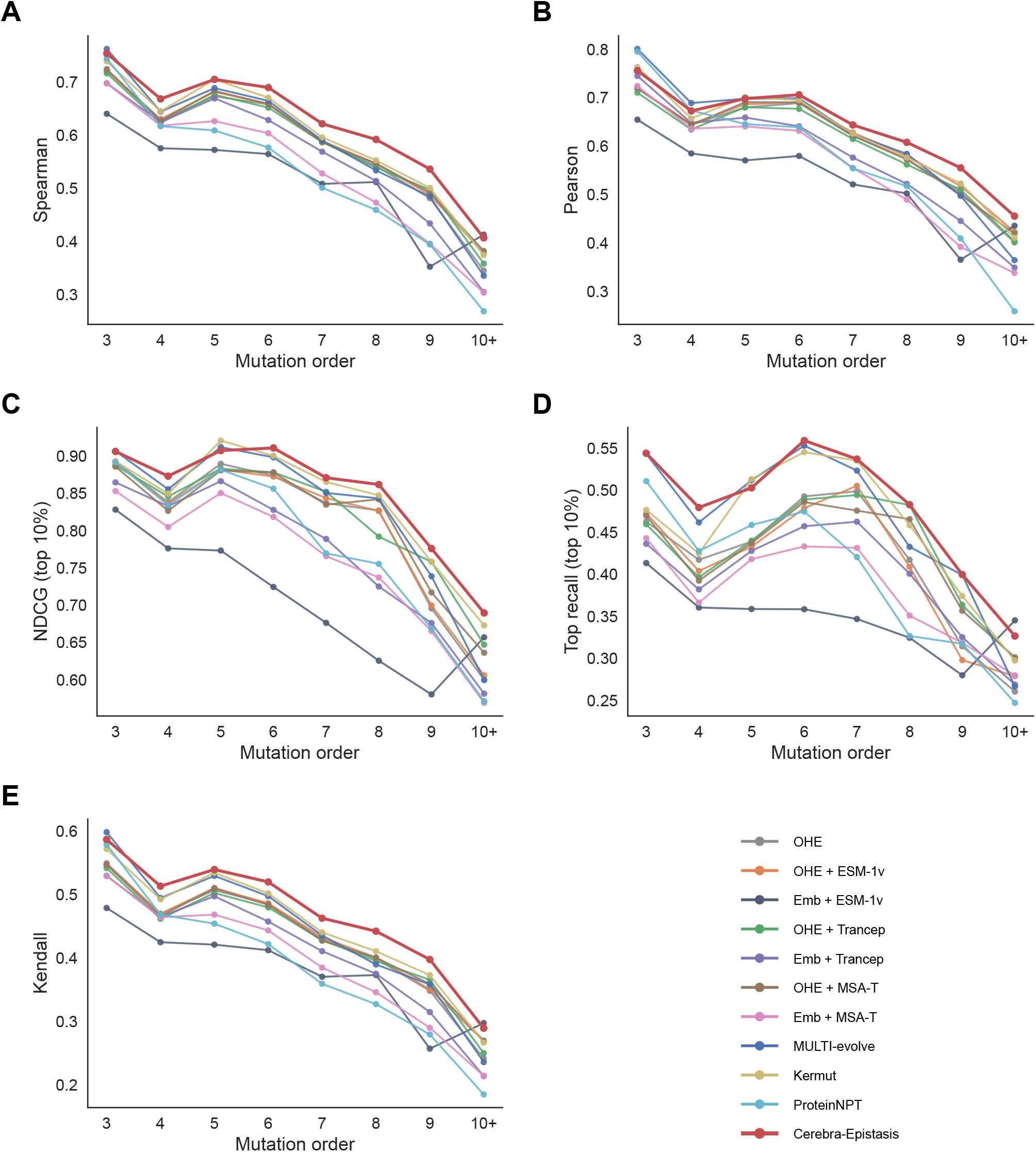
Mutation-order-resolved performance on the 1M/2M to 3M+ benchmark. Models were trained on single and double mutants and evaluated on held-out variants with three or more mutations. Curves show assay-averaged Spearman, Pearson, NDCG (top 10%), top recall (top 10%), and Kendall at each mutation order.

**Table S7.** Performance on the 1M/2M to 3M+ benchmark under overall and order-averaged assay-level evaluation. Values are reported as mean ± SD across assays.

| Model | Spearman $\rho$ | Pearson $r$ | Kendall $\tau$ | NDCG<br>(top 10%) | Top recall<br>(top 10%) |
| --- | --- | --- | --- | --- | --- |
| **Overall assay-level evaluation** |  |  |  |  |  |
| OHE | 0.57 ± 0.22 | 0.58 ± 0.19 | 0.42 ± 0.19 | 0.80 ± 0.13 | 0.39 ± 0.23 |
| OHE + ESM-1v | 0.62 ± 0.20 | 0.63 ± 0.18 | 0.46 ± 0.17 | 0.83 ± 0.14 | 0.45 ± 0.23 |
| Embeddings + ESM-1v | 0.59 ± 0.21 | 0.59 ± 0.17 | 0.43 ± 0.18 | 0.79 ± 0.16 | 0.40 ± 0.22 |
| OHE + Tranception | 0.65 ± 0.20 | 0.64 ± 0.16 | 0.48 ± 0.17 | 0.86 ± 0.12 | 0.46 ± 0.21 |
| Embeddings + Tranception | 0.63 ± 0.21 | 0.65 ± 0.14 | 0.46 ± 0.18 | 0.84 ± 0.12 | 0.43 ± 0.20 |
| OHE + MSA Transformer | 0.66 ± 0.20 | 0.65 ± 0.16 | 0.49 ± 0.16 | 0.87 ± 0.12 | 0.46 ± 0.19 |
| Embeddings + MSA Transformer | 0.61 ± 0.24 | 0.61 ± 0.20 | 0.46 ± 0.21 | 0.79 ± 0.17 | 0.41 ± 0.22 |
| MULTI-evolve | 0.62 ± 0.23 | 0.68 ± 0.18 | 0.47 ± 0.20 | 0.84 ± 0.13 | 0.47 ± 0.22 |
| Kermut | 0.67 ± 0.23 | 0.67 ± 0.19 | 0.51 ± 0.20 | **0.88 ± 0.14** | 0.50 ± 0.19 |
| ProteinNPT | 0.62 ± 0.24 | **0.69 ± 0.21** | 0.47 ± 0.22 | 0.86 ± 0.11 | 0.50 ± 0.20 |
| Cerebra-Epistasis | **0.69 ± 0.21** | 0.68 ± 0.17 | **0.53 ± 0.19** | **0.88 ± 0.11** | **0.52 ± 0.18** |
| **Order-averaged assay-level evaluation** |  |  |  |  |  |
| OHE | 0.59 ± 0.21 | 0.60 ± 0.18 | 0.44 ± 0.17 | 0.82 ± 0.13 | 0.43 ± 0.20 |
| OHE + ESM-1v | 0.59 ± 0.20 | 0.60 ± 0.18 | 0.44 ± 0.17 | 0.82 ± 0.13 | 0.43 ± 0.19 |
| Embeddings + ESM-1v | 0.55 ± 0.23 | 0.55 ± 0.20 | 0.41 ± 0.19 | 0.74 ± 0.20 | 0.40 ± 0.18 |
| OHE + Tranception | 0.59 ± 0.20 | 0.59 ± 0.18 | 0.44 ± 0.17 | 0.82 ± 0.12 | 0.43 ± 0.20 |
| Embeddings + Tranception | 0.57 ± 0.21 | 0.59 ± 0.16 | 0.42 ± 0.18 | 0.79 ± 0.13 | 0.41 ± 0.18 |
| OHE + MSA Transformer | 0.59 ± 0.20 | 0.60 ± 0.17 | 0.44 ± 0.16 | 0.81 ± 0.13 | 0.43 ± 0.19 |
| Embeddings + MSA Transformer | 0.56 ± 0.23 | 0.57 ± 0.19 | 0.42 ± 0.20 | 0.77 ± 0.15 | 0.41 ± 0.19 |
| MULTI-evolve | 0.60 ± 0.22 | **0.64 ± 0.18** | 0.46 ± 0.19 | 0.83 ± 0.13 | 0.47 ± 0.21 |
| Kermut | 0.60 ± 0.22 | 0.61 ± 0.19 | 0.45 ± 0.19 | 0.83 ± 0.14 | 0.44 ± 0.20 |
| ProteinNPT | 0.56 ± 0.21 | 0.62 ± 0.21 | 0.42 ± 0.19 | 0.80 ± 0.14 | 0.43 ± 0.22 |
| Cerebra-Epistasis | **0.63 ± 0.21** | 0.63 ± 0.18 | **0.48 ± 0.18** | **0.85 ± 0.12** | **0.50 ± 0.17** |

### S5.4 Supplementary Results for the cDNA proteolysis multi-mutant dataset in the 1M+ to 1M+ benchmark

Results for the cDNA proteolysis multi-mutant benchmark described in Supplementary Section S2.6 are summarized in Supplementary Table S8. Cerebra-Epistasis achieved the best performance on most metrics.

**Table S8.** Performance on the cDNA proteolysis multi-mutant dataset of the 1M+ to 1M+ benchmark. Results are reported using overall assay-level evaluation and order-averaged assay-level evaluation as mean ± standard deviation.

| Model | Spearman $\rho$ | Pearson $r$ | Kendall $\tau$ | NDCG<br>(top 10%) | Top recall<br>(top 10%) |
| --- | --- | --- | --- | --- | --- |
| **Overall assay-level evaluation** |  |  |  |  |  |
| OHE | 0.79 ± 0.06 | 0.75 ± 0.07 | 0.61 ± 0.07 | 0.89 ± 0.05 | 0.44 ± 0.12 |
| OHE + ESM-1v | 0.79 ± 0.08 | 0.79 ± 0.08 | 0.60 ± 0.08 | 0.89 ± 0.04 | 0.41 ± 0.11 |
| Embeddings + ESM-1v | 0.89 ± 0.05 | 0.89 ± 0.05 | 0.71 ± 0.06 | 0.94 ± 0.02 | 0.55 ± 0.10 |
| OHE + Tranception | 0.81 ± 0.07 | 0.81 ± 0.07 | 0.63 ± 0.07 | 0.90 ± 0.04 | 0.41 ± 0.11 |
| Embeddings + Tranception | 0.90 ± 0.05 | 0.90 ± 0.05 | 0.73 ± 0.06 | 0.94 ± 0.02 | 0.59 ± 0.09 |
| OHE + MSA Transformer | 0.82 ± 0.07 | 0.82 ± 0.07 | 0.64 ± 0.07 | 0.90 ± 0.04 | 0.42 ± 0.12 |
| Embeddings + MSA Transformer | 0.92 ± 0.04 | 0.92 ± 0.04 | 0.76 ± 0.05 | 0.95 ± 0.02 | 0.62 ± 0.10 |
| Kermut | 0.94 ± 0.03 | 0.94 ± 0.03 | 0.79 ± 0.05 | 0.96 ± 0.01 | 0.68 ± 0.09 |
| ProteinNPT | 0.94 ± 0.03 | 0.94 ± 0.03 | 0.81 ± 0.05 | **0.97 ± 0.02** | **0.71 ± 0.08** |
| Cerebra-Epistasis | **0.94 ± 0.03** | **0.94 ± 0.04** | **0.81 ± 0.05** | **0.97 ± 0.01** | 0.70 ± 0.08 |
| **Order-averaged assay-level evaluation** |  |  |  |  |  |
| OHE | 0.76 ± 0.09 | 0.73 ± 0.09 | 0.58 ± 0.09 | 0.89 ± 0.06 | 0.49 ± 0.09 |
| OHE + ESM-1v | 0.73 ± 0.09 | 0.75 ± 0.09 | 0.56 ± 0.08 | 0.89 ± 0.05 | 0.47 ± 0.08 |
| Embeddings + ESM-1v | 0.83 ± 0.08 | 0.85 ± 0.08 | 0.65 ± 0.08 | 0.93 ± 0.03 | 0.56 ± 0.09 |
| OHE + Tranception | 0.76 ± 0.08 | 0.78 ± 0.09 | 0.59 ± 0.08 | 0.90 ± 0.05 | 0.49 ± 0.08 |
| Embeddings + Tranception | 0.85 ± 0.08 | 0.86 ± 0.08 | 0.67 ± 0.08 | 0.94 ± 0.03 | 0.60 ± 0.08 |
| OHE + MSA Transformer | 0.78 ± 0.08 | 0.79 ± 0.09 | 0.60 ± 0.08 | 0.90 ± 0.05 | 0.50 ± 0.08 |
| Embeddings + MSA Transformer | 0.87 ± 0.07 | 0.88 ± 0.07 | 0.70 ± 0.08 | 0.94 ± 0.03 | 0.62 ± 0.08 |
| Kermut | 0.89 ± 0.07 | 0.91 ± 0.07 | 0.74 ± 0.08 | 0.96 ± 0.03 | 0.68 ± 0.08 |
| ProteinNPT | 0.90 ± 0.07 | 0.91 ± 0.07 | **0.76 ± 0.08** | **0.96 ± 0.03** | **0.71 ± 0.08** |
| Cerebra-Epistasis | **0.91 ± 0.07** | **0.92 ± 0.07** | **0.76 ± 0.08** | 0.96 ± 0.03 | 0.69 ± 0.08 |

### S5.5 Supplementary Results for the avGFP benchmark

Results for the avGFP experimental fitness prediction benchmark described in Supplementary Section S2.7 are summarized in Supplementary Table S9. Cerebra-Epistasis achieved the highest Spearman correlation (0.83) and Kendall correlation (0.67) among the compared methods, while attaining competitive performance on the remaining metrics, including an NDCG (top 5) of 0.67 and a TopRecall (top 5) of 0.40.

**Table S9.** Performance on the avGFP experimental fitness prediction benchmark. The best result for each metric is shown in bold.

| Model | Spearman $\rho$ | Pearson $r$ | Kendall $\tau$ | NDCG<br>(top 5) | Top recall<br>(top 5) |
| --- | --- | --- | --- | --- | --- |
| OHE | 0.47 | 0.40 | 0.33 | 0.48 | 0.20 |
| OHE + ESM-1v | 0.56 | 0.44 | 0.38 | 0.37 | 0.20 |
| Embeddings + ESM-1v | 0.51 | 0.40 | 0.35 | 0.42 | 0.20 |
| OHE + Tranception | 0.73 | 0.52 | 0.56 | **0.73** | **0.60** |
| Embeddings + Tranception | 0.66 | 0.52 | 0.48 | 0.43 | 0.00 |
| OHE + MSA Transformer | 0.74 | **0.57** | 0.57 | 0.62 | 0.40 |
| Embeddings + MSA Transformer | 0.70 | 0.53 | 0.54 | 0.50 | **0.60** |
| MULTI-evolve | 0.54 | 0.22 | 0.38 | 0.64 | 0.40 |
| ProteinNPT | 0.73 | 0.51 | 0.58 | 0.62 | 0.20 |
| Cerebra-Epistasis | **0.83** | 0.55 | **0.67** | 0.67 | 0.40 |

### S5.6 Supplementary Results for Low-*N* 1M/2M to held-out 2M (epistasis)

Results for the Low-*N* 1M/2M to held-out 2M (epistasis) benchmark described in Supplementary Section S2.8 are summarized in Supplementary Table S10. Cerebra-Epistasis showed strong performance across both correlation- and ranking-based metrics for predicting epistatic effects of unseen double-mutant combinations.

**Table S10.** Assay-level performance on the Low-*N* 1M/2M to held-out 2M (epistasis) benchmark. Values are reported as mean ± SD across assays. The best result for each metric is shown in bold.

| Model | Spearman $\rho$ | Pearson $r$ | Kendall $\tau$ | NDCG<br>(top 10%) | Top recall<br>(top 10%) |
| --- | --- | --- | --- | --- | --- |
| OHE | 0.00 ± 0.10 | 0.02 ± 0.12 | 0.00 ± 0.07 | 0.62 ± 0.12 | 0.12 ± 0.07 |
| OHE + ESM-1v | -0.07 ± 0.11 | -0.06 ± 0.12 | -0.04 ± 0.07 | 0.59 ± 0.14 | 0.08 ± 0.06 |
| Embeddings + ESM-1v | 0.19 ± 0.21 | 0.23 ± 0.22 | 0.13 ± 0.15 | 0.68 ± 0.14 | 0.23 ± 0.15 |
| OHE + Tranception | -0.05 ± 0.13 | -0.07 ± 0.15 | -0.04 ± 0.09 | 0.62 ± 0.13 | 0.11 ± 0.07 |
| Embeddings + Tranception | 0.37 ± 0.26 | 0.43 ± 0.28 | 0.26 ± 0.19 | 0.77 ± 0.13 | 0.34 ± 0.22 |
| OHE + MSA Transformer | -0.06 ± 0.09 | -0.09 ± 0.10 | -0.04 ± 0.07 | 0.59 ± 0.13 | 0.09 ± 0.06 |
| Embeddings + MSA Transformer | 0.23 ± 0.28 | 0.24 ± 0.28 | 0.16 ± 0.20 | 0.68 ± 0.15 | 0.20 ± 0.16 |
| MULTI-evolve | 0.19 ± 0.20 | 0.29 ± 0.29 | 0.13 ± 0.14 | 0.78 ± 0.12 | 0.33 ± 0.27 |
| Kermut | 0.35 ± 0.26 | 0.39 ± 0.28 | 0.25 ± 0.19 | 0.79 ± 0.12 | 0.34 ± 0.23 |
| Cerebra-Epistasis | **0.55 ± 0.25** | **0.60 ± 0.26** | **0.41 ± 0.20** | **0.84 ± 0.11** | **0.47 ± 0.24** |

### S5.7 Effect of end-to-end training on ΔΔ*G* prediction

To assess whether end-to-end training improves downstream fitness prediction, we focused this comparison on protein stability prediction, because large-scale cDNA proteolysis data support training across diverse proteins, while established ΔΔ*G* benchmarks provide standardized external evaluation.

We compared a downstream-only setting, in which the pretrained Cerebra-Seq module was frozen, with an end-to-end setting, in which the Cerebra-Seq module and downstream prediction network were trained end to end. The two model variants were evaluated alongside established ΔΔ*G* prediction methods on the S461 benchmark using the evaluation protocol described in Supplementary Section S2.9. The results are shown in Supplementary Table S11.

**Table S11.** Comparison of ΔΔ*G* prediction performance on the S461 benchmark. Cerebra-Epistasis^∗^ denotes end-to-end training of Cerebra-Seq and the downstream predictor on the complete cDNA proteolysis dataset; Cerebra-Epistasis denotes the corresponding setting in which Cerebra-Seq was frozen and only the downstream predictor was optimized. Values are the mean ± standard deviation across proteins. Bold values indicate the best mean performance for each metric, including ties.

| Model | Spearman $\rho$ | Kendall $\tau$ | Pearson $r$ | Top-5 Precision | Top-10 Precision |
| --- | --- | --- | --- | --- | --- |
| DDMut | 0.48 ± 0.40 | 0.37 ± 0.31 | 0.52 ± 0.39 | 0.45 ± 0.26 | 0.61 ± 0.18 |
| ProteinEBM | 0.51 ± 0.22 | 0.38 ± 0.16 | 0.53 ± 0.26 | 0.47 ± 0.23 | 0.67 ± 0.19 |
| PROSTATA | 0.53 ± 0.33 | 0.40 ± 0.25 | 0.55 ± 0.34 | 0.53 ± 0.25 | 0.67 ± 0.23 |
| RaSP | 0.56 ± 0.30 | 0.43 ± 0.23 | 0.54 ± 0.28 | 0.47 ± 0.25 | 0.64 ± 0.22 |
| SPIRED-Fitness (zero-shot) | 0.58 ± 0.33 | 0.45 ± 0.25 | 0.59 ± 0.30 | 0.58 ± 0.16 | 0.70 ± 0.17 |
| GeoDDG-3D | 0.59 ± 0.23 | 0.44 ± 0.20 | 0.59 ± 0.23 | **0.63 ± 0.17** | 0.72 ± 0.15 |
| GeoDDG-Seq | 0.60 ± 0.23 | 0.46 ± 0.18 | 0.63 ± 0.20 | 0.55 ± 0.19 | 0.72 ± 0.17 |
| Mutate Everything (ESM-2) | 0.63 ± 0.19 | 0.46 ± 0.16 | 0.62 ± 0.18 | 0.45 ± 0.24 | 0.69 ± 0.18 |
| ThermoMPNN | 0.63 ± 0.32 | 0.48 ± 0.26 | 0.60 ± 0.31 | 0.48 ± 0.23 | 0.67 ± 0.21 |
| Pythia | 0.64 ± 0.19 | 0.48 ± 0.16 | 0.63 ± 0.18 | 0.57 ± 0.22 | 0.68 ± 0.17 |
| SPURS | 0.69 ± 0.25 | 0.55 ± 0.23 | 0.70 ± 0.23 | 0.62 ± 0.26 | 0.73 ± 0.14 |
| SPIRED-Stab | **0.71 ± 0.24** | **0.57 ± 0.21** | 0.69 ± 0.21 | 0.62 ± 0.22 | 0.68 ± 0.24 |
| Cerebra-Epistasis | 0.61 ± 0.32 | 0.48 ± 0.28 | 0.66 ± 0.30 | 0.52 ± 0.26 | 0.67 ± 0.21 |
| Cerebra-Epistasis* | **0.71 ± 0.24** | 0.57 ± 0.20 | **0.71 ± 0.22** | 0.57 ± 0.14 | **0.74 ± 0.17** |

## S6 Ablation and sensitivity analyses

We performed a series of ablation and sensitivity analyses to examine two key design principles underlying Cerebra-Epistasis: structure-aware representation learning and explicit modeling of epistatic interactions across mutation orders. We first assessed how Cerebra-Seq-derived structure-aware features contribute to higher-order extrapolation and whether explicit geometric information helps recover the spatial organization of epistasis. We then examined the contribution of explicit epistasis modeling and investigated how the mutation orders and pairwise interaction coverage represented during training affect generalization to unseen higher-order combinations. Finally, we evaluated whether end-to-end optimization of the structure and fitness prediction modules further improves mutation-effect prediction.

### S6.1 Contribution of Cerebra-Seq-derived structure-aware features to higher-order extrapolation

To assess whether structure-aware information improves higher-order extrapolation beyond sequence representations alone, we progressively augmented the ESM-2-derived features with three-dimensional coordinates, Cerebra residue-level node representations, and pairwise edge representations. This progression introduces increasingly rich geometric and structural context, from explicit residue coordinates to learned residue- and pair-level representations derived by Cerebra. On the 1M/2M to 3M+ benchmark, progressively incorporating these features generally improved both overall performance and performance across mutation orders (Supplementary Fig S2). These results indicate that Cerebra-Seq-derived structural information provides complementary signals beyond sequence embeddings for predicting unseen higher-order mutant combinations.

**Figure S2.**
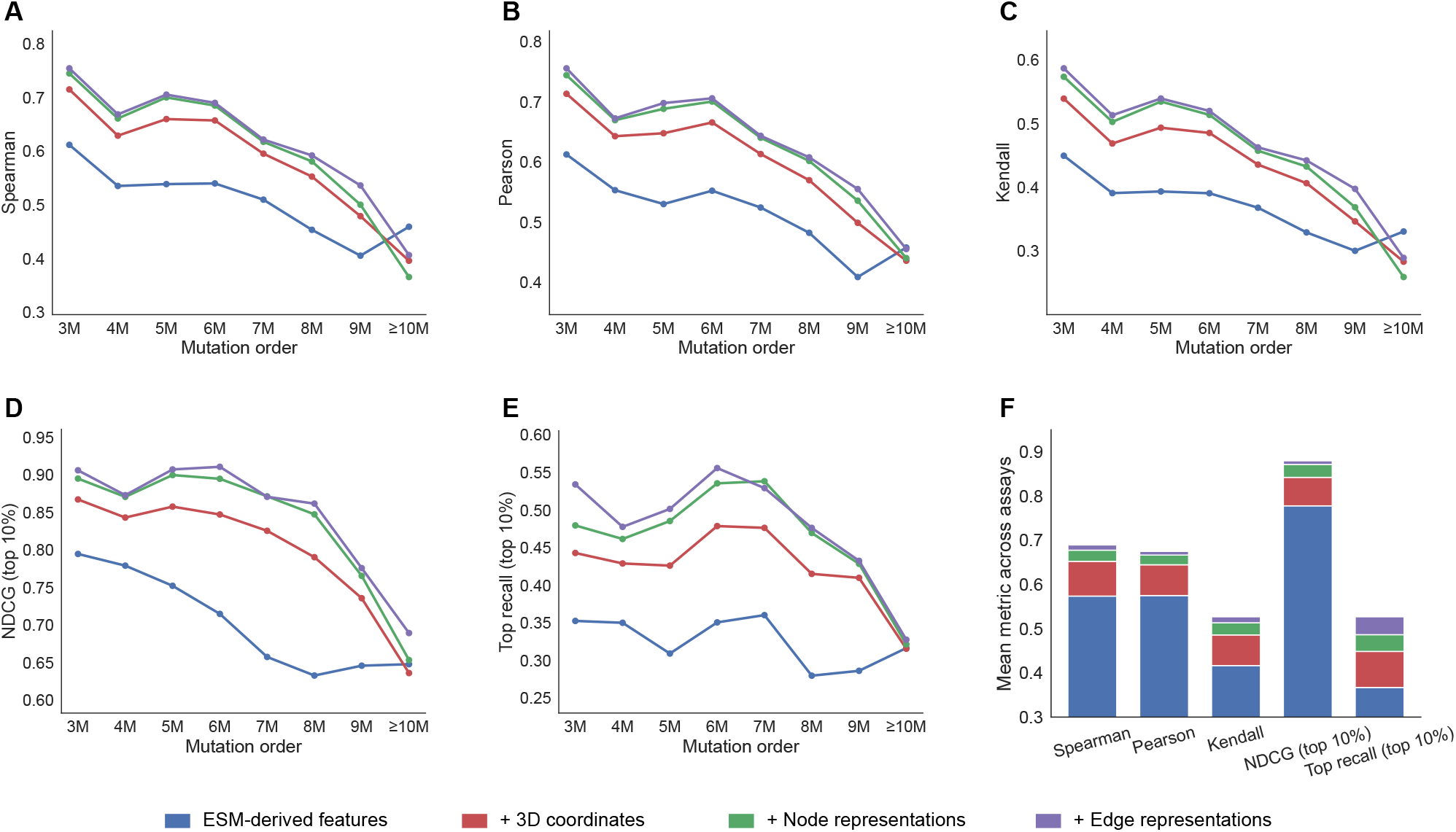
Contribution of Cerebra-Seq-derived structure-aware features to higher-order extrapolation. Models were trained on single- and double-mutant variants and evaluated on held-out variants containing three or more mutations. Starting from ESM-2-derived sequence features, three-dimensional coordinates, Cerebra residue-level node representations, and pairwise edge representations were progressively incorporated. **A–E**, Spearman, Pearson, Kendall, NDCG (top 10%), and Top recall (top 10%) across mutation orders. **F**, Assay-averaged performance across the five evaluation metrics, summarizing the contribution of progressively added structure-aware features.

### S6.2 Contribution of explicit epistasis modeling to higher-order extrapolation

To determine whether higher-order extrapolation in the 1M/2M to 3M+ benchmark benefits from explicit epistasis modeling, we compared the complete Cerebra-Epistasis model with an additive-only model. Both models were trained on the same single- and double-mutant data and evaluated on the same held-out 3M+ variants. The additive-only model predicted multi-mutant fitness as the sum of its constituent single-mutation contributions, whereas the complete model combined these additive contributions with an explicit epistatic term. The complete model outperformed the additive-only model in overall performance and generally maintained stronger performance across mutation orders (Supplementary Fig S3), demonstrating that modeling non-additive interactions explicitly is important for transferring epistatic information from lower-order measurements to unseen higher-order mutant combinations.

**Figure S3.**
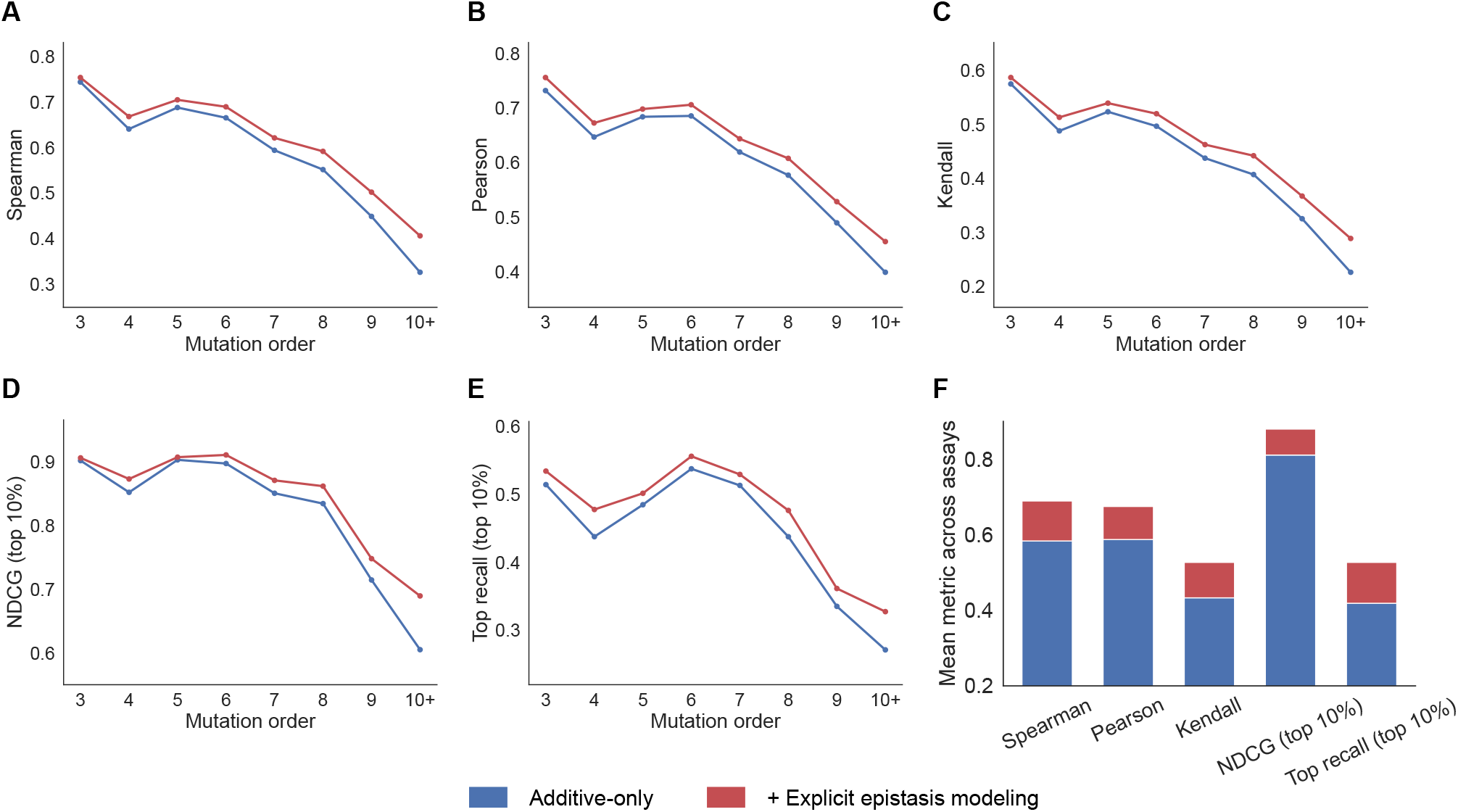
Effect of explicit epistasis modeling on higher-order fitness extrapolation. Comparison of Cerebra-Epistasis and the additive-only model on the 1M/2M to 3M+ benchmark. **A–E**, Assay-averaged Spearman, Pearson, Kendall, NDCG (top 10%), and top recall (top 10%) at each mutation order. **F**, Overall assay-level performance across held-out higher-order variants, with the incremental gain over the additive-only model indicating the contribution of explicit epistasis modeling.

### S6.3 Effect of maximum training mutation order on higher-order extrapolation

To examine how the maximum mutation order included in the training data affects extrapolation to more complex variants, models were trained on progressively expanded mutation-order ranges and evaluated on the same higher-order test variants (Supplementary Fig S4A). Adding double mutants to the single-mutant training set produced a marked improvement in higher-order prediction, indicating that pairwise measurements provide important information about non-additive interactions that is difficult to infer from single-mutant effects alone. Including variants with progressively higher mutation orders yielded further improvements, although these gains were generally smaller than the improvement obtained from introducing double mutants. These results suggest that pairwise epistasis provides much of the interaction information needed for higher-order extrapolation, whereas higher-order training examples contribute additional information about more complex mutational dependencies.

### S6.4 Effect of double-mutant coverage on higher-order extrapolation

To investigate how extensively pairwise interactions must be sampled before reliable higher-order extrapolation becomes possible, the single-mutant training set and higher-order test set were held fixed, while the proportion of available double mutants used for training was varied (Supplementary Fig S4B). Prediction performance improved as double-mutant coverage increased, confirming that direct observations of pairwise epistasis provide transferable information for predicting unseen higher-order combinations. Notably, substantial gains were already obtained from partial double-mutant coverage, suggesting that the model can infer useful interaction patterns without exhaustively measuring the complete pairwise landscape. The progressively smaller improvements at higher coverage further indicate diminishing returns, with additional double-mutant measurements mainly refining an interaction representation that is already partially established from a limited subset of pairwise observations.

**Figure S4.**
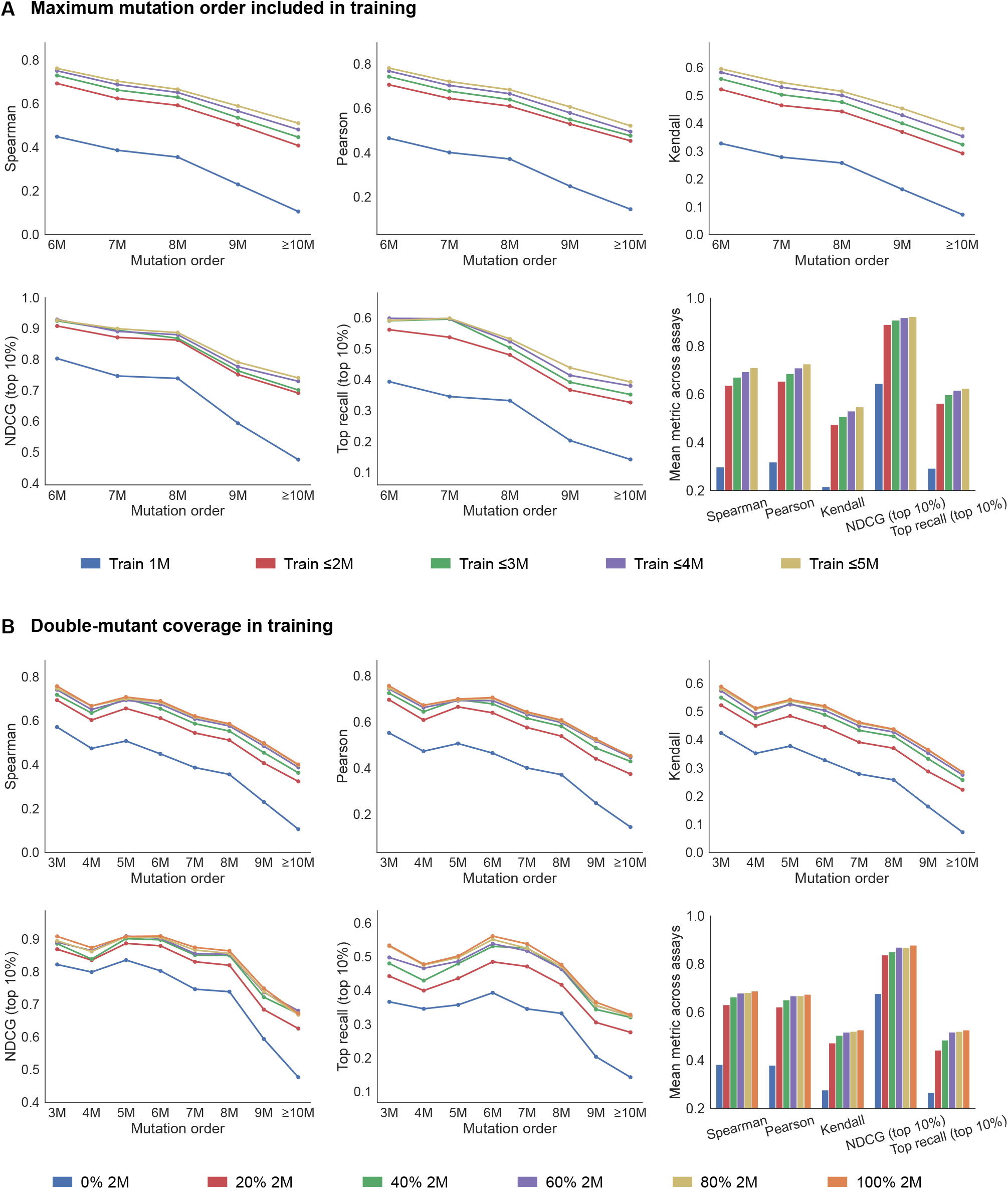
Effects of training mutation order and double-mutant coverage on higher-order extrapolation. **A**, Performance of models trained with progressively increasing maximum mutation orders. **B**, Performance of models trained with different proportions of the available double mutants. For each ablation, the five mutation-order plots show assay-averaged Spearman, Pearson, Kendall, NDCG (top 10%), and top recall (top 10%), while the final plot shows overall assay-level performance across held-out 3M+ variants.

### S6.5 Effect of end-to-end training on fitness and structure prediction

To distinguish the contributions of fitness gradients and structural supervision, we compared four settings on identical sequence-grouped five-fold partitions of the cDNA proteolysis multi-mutant dataset. Fitness gradients updated the downstream components in every setting, with Cerebra-Seq kept fixed in the frozen baseline. In structure-only training, only structural gradients updated Cerebra-Seq, while fitness gradients were restricted to the downstream components. Fitness-only training propagated fitness gradients through both modules without structural supervision. End-to-end training (Structure + Fitness) allowed both fitness and structural gradients to update Cerebra-Seq. These comparisons also tested whether adding structural supervision improves fitness prediction when fitness gradients already update the upstream model. Across epochs 1–100, all three update settings improved trajectory-averaged Spearman correlations over frozen Cerebra-Seq. End-to-end training achieved the highest correlation, exceeding structure-only training and modestly improving on fitness-only training (Supplementary Fig S5A,B). Fitness-only training progressively increased side-chain *χ* torsion-angle loss and reduced C_*α*_ lDDT. End-to-end training instead reduced torsion-angle loss below the pretrained baseline and largely preserved lDDT (Supplementary Fig S5C,D). Its trajectory-averaged lDDT remained slightly below that of structure-only training. These results indicate that fitness gradients improve prediction beyond structural supervision alone, while additional structural supervision further improves prediction and helps preserve structural accuracy.

**Figure S5.**
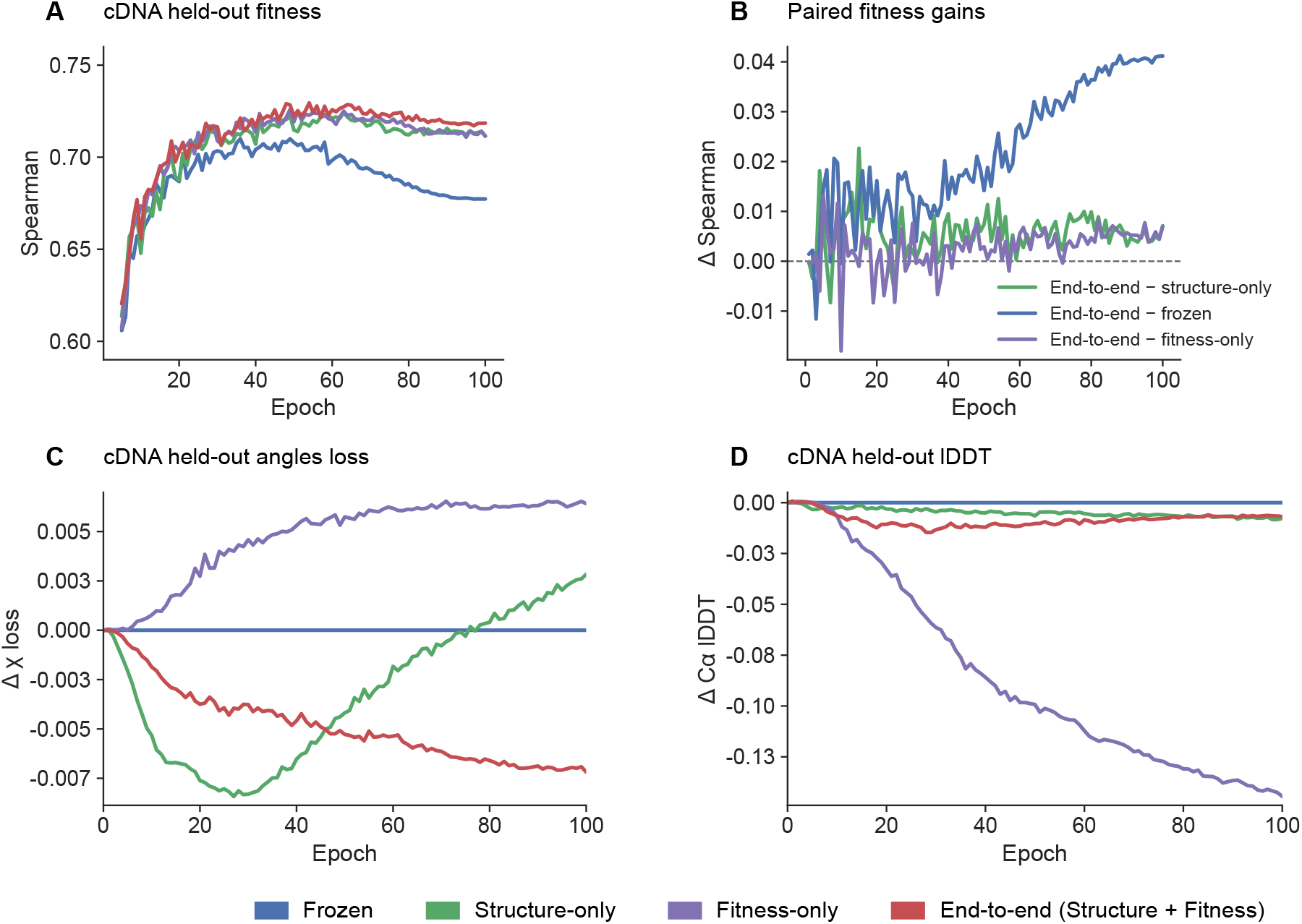
Contributions of fitness gradients and structural supervision to end-to-end training. Models used identical sequence-grouped five-fold partitions of the cDNA proteolysis multi-mutant dataset, with fitness gradients updating the downstream components in every setting. In **A, C** and **D**, blue denotes frozen Cerebra-Seq; green, structure-only training; purple, fitness-only training; and red, end-to-end training (Structure + Fitness). Only structural gradients updated Cerebra-Seq in structure-only training, only fitness gradients in fitness-only training, and both in end-to-end training. **A**, Spearman correlations for held-out proteins. **B**, Paired gains in Spearman correlation from end-to-end training over structure-only training (green), frozen Cerebra-Seq (blue), and fitness-only training (purple). Differences were calculated within the same fold and epoch before averaging. **C**, Changes in side-chain *χ* torsion-angle loss from the pretrained baseline; negative values indicate improvement. **D**, Changes in C_*α*_ lDDT from the pretrained baseline; positive values indicate improvement. Zero denotes the fold-specific pretrained baseline in **C** and **D**. Lines show equal-weight means across five folds.

## S7 Cerebra-Epistasis Inference

This section presents pseudocode for the complete inference workflow of Cerebra-Epistasis and its main components. Supplementary Algorithm S1 first provides a high-level overview of the framework, encompassing one-time wild-type encoding by Cerebra-Seq, structure-aware mutation-atlas construction, and fitness and epistasis prediction for arbitrary-order mutants. The subsequent algorithms describe the Cerebra-Seq structure predictor, the downstream fitness prediction modules, and the core SE(3)-equivariant operations in detail.

### S7.1 Features and Notation

Supplementary Table S12 summarizes the principal features and notation shared across the algorithms. Shapes of short-lived intermediate variables are given locally in the rightmost column of each algorithm.

**Table S12.** Features and notation used in the pseudocode algorithms.

| Feature | Description | Shape |
| --- | --- | --- |
| $L$ | Number of residues in the wild-type protein | |
| $K$ | Number of anchor-conditioned structure predictions | |
| $B$ | Number of candidate mutants evaluated together | |
| $m$ | Number of substitutions in an individual mutant | |
| $r_H$ | Dimension of the learned epistasis representation | |
| **wt_seq** | Wild-type amino-acid sequence | $[L]$ |
| $\mathbf{X}^{\text{ESM-2}}$ | Per-residue ESM-2 representation | $[L, 1280]$ |
| $\mathbf{X}^{\text{ESMC}}$ | Per-residue ESMC representation | $[L, 1152]$ |
| $\mathbf{X}^{\text{ESM3}}$ | Per-residue ESM3 representation | $[L, 1536]$ |
| **m** | Cerebra-Seq single representation arranged over ten streams | $[10, L, 256]$ |
| **z** | Cerebra-Seq residue-pair representation | $[L, L, 128]$ |
| **T** | Anchor-conditioned residue translations | $[K, L, 3]$ |
| **Q** | Anchor-conditioned residue orientations represented as quaternions | $[K, L, 4]$ |
| **N** | Cerebra-Seq residue-level structural representation | $[L, 256]$ |
| **E** | Cerebra-Seq residue-pair structural representation | $[L, L, 128]$ |
| $\mathcal{F}$ | Collection of degree-wise SE(3) features, $\{\mathbf{F}^{(0)}, \mathbf{F}^{(1)}\}$ | $[L, C_0, 1]; [L, C_1, 3]$ |
| $\tilde{\mathbf{E}}$ | Pair representation projected to the SE(3) edge channels | $[L, L, 32]$ |
| **Z** | Degree-0 invariant output representation of the SE(3) encoder | $[L, 128]$ |
| **V** | Degree-1 equivariant output representation of the SE(3) encoder | $[L, 32, 3]$ |
| **D<sup>stat</sup>** | Mean and standard deviation of the selected neighborhood distances | $[L, 2]$ |
| **S** | WT-centered single-mutation score tensor | $[L, 20]$ |
| **U** | WT-centered epistasis representation tensor | $[L, 20, r_H]$ |
| $\mathcal{M}$ | Individual mutant represented as $\{(i_k, a_k)\}_{k=1}^m$ | $[m, 2]$ |
| $\mathcal{Q}$ | Collection of candidate mutants, $\{\mathcal{M}_b\}_{b=1}^B$ | $[B]$ |
| $\hat{y}_{\mathcal{M}}$ | Predicted fitness of mutant $\mathcal{M}$ | |
| $\hat{\epsilon}_{\mathcal{M}}$ | Predicted total epistatic contribution of mutant $\mathcal{M}$ | |

### S7.2 Overall Cerebra-Epistasis workflow

Given a wild-type sequence and a collection of candidate mutants, Cerebra-Epistasis first runs Cerebra-Seq once to obtain atom14 coordinates together with residue-level node and residue-pair edge representations. The downstream encoder then converts these shared wild-type features into the single-mutation score tensor **S** and epistasis representation tensor **U**. Each candidate mutant is subsequently evaluated by lightweight indexing and permutation-invariant assembly, without repeating sequence or structure encoding.

#### Algorithm S1

Complete Cerebra-Epistasis workflow

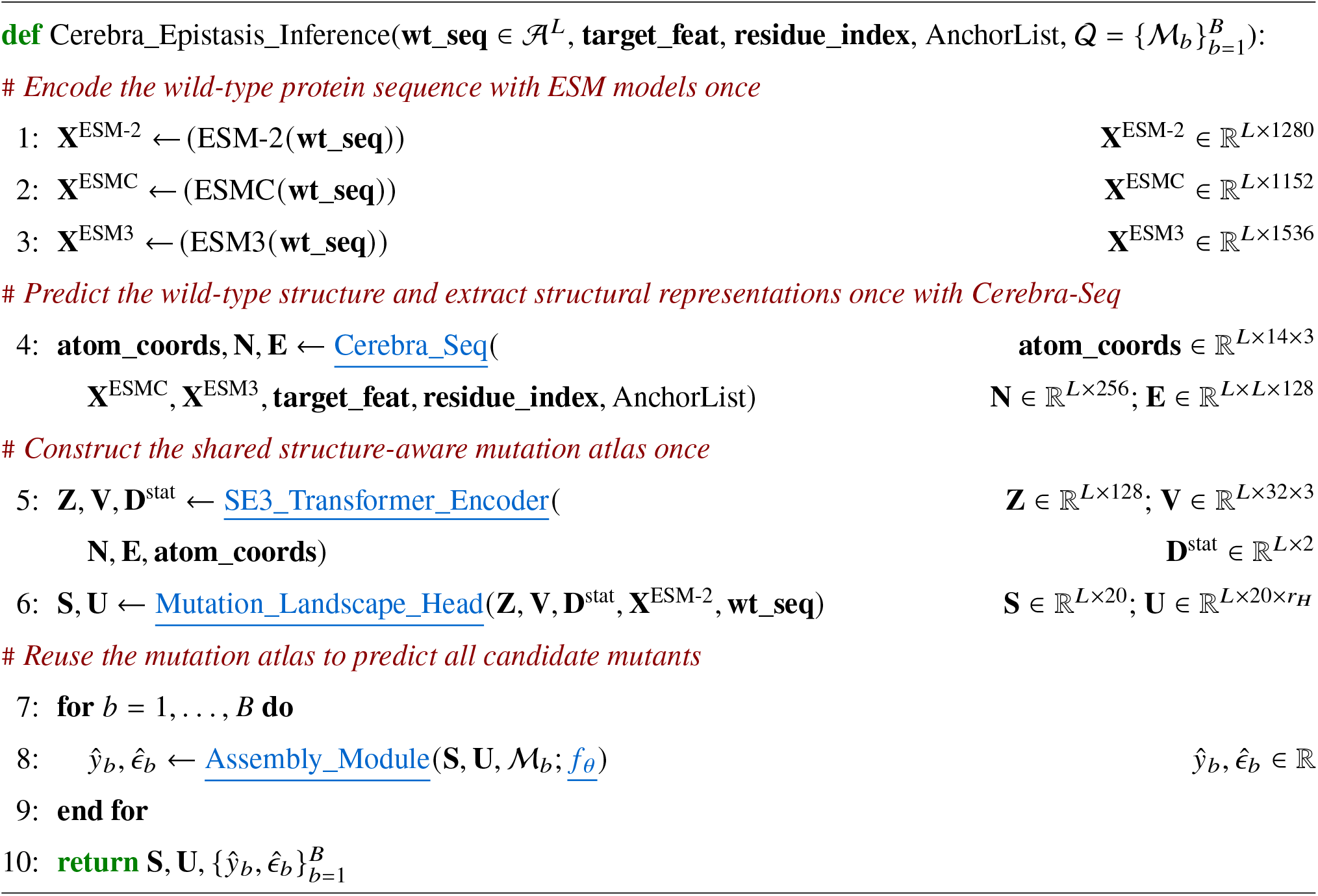

### S7.3 Cerebra-Seq structure prediction

Supplementary Algorithm S2 and Supplementary Algorithm S3 summarize the Cerebra-Seq forward pass using the module-oriented notation of the original Cerebra description [18]. Supplementary Algorithm S2 presents inference for one Cerebra-Seq checkpoint. Frozen ESMC and ESM3 representations are extracted once and reused across four prediction cycles, consisting of an initial pass and three recycling updates with shared network weights. In each cycle, the single and pair representations are initialized together with the recycled states, refined by the 32-block Evoformer Module, and passed through four sequential Cerebra StructureGenerationMotifs. Each motif retains the original Cerebra ordering of the 1D encoder, structure encoder, 1D decoder, structure decoder, and PSA operations. After the final cycle, the anchor-conditioned residue frames are aligned and combined, and the resulting frames and seven torsion-angle representations are converted to atom14 coordinates. The next algorithm isolates the ESMC–ESM3 fusion and mHC-inspired mixing used in the input pathway. Unless otherwise specified, modules not detailed here retain the architectures described in Cerebra [18].

#### Algorithm S2

Cerebra-Seq model inference

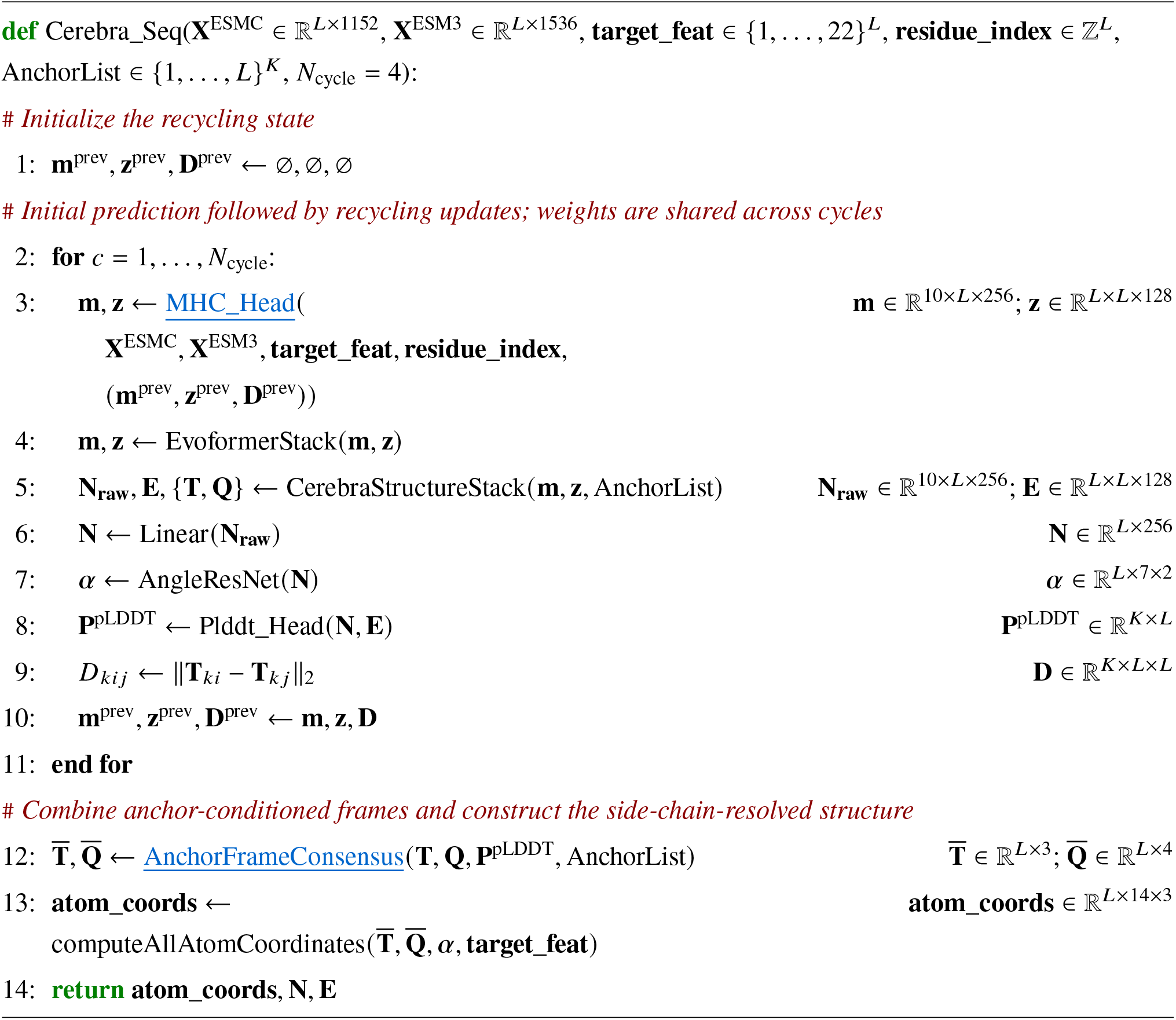

Supplementary Algorithm S3 details the component that distinguishes the Cerebra-Seq input pathway from the MSA-based Cerebra Embedding Module. ESMC and ESM3 representations are concatenated residue-wise, projected to 2,560 channels, and partitioned into ten streams of width 256. A shared learnable logit matrix is converted in log space to a doubly stochastic matrix by 20 Sinkhorn iterations and then mixes the ten streams. The mixed language-model representation is combined with the target-residue embedding, while the 128-channel pair representation is initialized from target-residue and relative-position embeddings. From the second prediction cycle onward, the normalized single and pair states and the binned structural distances from the preceding cycle are added to these initial representations.

#### Algorithm S3

MHCMixingEmbedder Head

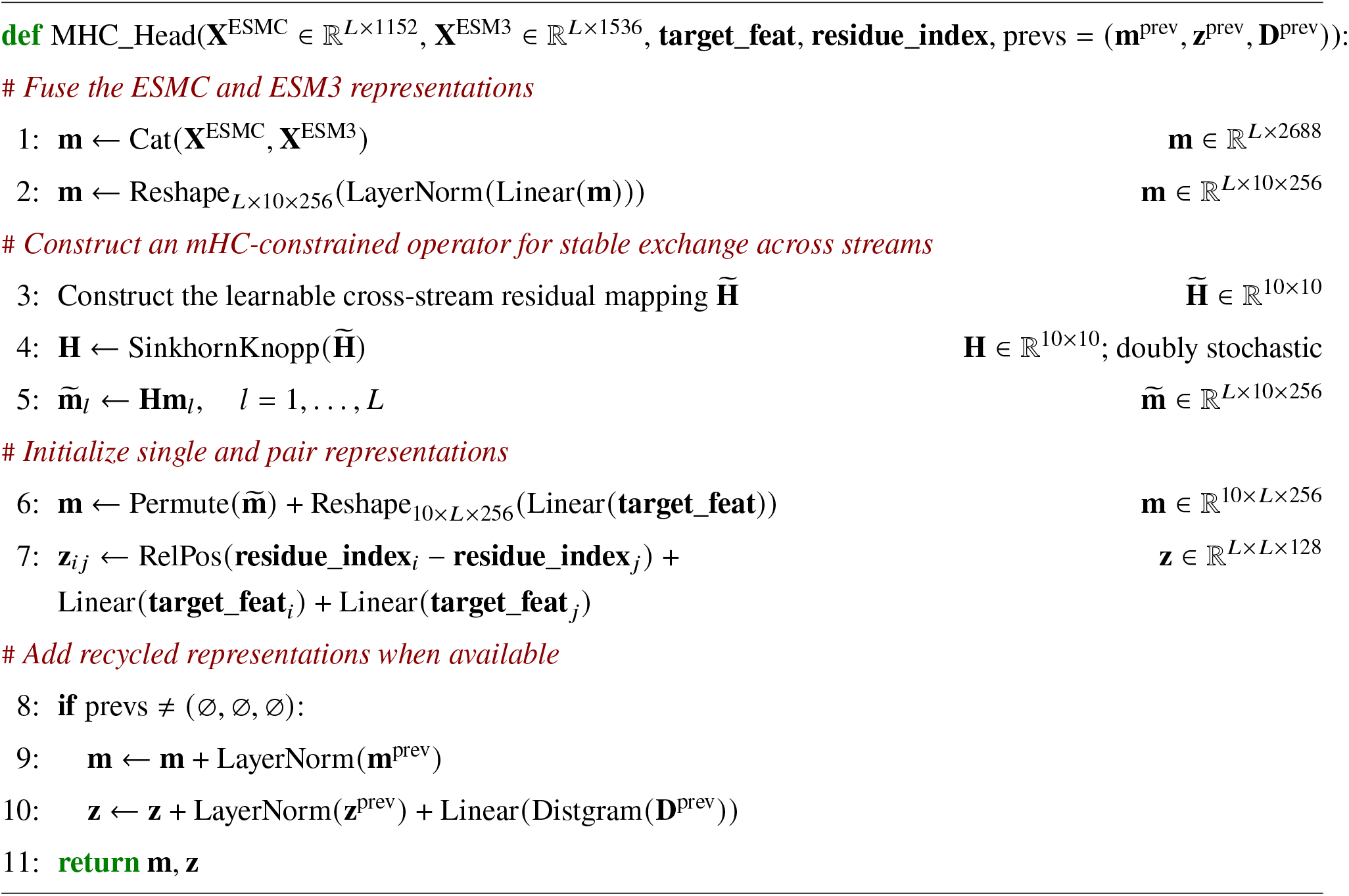

Supplementary Algorithm S4 summarizes how the anchor-conditioned predictions are combined. The prediction associated with the middle anchor is used as the reference. Each remaining prediction is rotated into this reference coordinate system. For every residue, the five aligned predictions with the highest pLDDT scores are then averaged to obtain one complete set of residue frames.

#### Algorithm S4

Consensus of anchor-conditioned residue frames

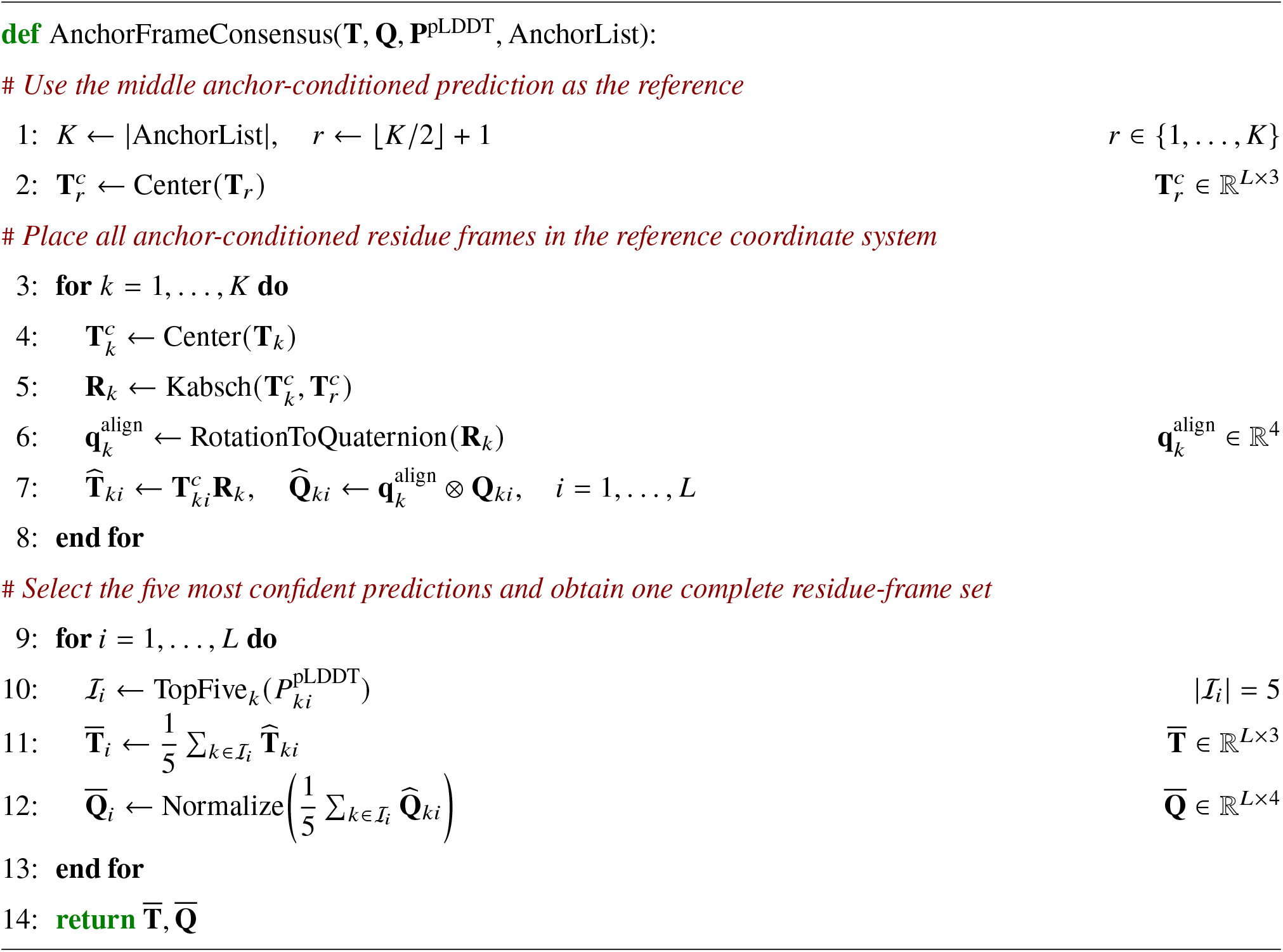

### S7.4 Downstream fitness prediction

The following algorithms expand the downstream part of Supplementary Algorithm S1. They describe the one-time encoding of the Cerebra-Seq representations, construction of the mutation atlas, and assembly of arbitrary-order mutant fitness and epistasis predictions.

#### S7.4.1 Downstream model workflow

Following the complete workflow in Supplementary Algorithm S1, the downstream model integrates the Cerebra-Seq residue, pairwise, and coordinate features of the wild-type protein using the SE(3)-Transformer encoder. The encoder integrates sequence and structural information into a degree-0 invariant representation **Z** and a degree-1 equivariant representation **V**, while also summarizing the selected neighborhood distances as local distance statistics **D**^stat^. These outputs, together with the ESM-2 650M embedding, are used to construct the single-mutation score tensor **S** ∈ ℝ^*L*×20^ and the epistasis representation tensor 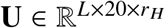. The entries corresponding to an arbitrary-order mutant are then indexed and assembled to predict its fitness and epistatic component.

#### S7.4.2 SE(3)-Transformer encoder

Supplementary Algorithm S5 describes the SE(3)-Transformer encoder used to integrate the Cerebra-Seq-derived residue, pairwise, and coordinate features of the wild-type protein. The residue features are projected into degree-0 invariant features, the atom coordinates initialize degree-1 equivariant features, and the pairwise features are projected into edge features. Geometric and pairwise-representation neighborhoods are then constructed to define the residue graph. The resulting features are processed by an input equivariant convolution, a single SE(3)-Transformer block, and an output equivariant convolution, yielding the degree-0 invariant representation **Z** ∈ ℝ^*L*×128^ and the degree-1 equivariant representation **V** ∈ ℝ^*L*×32×3^. The distances within the selected neighborhoods are additionally summarized by their residue-wise mean and standard deviation, producing the local distance statistics **D**^stat^ ∈ ℝ^*L*×2^.

##### Algorithm S5

SE(3)-Transformer encoder

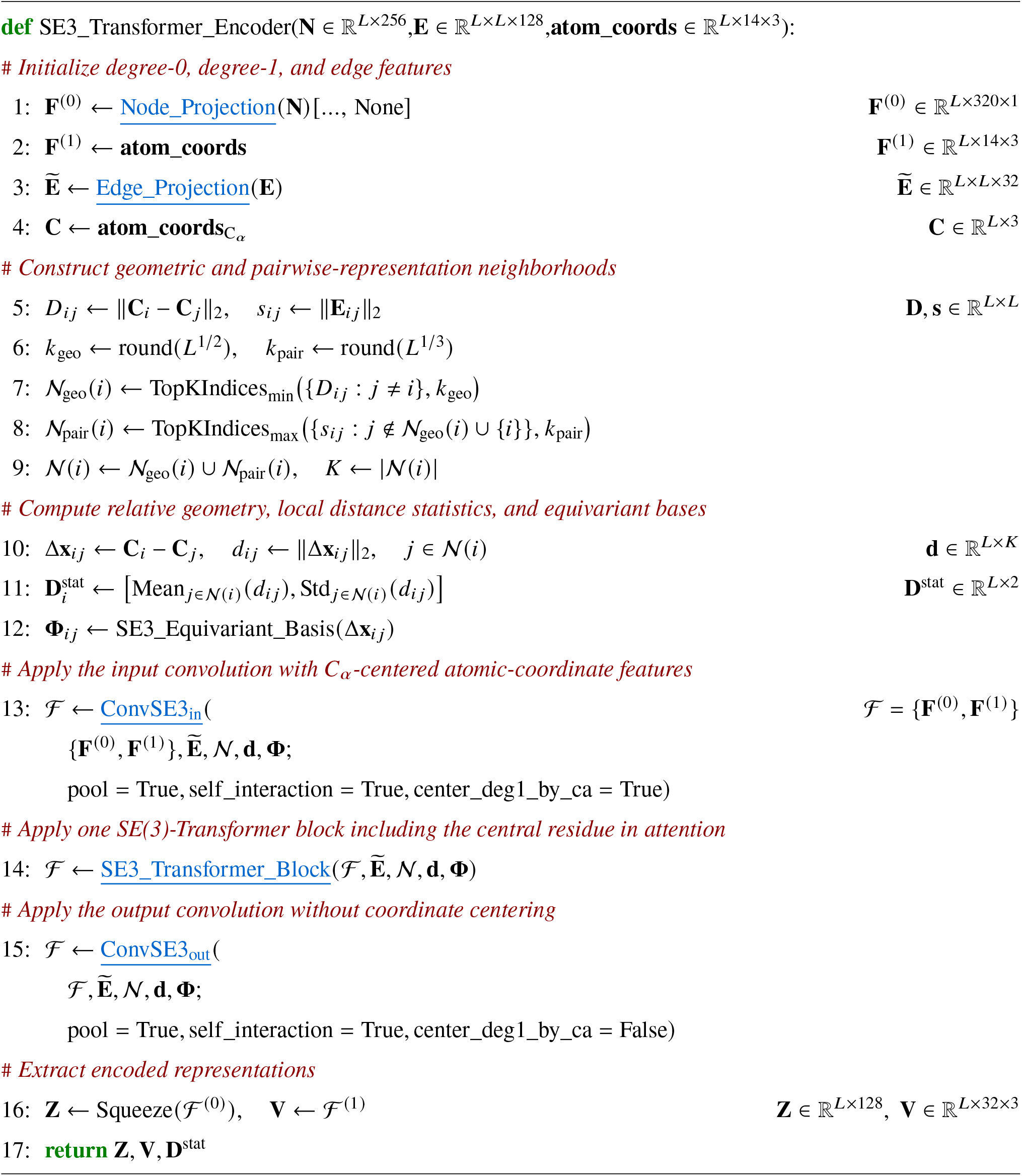

#### S7.4.3 Construction of the single-mutation score tensor and the epistasis representation tensor

Supplementary Algorithm S6 defines the mutation landscape head, which coordinates the single-mutation score branch and the epistasis representation branch detailed in Supplementary Algorithm S7 and Supplementary Algorithm S8, respectively. Given **Z, V, D**^stat^, **X**^ESM-2^, and **wt**_**seq**, the two branches construct the single-mutation score tensor **S** ∈ ℝ^*L*×20^ and the epistasis representation tensor

##### Algorithm S6

Mutation landscape head

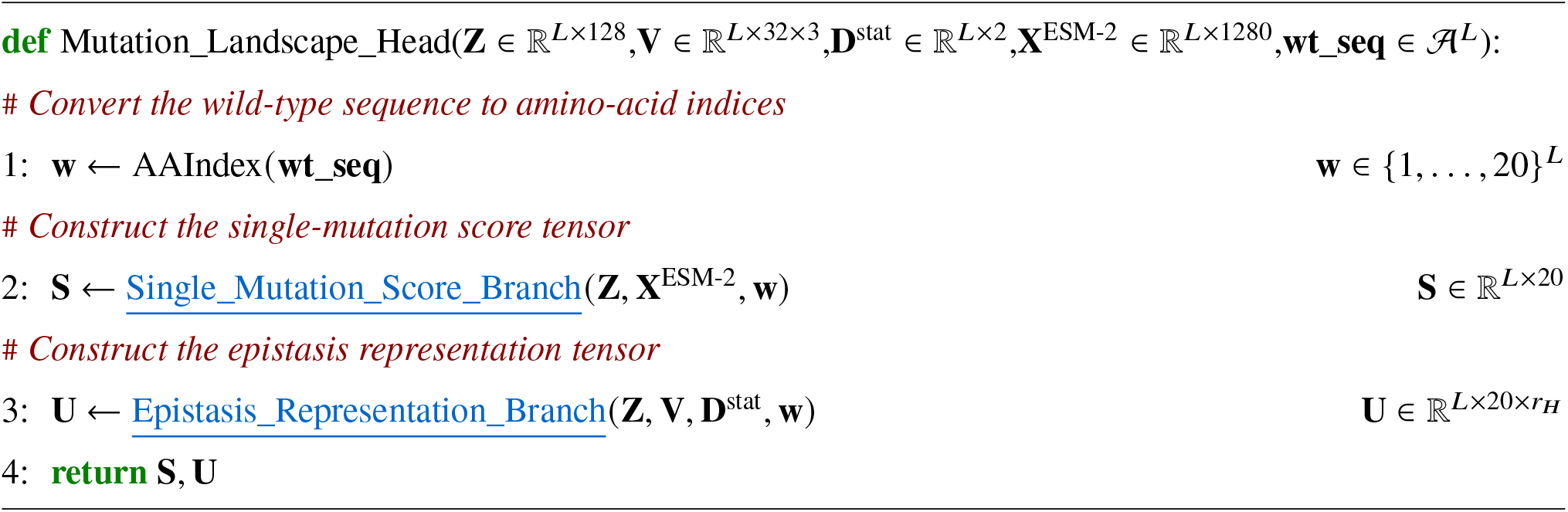

##### Algorithm S7

Single-mutation score branch

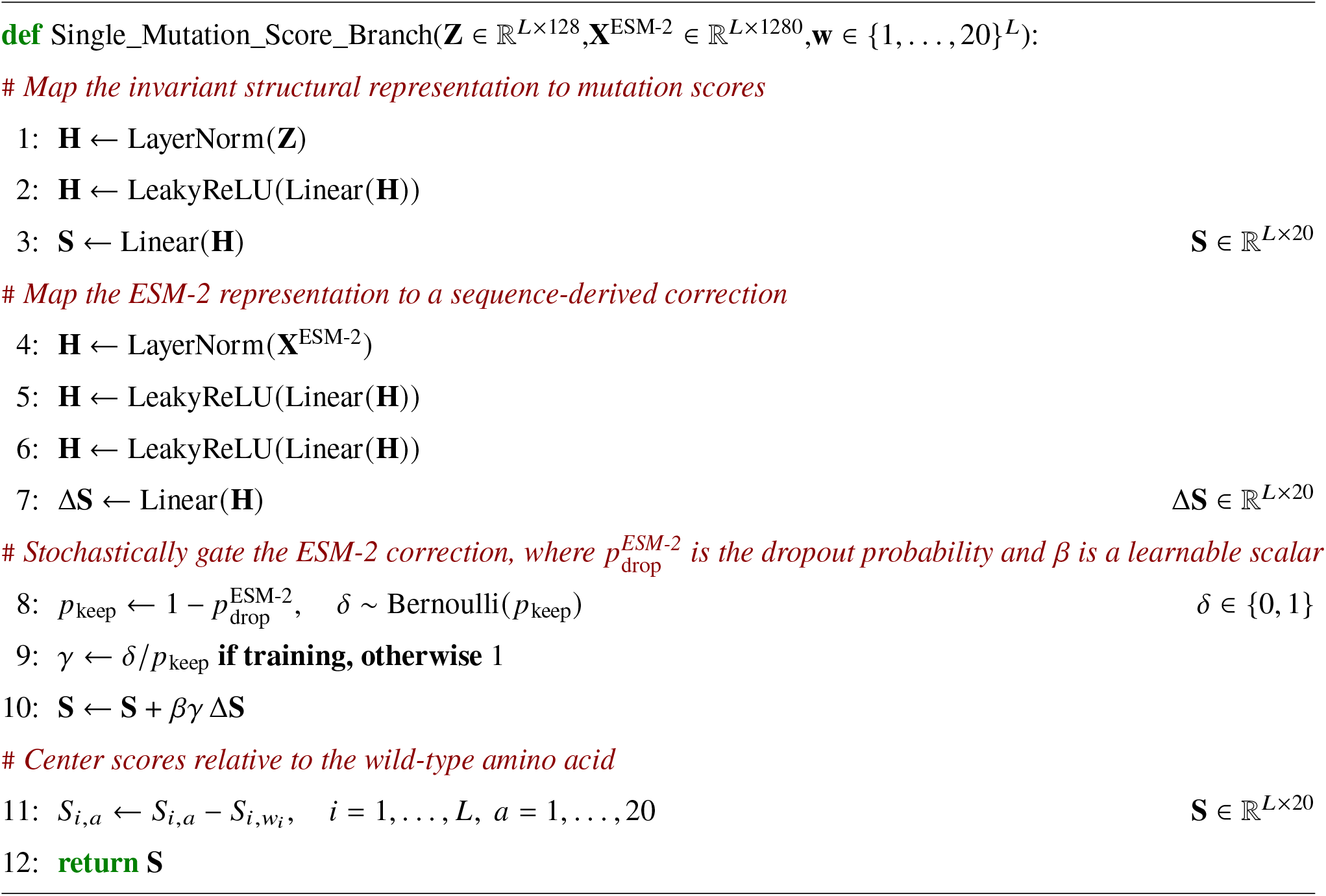

##### Algorithm S8

Epistasis representation branch

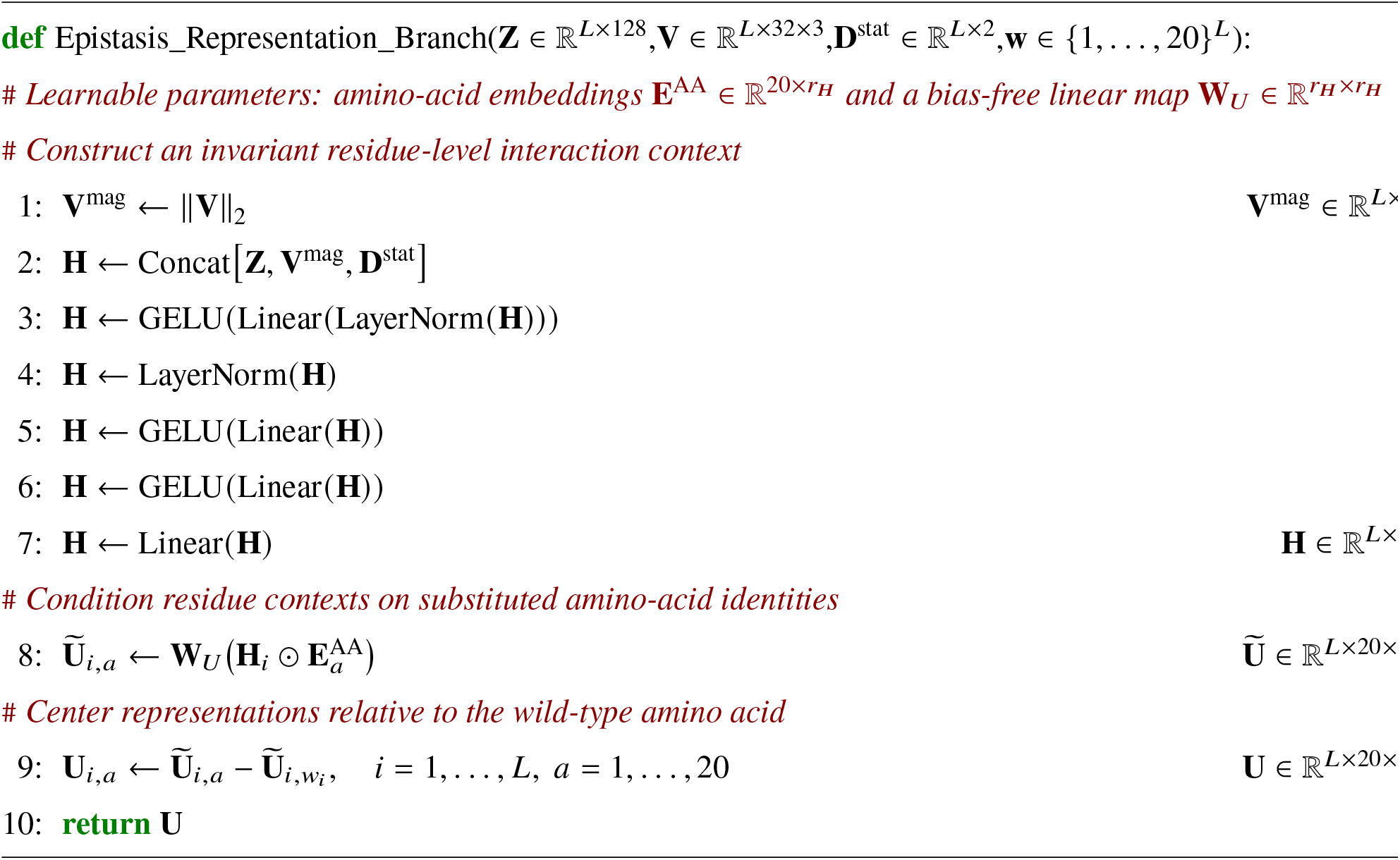

#### S7.4.4 Assembly and prediction of arbitrary-order mutants

Given the precomputed single-mutation score tensor **S** and epistasis representation tensor **U**, the assembly module predicts arbitrary-order mutants without re-encoding their sequences or structures. For notational simplicity, consider an individual mutant 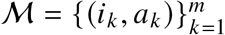, where the mutation-specific score 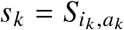 and epistasis representation 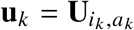 are directly indexed from **S** and **U** for each substitution. The mutant fitness is predicted as

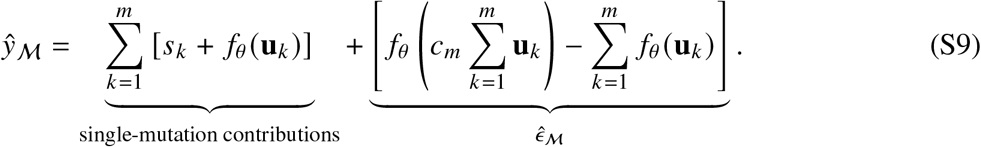

The first term represents the sum of the individual mutation contributions, whereas the second term defines the total epistatic contribution 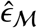. Because the assembly depends only on summation over substitutions, both *ŷ*_M_ and 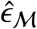 are invariant to mutation ordering. Supplementary Algorithm S9 summarizes the corresponding indexing and assembly rule for an individual mutant. For complete-landscape inference, the same rule is vectorized over batches of mutants in the implementation.

##### Algorithm S9

Assembly and prediction of an arbitrary-order mutant

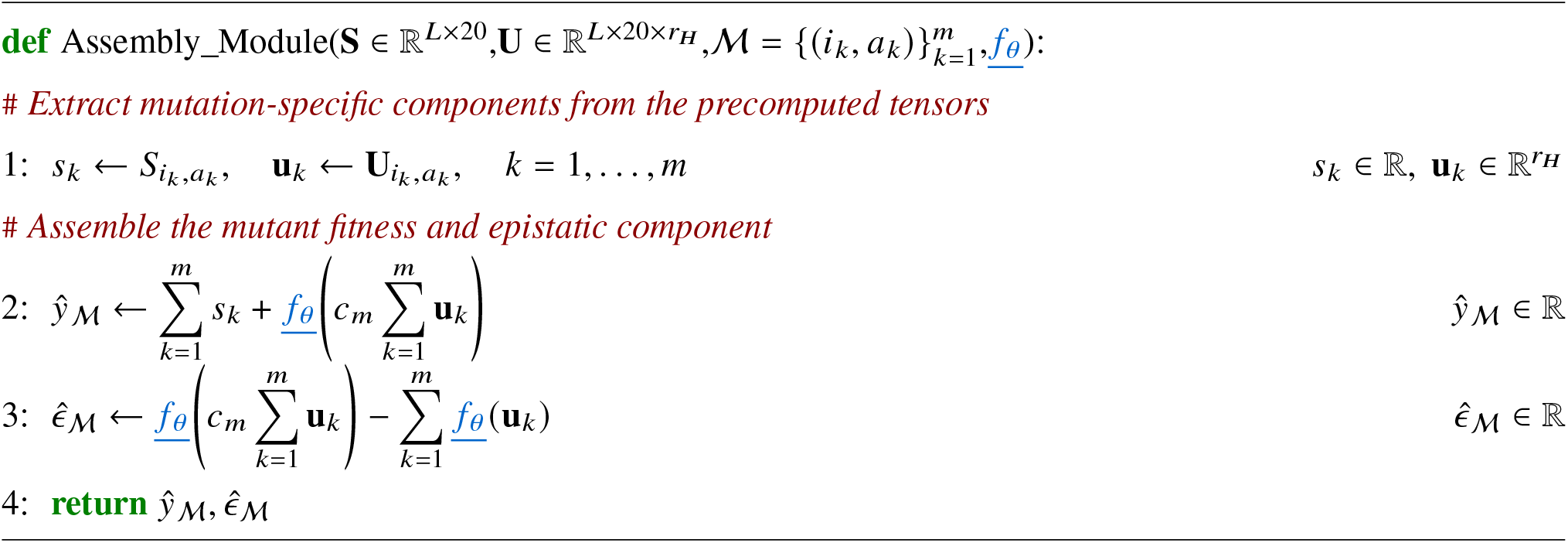

The shared nonlinear interaction function *f*_*θ*_ used in the assembly module maps either an individual epistasis representation or the normalized aggregate of multiple epistasis representations to a scalar response. Supplementary Algorithm S10 details its architecture.

##### Algorithm S10

Learnable interaction function

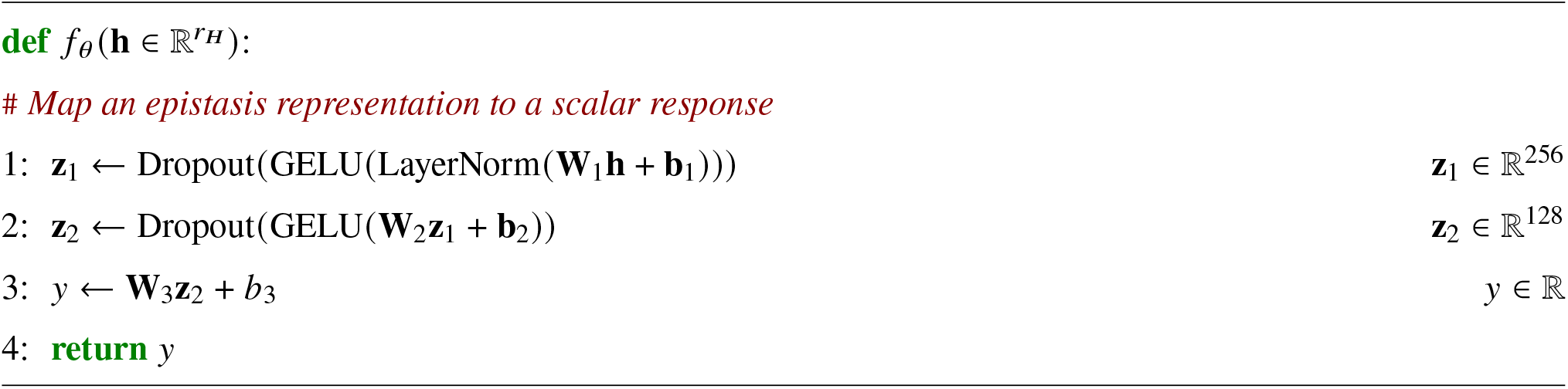

#### S7.4.5 Core operations of the SE(3)-Transformer encoder

Supplementary Algorithm S11 and Supplementary Algorithm S12 describe the input projections applied to the Cerebra residue and pair representations before equivariant processing. The node projection maps the normalized 256-channel residue representation to 320 degree-0 input channels through two linear layers with a LeakyReLU activation and final normalization. The edge projection maps the 128-channel pair representation through a GELU-activated hidden representation to the 32 edge channels used by the equivariant kernels. Supplementary Algorithm S14 and Supplementary Algorithm S13 then summarize the two core equivariant modules used within the encoder. ConvSE3 gathers degree-wise features from the selected neighbors and transforms them using SE(3)-equivariant kernels conditioned on Fourier-encoded inter-residue distances and projected Cerebra pairwise features, together with spherical-harmonic equivariant bases. In ConvSE3_in_, the degree-1 atomic-coordinate features of each neighboring residue are expressed relative to the *C*_*α*_ coordinate of the central residue, with atom-validity masking handled internally and omitted from the pseudocode for clarity. The SE(3)-Transformer block subsequently updates the degree-wise representations through equivariant attention and feed-forward transformations. The remaining mathematical formulations follow the standard SE(3)-Transformer architecture [37].

##### Algorithm S11

Node feature projection

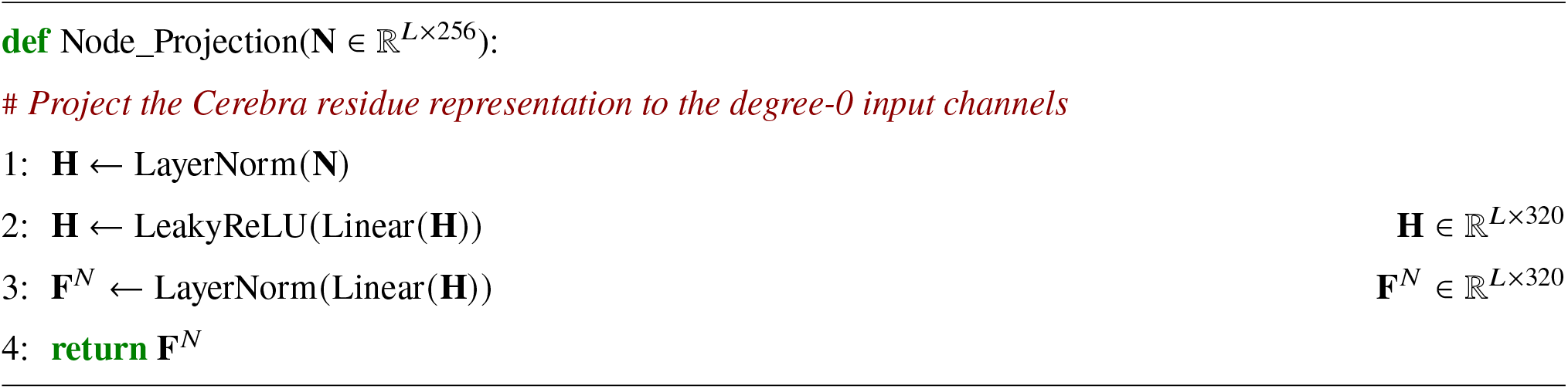

##### Algorithm S12

Edge feature projection

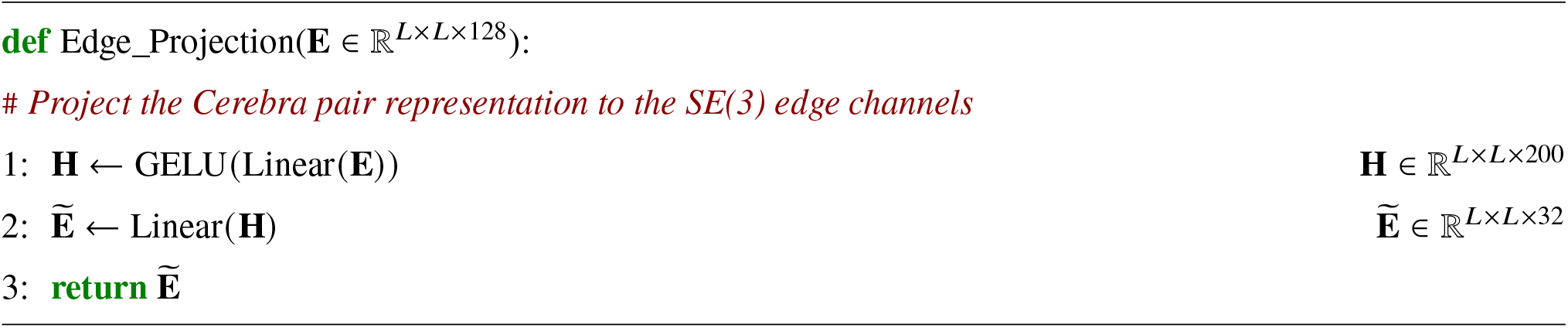

##### Algorithm S13

SE(3)-Transformer block

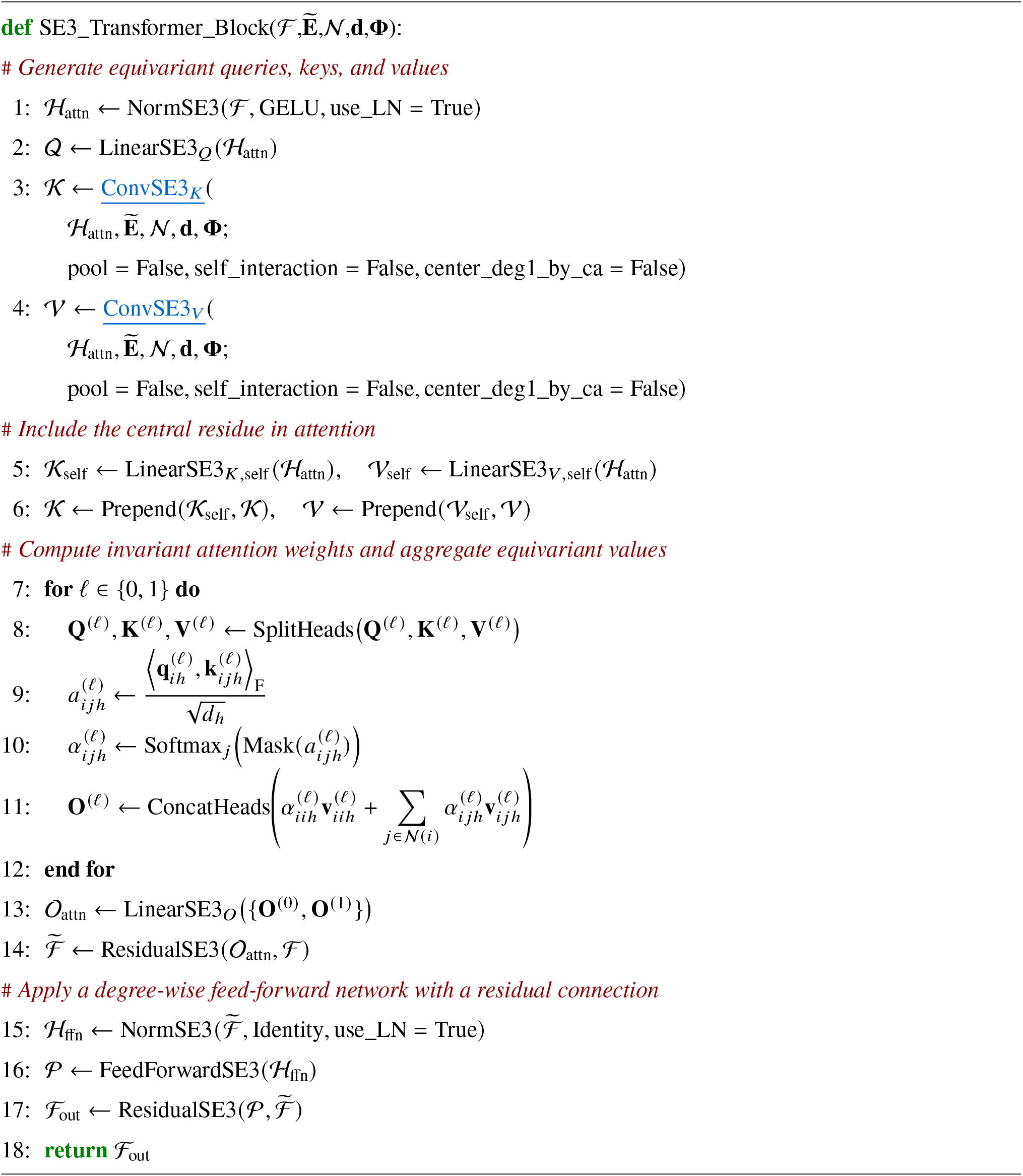

##### Algorithm S14

SE(3)-equivariant convolution

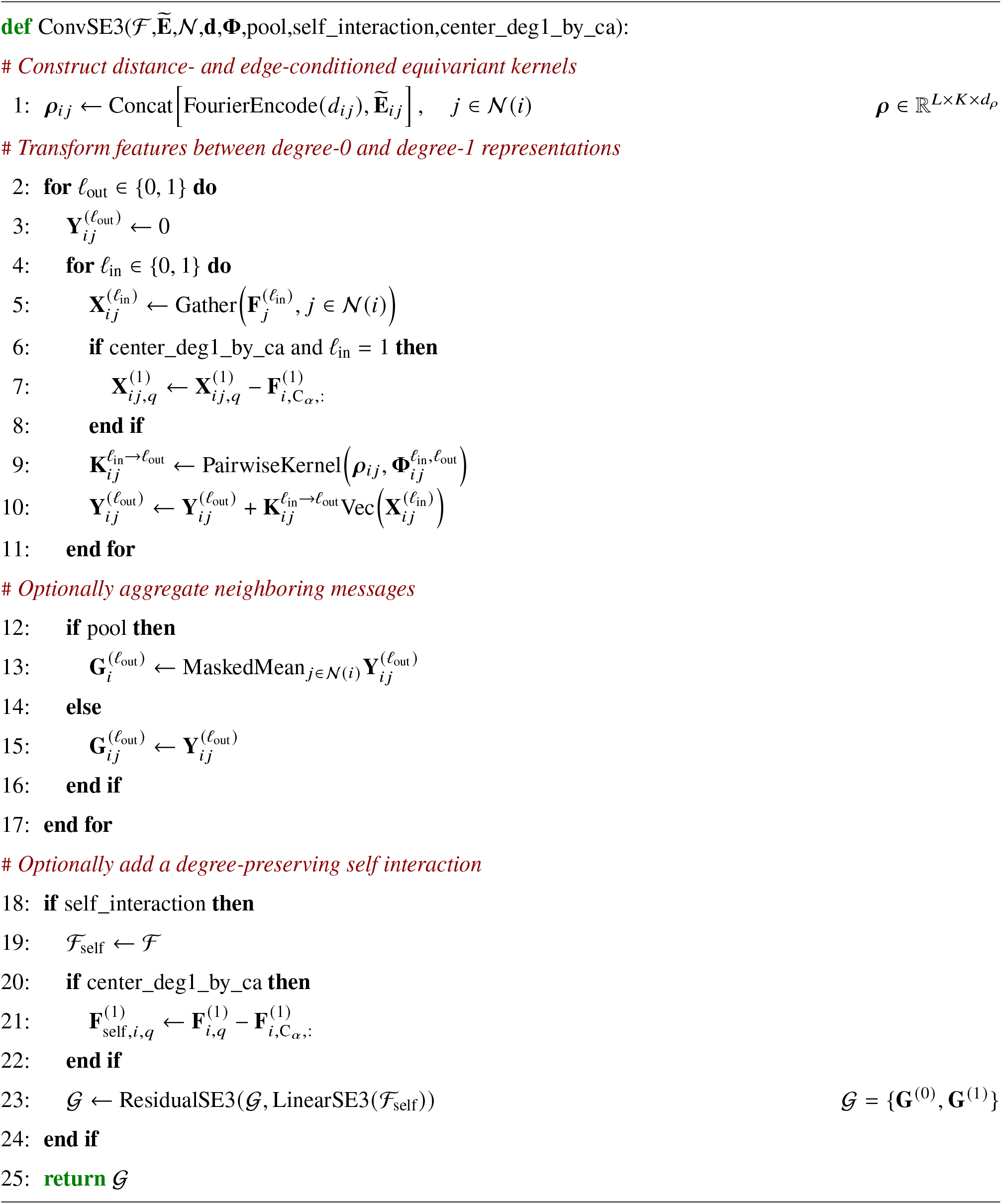

## References

[1] Philip A. Romero and Frances H. Arnold. “Exploring protein fitness landscapes by directed evolution”. In: Nature Reviews Molecular Cell Biology 10.12 (Dec. 2009), pp. 866–876. ISSN: 1471-0080. DOI: 10.1038/nrm2805. URL: http://dx.doi.org/10.1038/nrm2805.

[2] Douglas M Fowler and Stanley Fields. “Deep mutational scanning: a new style of protein science”. In: Nature Methods 11.8 (July 2014), pp. 801–807. ISSN: 1548-7105. DOI: 10.1038/nmeth.3027. URL: http://dx.doi.org/10.1038/nmeth.3027.

[3] Thomas A Hopf et al. “Mutation effects predicted from sequence co-variation”. In: Nature Biotechnology 35.2 (Jan. 2017), pp. 128–135. ISSN: 1546-1696. DOI: 10.1038/nbt.3769. URL: http://dx.doi.org/10.1038/nbt.3769.

[4] Joshua Meier et al. “Language models enable zero-shot prediction of the effects of mutations on protein function”. In: Advances in Neural Information Processing Systems. Ed. by M. Ranzato et al. Vol. 34. Curran Associates, Inc., 2021, pp. 29287–29303. URL: https://proceedings.neurips.cc/paper_files/paper/2021/file/f51338d736f95dd42427296047067694-Paper.pdf.

[5] Pascal Notin et al. “ProteinGym: Large-Scale Benchmarks for Protein Fitness Prediction and Design”. In: Advances in Neural Information Processing Systems 36. NeurIPS 2023. Neural Information Processing Systems Foundation, Inc. (NeurIPS), 2023, pp. 64331–64379. DOI: 10.52202/075280-2810. URL: http://dx.doi.org/10.52202/075280-2810.

[6] Nicholas C Wu et al. “Adaptation in protein fitness landscapes is facilitated by indirect paths”. In: eLife 5 (July 2016). ISSN: 2050-084X. DOI: 10.7554/elife.16965. URL: http://dx.doi.org/10.7554/eLife.16965.

[7] Pascal Notin et al. “ProteinNPT: Improving Protein Property Prediction and Design with Non-Parametric Transformers”. In: Advances in Neural Information Processing Systems 36. NeurIPS 2023. Neural Information Processing Systems Foundation, Inc. (NeurIPS), 2023, pp. 33529–33563. DOI: 10.52202/075280-1458. URL: http://dx.doi.org/10.52202/075280-1458.

[8] Kaiyi Jiang et al. “Rapid in silico directed evolution by a protein language model with EVOLVEpro”. In: Science 387.6732 (Jan. 2025). ISSN: 1095-9203. DOI: 10.1126/science.adr6006. URL: http://dx.doi.org/10.1126/science.adr6006.

[9] Vincent Q. Tran et al. “Rapid directed evolution guided by protein language models and epistatic interactions”. In: Science 392.6798 (May 2026). ISSN: 1095-9203. DOI: 10.1126/science.aea1820. URL: http://dx.doi.org/10.1126/science.aea1820.

[10] Palash Sethi and Juannan Zhou. “An interpretable neural network unveils higher-order epistasis in large protein sequence-function relationships”. In: (Sept. 2024). DOI: 10.1101/2024.09.22.614318. URL: http://dx.doi.org/10.1101/2024.09.22.614318.

[11] Steffanie Paul et al. “Combining Structure and Sequence for Superior Fitness Prediction”. In: GenBio Workshop at NeurIPS. 2023.

[12] Lasse M. Blaabjerg et al. “SSEmb: A joint embedding of protein sequence and structure enables robust variant effect predictions”. In: Nature Communications 15.1 (Nov. 2024). ISSN: 2041-1723. DOI: 10.1038/s41467-024-53982-z. URL: http://dx.doi.org/10.1038/s41467-024-53982-z.

[13] Peter Groth et al. “Kermut: Composite kernel regression for protein variant effects”. In: Advances in Neural Information Processing Systems 37. NeurIPS 2024. Neural Information Processing Systems Foundation, Inc. (NeurIPS), 2024, pp. 29514–29565. DOI: 10.52202/079017-0929. URL: http://dx.doi.org/10.52202/079017-0929.

[14] Yinghui Chen et al. “An end-to-end framework for the prediction of protein structure and fitness from single sequence”. In: Nature Communications 15.1 (Aug. 2024). ISSN: 2041-1723. DOI: 10.1038/s41467-024-51776-x. URL: http://dx.doi.org/10.1038/s41467-024-51776-x.

[15] John Jumper et al. “Highly accurate protein structure prediction with AlphaFold”. In: Nature 596.7873 (July 2021), pp. 583–589. ISSN: 1476-4687. DOI: 10.1038/s41586-021-03819-2. URL: http://dx.doi.org/10.1038/s41586-021-03819-2.

[16] Zeming Lin et al. “Evolutionary-scale prediction of atomic-level protein structure with a language model”. In: Science 379.6637 (Mar. 2023), pp. 1123–1130. ISSN: 1095-9203. DOI: 10.1126/science.ade2574. URL: http://dx.doi.org/10.1126/science.ade2574.

[17] Ruidong Wu et al. “High-resolution de novo structure prediction from primary sequence”. In: (July 2022). DOI: 10.1101/2022.07.21.500999. URL: http://dx.doi.org/10.1101/2022.07.21.500999.

[18] Jian Hu, Weizhe Wang, and Haipeng Gong. “Cerebra: a computationally efficient framework for accurate protein structure prediction”. In: (Feb. 2024). DOI: 10.1101/2024.02.02.578551. URL: http://dx.doi.org/10.1101/2024.02.02.578551.

[19] Salvatore Candido et al. “Language Modeling Materializes a World Model of Protein Biology”. In: bioRxiv (2026). DOI: 10.64898/2026.06.03.729735. URL: https://www.biorxiv.org/content/10.64898/2026.06.03.729735.

[20] Thomas Hayes et al. “Simulating 500 million years of evolution with a language model”. In: Science 387.6736 (Feb. 2025), pp. 850–858. ISSN: 1095-9203. DOI: 10.1126/science.ads0018. URL: http://dx.doi.org/10.1126/science.ads0018.

[21] Zhenda Xie et al. “mHC: Manifold-Constrained Hyper-Connections”. In: (2025). DOI: 10.48550/ARXIV.2512.24880. URL: https://arxiv.org/abs/2512.24880.

[22] Jürgen Haas et al. “Continuous Automated Model EvaluatiOn (CAMEO) complementing the critical assessment of structure prediction in CASP12”. In: Proteins: Structure, Function, and Bioinformatics 86.S1 (2018), pp. 387–398. DOI: 10.1002/prot.25431.

[23] Kadina E. Johnston et al. “A combinatorially complete epistatic fitness landscape in an enzyme active site”. In: Proceedings of the National Academy of Sciences 121.32 (July 2024). ISSN: 1091-6490. DOI: 10.1073/pnas.2400439121. URL: http://dx.doi.org/10.1073/pnas.2400439121.

[24] C. Anders Olson, Nicholas C. Wu, and Ren Sun. “A Comprehensive Biophysical Description of Pairwise Epistasis throughout an Entire Protein Domain”. In: Current Biology 24.22 (Nov. 2014), pp. 2643–2651. ISSN: 0960-9822. DOI: 10.1016/j.cub.2014.09.072. URL: http://dx.doi.org/10.1016/j.cub.2014.09.072.

[25] Fabian Fuchs et al. “SE(3)-Transformers: 3D Roto-Translation Equivariant Attention Networks”. In: Advances in Neural Information Processing Systems. Ed. by H. Larochelle et al. Vol. 33. Curran Associates, Inc., 2020, pp. 1970–1981. URL: https://proceedings.neurips.cc/paper_files/paper/2020/file/15231a7ce4ba789d13b722cc5c955834-Paper.pdf.

[26] Yang Zhang and Jeffrey Skolnick. “Scoring function for automated assessment of protein structure template quality”. In: Proteins: Structure, Function, and Bioinformatics 57.4 (2004), pp. 702–710. DOI: 10.1002/prot.20264. URL: https://doi.org/10.1002/prot.20264.

[27] Valerio Mariani et al. “lDDT: a local superposition-free score for comparing protein structures and models using distance difference tests”. In: Bioinformatics 29.21 (Aug. 2013), pp. 2722–2728. DOI: 10.1093/bioinformatics/btt473. URL: https://doi.org/10.1093/bioinformatics/btt473.

## Supplementary References

[1] Mihaly Varadi et al. “AlphaFold Protein Structure Database: massively expanding the structural coverage of protein-sequence space with high-accuracy models”. In: Nucleic Acids Research 50.D1 (2022), pp. D439–D444. DOI: 10.1093/nar/gkab1061.

[2] Inigo Barrio-Hernandez et al. “Clustering predicted structures at the scale of the known protein universe”. In: Nature 622.7983 (2023), pp. 637–645. DOI: 10.1038/s41586-023-06510-w.

[3] Wolfgang Kabsch and Christian Sander. “Dictionary of protein secondary structure: Pattern recognition of hydrogen-bonded and geometrical features”. In: Biopolymers 22.12 (1983), pp. 2577–2637. DOI: 10.1002/bip.360221211.

[4] Kotaro Tsuboyama et al. “Mega-scale experimental analysis of protein folding stability in biology and design”. In: Nature 620.7973 (July 2023), pp. 434–444. ISSN: 1476-4687. DOI: 10.1038/s41586-023-06328-6. URL: http://dx.doi.org/10.1038/s41586-023-06328-6.

[5] Andre J. Faure et al. “The genetic architecture of protein stability”. In: Nature 634.8035 (Sept. 2024), pp. 995–1003. ISSN: 1476-4687. DOI: 10.1038/s41586-024-07966-0. URL: http://dx.doi.org/10.1038/s41586-024-07966-0.

[6] Thuy-Lan V Lite et al. “Uncovering the basis of protein-protein interaction specificity with a combinatorially complete library”. In: eLife 9 (Oct. 2020). ISSN: 2050-084X. DOI: 10.7554/elife.60924. URL: http://dx.doi.org/10.7554/eLife.60924.

[7] David Ding et al. “Protein design using structure-based residue preferences”. In: Nature Communications 15.1 (Feb. 2024). ISSN: 2041-1723. DOI: 10.1038/s41467-024-45621-4. URL: http://dx.doi.org/10.1038/s41467-024-45621-4.

[8] Kadina E. Johnston et al. “A combinatorially complete epistatic fitness landscape in an enzyme active site”. In: Proceedings of the National Academy of Sciences 121.32 (July 2024). ISSN: 1091-6490. DOI: 10.1073/pnas.2400439121. URL: http://dx.doi.org/10.1073/pnas.2400439121.

[9] Christoph Küng et al. “Deep mutational scanning reveals a de novo disulfide bond and combinatorial mutations for engineering thermostable myoglobin”. In: Protein Science 34.5 (Apr. 2025). ISSN: 1469-896X. DOI: 10.1002/pro.70112. URL: http://dx.doi.org/10.1002/pro.70112.

[10] Allison Judge et al. “Network of epistatic interactions in an enzyme active site revealed by large-scale deep mutational scanning”. In: Proceedings of the National Academy of Sciences 121.12 (Mar. 2024). ISSN: 1091-6490. DOI: 10.1073/pnas.2313513121. URL: http://dx.doi.org/10.1073/pnas.2313513121.

[11] Pascal Notin et al. “ProteinGym: Large-Scale Benchmarks for Protein Fitness Prediction and Design”. In: Advances in Neural Information Processing Systems 36. NeurIPS 2023. Neural Information Processing Systems Foundation, Inc. (NeurIPS), 2023, pp. 64331–64379. DOI: 10.52202/075280-2810. URL: http://dx.doi.org/10.52202/075280-2810.

[12] Ziyu Shi et al. “MutCleaner: Cleaning and Standardizing Biological Mutation Datasets for Variant Effect Prediction”. In: bioRxiv (2026). DOI: 10.64898/2026.09.06.749687. URL: https://doi.org/10.64898/2026.09.06.749687.

[13] Tianyu Mi, Nan Xiao, and Haipeng Gong. “GDFold2: A fast and parallelizable protein folding environment with freely defined objective functions”. In: Protein Science 34.2 (2025), e70041. DOI: 10.1002/pro.70041. URL: https://doi.org/10.1002/pro.70041.

[14] Vincent Q. Tran et al. “Rapid directed evolution guided by protein language models and epistatic interactions”. In: Science 392.6798 (May 2026). ISSN: 1095-9203. DOI: 10.1126/science.aea1820. URL: http://dx.doi.org/10.1126/science.aea1820.

[15] Tianyu Mi et al. “Accelerating Virtual Directed Evolution of Proteins via Reinforcement Learning”. In: (June 2025). DOI: 10.1101/2025.06.25.661516. URL: http://dx.doi.org/10.1101/2025.06.25.661516.

[16] Iván Martín Hernández et al. “Predicting protein stability changes upon mutation using a simple orientational potential”. In: Bioinformatics 39.1 (Jan. 2023). Ed. by Alfonso Valencia. ISSN: 1367-4811. DOI: 10.1093/bioinformatics/btad011. URL: http://dx.doi.org/10.1093/bioinformatics/btad011.

[17] Corrado Pancotti et al. “Predicting protein stability changes upon single-point mutation: a thorough comparison of the available tools on a new dataset”. In: Briefings in Bioinformatics 23.2 (Jan. 2022). ISSN: 1477-4054. DOI: 10.1093/bib/bbab555. URL: http://dx.doi.org/10.1093/bib/bbab555.

[19] Zeming Lin et al. “Evolutionary-scale prediction of atomic-level protein structure with a language model”. In: Science 379.6637 (Mar. 2023), pp. 1123–1130. ISSN: 1095-9203. DOI: 10.1126/science.ade2574. URL: http://dx.doi.org/10.1126/science.ade2574.

[20] Ruidong Wu et al. “High-resolution de novo structure prediction from primary sequence”. In: (July 2022). DOI: 10.1101/2022.07.21.500999. URL: http://dx.doi.org/10.1101/2022.07.21.500999.

[21] Thomas Hayes et al. “Simulating 500 million years of evolution with a language model”. In: Science 387.6736 (Feb. 2025), pp. 850–858. ISSN: 1095-9203. DOI: 10.1126/science.ads0018. URL: http://dx.doi.org/10.1126/science.ads0018.

[22] Yinghui Chen et al. “An end-to-end framework for the prediction of protein structure and fitness from single sequence”. In: Nature Communications 15.1 (Aug. 2024). ISSN: 2041-1723. DOI: 10.1038/s41467-024-51776-x. URL: http://dx.doi.org/10.1038/s41467-024-51776-x.

[23] Joshua Meier et al. “Language models enable zero-shot prediction of the effects of mutations on protein function”. In: Advances in Neural Information Processing Systems. Ed. by M. Ranzato et al. Vol. 34. Curran Associates, Inc., 2021, pp. 29287–29303. URL: https://proceedings.neurips.cc/paper_files/paper/2021/file/f51338d736f95dd42427296047067694-Paper.pdf.

[24] Pascal Notin et al. “Tranception: Protein Fitness Prediction with Autoregressive Transformers and Inference-time Retrieval”. In: Proceedings of the 39th International Conference on Machine Learning. Vol. 162. PMLR, 2022, pp. 16990–17017.

[25] Roshan M. Rao et al. “MSA Transformer”. In: Proceedings of the 38th International Conference on Machine Learning. Vol. 139. PMLR, 2021, pp. 8844–8856.

[26] Peter Groth et al. “Kermut: Composite kernel regression for protein variant effects”. In: Advances in Neural Information Processing Systems 37. NeurIPS 2024. Neural Information Processing Systems Foundation, Inc. (NeurIPS), 2024, pp. 29514–29565. DOI: 10.52202/079017-0929. URL: http://dx.doi.org/10.52202/079017-0929.

[27] Pascal Notin et al. “ProteinNPT: Improving Protein Property Prediction and Design with Non-Parametric Transformers”. In: Advances in Neural Information Processing Systems 36. NeurIPS 2023. Neural Information Processing Systems Foundation, Inc. (NeurIPS), 2023, pp. 33529–33563. DOI: 10.52202/075280-1458. URL: http://dx.doi.org/10.52202/075280-1458.

[28] Yunzhuo Zhou et al. “DDMut: predicting effects of mutations on protein stability using deep learning”. In: Nucleic Acids Research 51.W1 (2023), W122–W128. DOI: 10.1093/nar/gkad472.

[29] James P. Roney, Chenxi Ou, and Sergey Ovchinnikov. “Protein Diffusion Models as Statistical Potentials”. In: bioRxiv (2025). DOI: 10.64898/2025.12.09.693073.

[30] Dmitriy Umerenkov et al. “PROSTATA: a framework for protein stability assessment using transformers”. In: Bioinformatics 39.11 (2023), btad671. DOI: 10.1093/bioinformatics/btad671.

[31] Lasse M. Blaabjerg et al. “Rapid protein stability prediction using deep learning representations”. In: eLife 12 (2023), e82593. DOI: 10.7554/eLife.82593.

[32] Yunxin Xu, D. Liu, and Haipeng Gong. “Improving the prediction of protein stability changes upon mutations by geometric learning and a pre-training strategy”. In: Nature Computational Science 4 (2024), pp. 840–850. DOI: 10.1038/s43588-024-00716-2.

[33] Jeffrey Ouyang-Zhang et al. “Predicting a Protein’s Stability under a Million Mutations”. In: Advances in Neural Information Processing Systems. Vol. 36. 2023, pp. 76229–76247. DOI: 10.52202/075280-3332.

[34] Henry Dieckhaus et al. “Transfer learning to leverage larger datasets for improved prediction of protein stability changes”. In: Proceedings of the National Academy of Sciences 121.6 (2024), e2314853121. DOI: 10.1073/pnas.2314853121.

[35] Jinyuan Sun et al. “Structure-based self-supervised learning enables ultrafast protein stability prediction upon mutation”. In: The Innovation 6.1 (2025), p. 100750. DOI: 10.1016/j.xinn.2024.100750.

[36] Ziang Li and Yunan Luo. “Generalizable and scalable protein stability prediction with rewired protein generative models”. In: Nature Communications 17 (2026), p. 891. DOI: 10.1038/s41467-025-67609-4.

[37] Fabian Fuchs et al. “SE(3)-Transformers: 3D Roto-Translation Equivariant Attention Networks”. In: Advances in Neural Information Processing Systems. Ed. by H. Larochelle et al. Vol. 33. Curran Associates, Inc., 2020, pp. 1970–1981. URL: https://proceedings.neurips.cc/paper_files/paper/2020/file/15231a7ce4ba789d13b722cc5c955834-Paper.pdf.

